# Anatomical input-output streams within mouse orbitofrontal cortex subdivisions

**DOI:** 10.1101/2025.09.25.678540

**Authors:** Anushree Tripathi, Luciano Censoni, Paolo Medini

## Abstract

The orbitofrontal cortex (OFC) supports flexible value representation and decision making in primates, where different OFC subdivisions encode rewarding or aversive stimuli, and OFC dysfunction is implicated in depression. Recent anatomical and functional investigations have been performed in mice, but species differences in reward and value encoding physiology calls for deeper investigation of mouse OFC connectivity to better support translational research. Here, we compared density of retrograde inputs, axonal outputs and inter-subdivision connectivity, focusing on MO (medial-orbital), VO (ventral-orbital), (D)LO ((dorso)lateral-orbital) and AI (agranular insula).

Central subdivisions (LO,VO) received dominant inputs from mediodorsal and submedius thalamic nuclei and projected respectively to medial prefrontal areas and sensory cortices. The striatal output was directed towards dorsal striatum, with reciprocal brainstem dopaminergic innervation.

Input sources were more distributed for MO and AI, with reciprocal outputs to medial prefrontal cortices and amygdala, respectively. Striatal output was mainly to ventral striatum, and AI also received strong serotoninergic innervation.

Cluster analysis revealed that VO/LO, and to lesser extent MO/DLO, shared strong similar input distributions, distinct from AI. Output clustering separated VO targeting sensory areas and AI targeting amygdala. Intra-OFC connectivity suggested information flow preferentially from (D)LO (entry nodes), to VO, MO and finally AI (output node).

Together these data suggest a model that integrates in-series, information on sensory-motor plans (at (D)LO)), motivational state and cue uncertainty (at VO/LO) and current goals (at MO), via AI modulating the amygdaloidal and ventral striatal outputs, thus proposing an experimentally testable circuit to control emotional reactivity and decision-making in mouse OFC.

## INTRODUCTION

In rodents, the Orbitofrontal Cortex (OFC) is situated along the dorsal bank of the rhinal sulcus, where three major subdivisions - the medial (MO), ventral (VO), and lateral (LO) OFC - can be distinguished (Krettek and Price 1977). The VO extends into the rhinal cortex, continuing into the agranular insular cortex (AI). The portion of the AI located rostral to the claustrum is now commonly referred to as the dorsolateral OFC (DLO)(Price 2007). The OFC has been widely implicated in emotional regulation, stimulus salience attribution, reward prediction, and adaptive decision making in both primates and rodents (Furuyashiki and Gallagher 2007, Rudebeck and Rich 2018, Rolls, Cheng et al. 2020). Previous studies have increasingly demonstrated both anatomical and functional heterogeneity of different subdivisions within the rodent OFC, particularly in rats (Barreiros, Ishii et al. 2021), which served as the experimental model in most original studies. Of relevance, medial and lateral OFC process preferentially rewarding and aversive stimuli in primates (Rolls, Cheng et al. 2020), a functional organization that is disrupted in major depression (Zhang, Rolls et al. 2024).

With the advancement of genetic manipulation and viral transfection techniques, the use of mice as a model system has steadily expanded. Several physiological studies have already demonstrated distinct functional heterogeneity across mouse OFC subdivisions. For example, OFC projections emanating from specific OFC subdivisions have been shown to play critical roles in updating reward salience (Gourley, Zimmermann et al. 2016) and integrating stimulus and contextual cues with inferred outcomes to guide adaptive behavior (Ward, Winiger et al. 2015, Cazares, Schreiner et al. 2022). These functional distinctions are potentially correlated to diverse underlying cytoarchitectural patterns. However, in contrast to rats, where input–output relationships of OFC subdivisions have been systematically investigated (Izquierdo 2017, Barreiros, Panayi et al. 2021), current knowledge in mice remains incomplete. While individual studies have examined inputs (Zimmermann, Yamin et al. 2017, Yang, Yang et al. 2025) and outputs (Atlas. 2011, Oh, Harris et al. 2014) of specific OFC subdivisions, a comprehensive framework of the full input–output matrix including the intra-area connectivity is still lacking.

In the present study, we aim to propose such a framework by systematically characterizing the anatomical organization of input–output circuits across OFC subdivisions in mice and relating it to the interconnectivity among subdivisions. Using high-resolution anatomical tracing of retrograde afferents in conjunction with the extensive OFC output data available from the Allen Brain Atlas (Atlas. 2011, Oh, Harris et al. 2014), we highlight key anatomical disparities, reciprocal connections, and topographical patterns. Our analyses reveal distinct subdivision-specific biases, such as preferential VO projections to sensory cortices, AI projections to the amygdala, and MO projections to the mPFC. Despite strong reciprocal connections, we also identify a putative preferential flow within intra-OFC circuitry from the DLO to LO/ VO to MO, and subsequently to AI. This proposed pathway can integrate inputs from the sensory cortices, brainstem, thalamus and amygdala at different stages of processing, suggesting a role for both intra-OFC convergence and hierarchical information flow in the processes of economic value encoding and decision making.

We propose that this organizational framework provides a mechanistic basis for flexible behavioral outputs, arising from differential computations of sensory cues, contextual information, and motivation, past actions, and interoceptive and emotional states. Furthermore, by delineating this circuitry, our study offers concrete, testable hypotheses for future microcircuit manipulations aimed at understanding the causal basis of OFC-mediated functions.

## MATERIALS AND METHODS

### Ethical Considerations and Animal Use

All experimental procedures were approved by the Northern Sweden Ethical Committee (permit A26-2018). Male C57BL/6J mice (n = 16; age 10–13 weeks; Charles River Laboratories) were housed at the Umeå Center for Comparative Biology under a 12-hour light/dark cycle, with ad libitum access to food and water.

### Anatomical Injections and Tissue Processing

Mice were anesthetized with a combination of 10% ketamine and 5% xylazine (0.1ml/15g body weight, intraperitoneally) and placed in a stereotaxic apparatus (WPI, UK). Anesthesia depth was verified by the absence of a pinch reflex and stable respiratory rate. Body temperature was maintained at 37°C using a rectal probe connected to a heating plate. The incision site was depilated, disinfected with betadine, and locally anesthetized with Marcain (0.02ml of 2.5mg/ml, subcutaneously).

Iontophoretic injections of 1% Fluorogold (FG; Fluorochrome LLC) in 0.1M sodium cacodylate buffer (pH 7.36) were made unilaterally into specific subdivisions of the orbitofrontal cortex (OFC) using glass micropipettes (tip diameter 15–20 μm). Pulsed positive currents of 5μA (7 sec on/7 sec off (Midgard Precision, Stoelting) were applied for 10 minutes. The pipette remained in place for an additional minute before slow retraction. Stereotaxic coordinates for OFC subdivisions, adapted from Paxinos and Franklin mouse brain atlas, are provided in Table 1. Coordinates for all OFC subdivisions were kept within the same antero-posterior range for VO, LO and MO. A graphic representation of the workflow is given in Figure 1 while all injection sites analyzed in the present study are in Figure 2.

**Figure 1:**
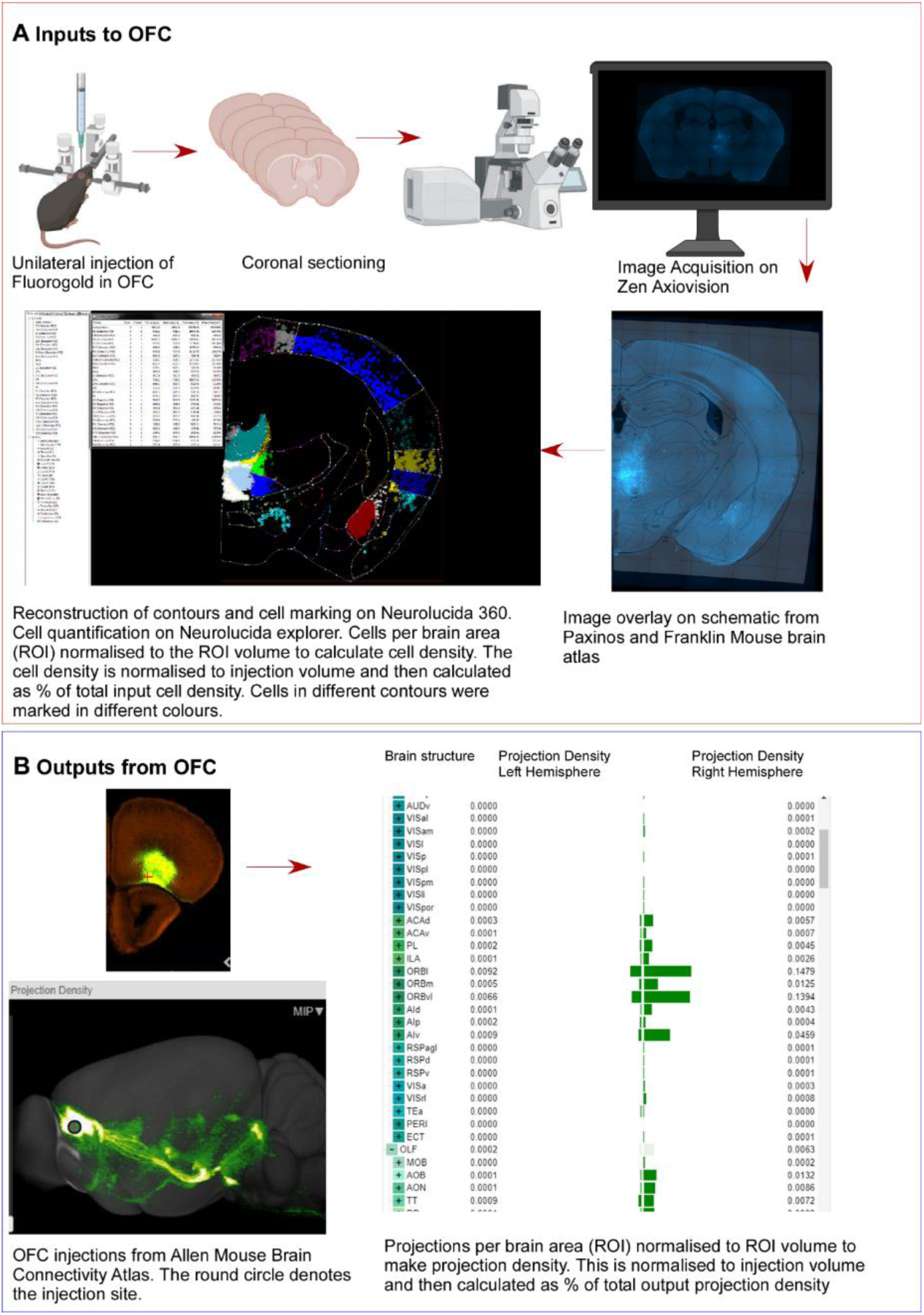
Pipeline for Imaging and Analysis of Retrograde and Anterograde Projections in OFC subdivisions. **A)** *Retrograde Tracing to Map Inputs to OFC:* To investigate afferent inputs to specific orbitofrontal cortex (OFC) subdivisions, the retrograde tracer Fluorogold was unilaterally injected into defined OFC subdivisions. Following transcardial perfusion, brains were coronally sectioned and imaged at 10X magnification. These sections were aligned to corresponding schematics from the Paxinos and Franklin mouse brain atlas, then imported into Neurolucida 360. Within Neurolucida, brain regions were contoured, and Fluorogold-positive cells were marked. Cell quantification was performed using Neurolucida Explorer, which allowed automated analysis and calculation of percentage input cell density as defined in the corresponding figure. **B)** *Anterograde Tracing to Map Outputs from OFC:* Efferent projections from OFC subdivisions were analyzed using publicly available data from the Allen Mouse Brain Connectivity Atlas. For each OFC subdivision, tracer injection experiments were selected (e.g., lateral OFC, http://connectivity.brain-map.org/projection/experiment/168164972). Representative images include the injection site and maximum intensity projection (MIP) of the tracer signal. From the dataset, projection density values were extracted for various brain structures. These values were used to compute percentage output projection density for the ipsilateral hemisphere, as outlined in the corresponding figure.

**Figure 2.**
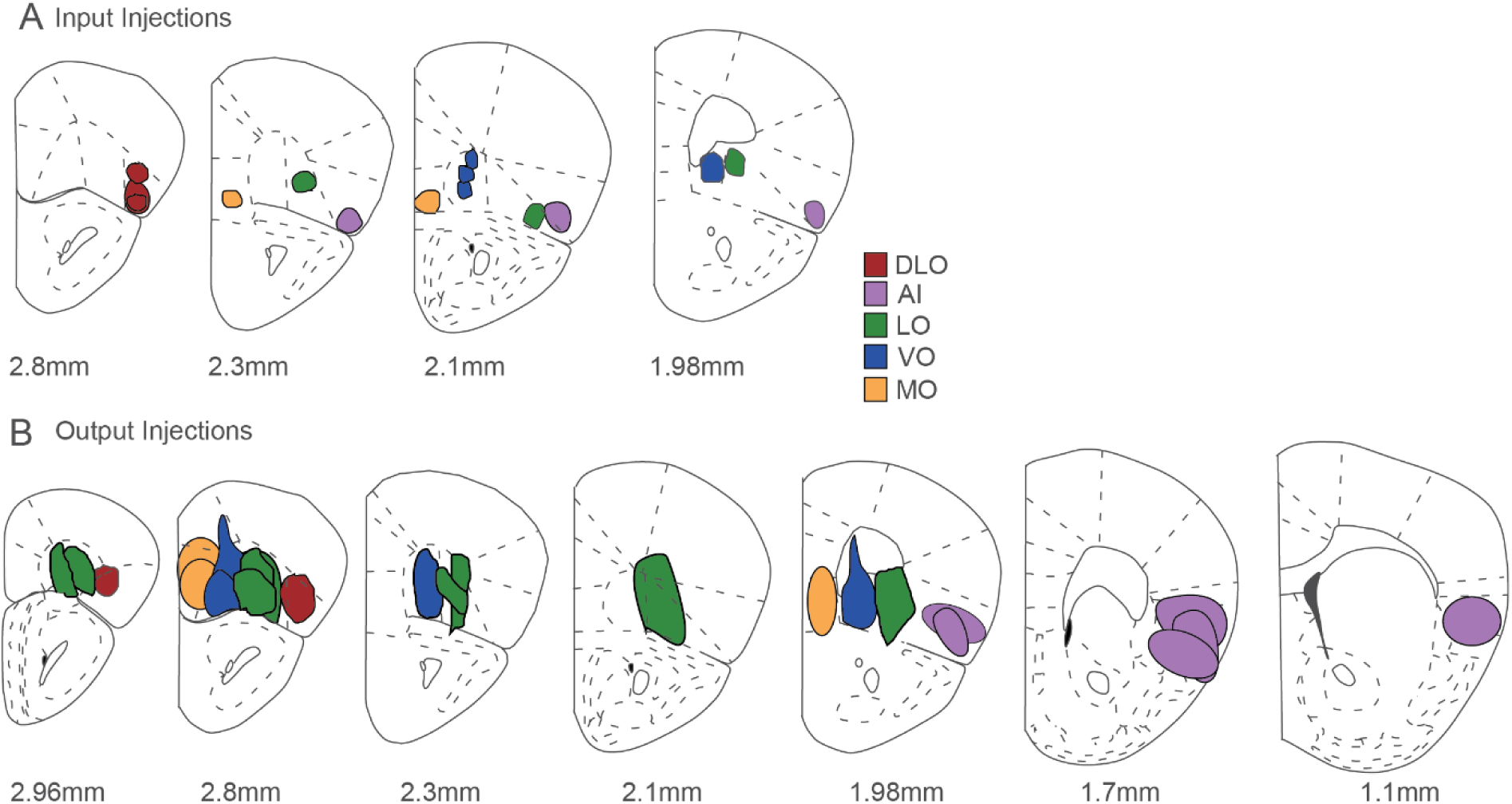
Coronal Section Schematics Depicting the Core of Tracer Deposits in Different OFC Subdivisions. **A)** Polygonal drawings represent individual Fluorogold injection sites in various OFC subdivisions, each annotated with a distinct color. **B)** Injections from the Allen Mouse Brain Connectivity Atlas used in the present study, segregated into different OFC subdivisions. Due to the larger size of these injections, only the densest part of each injection site has been mapped to illustrate the core of the deposit.

**Table 1.**
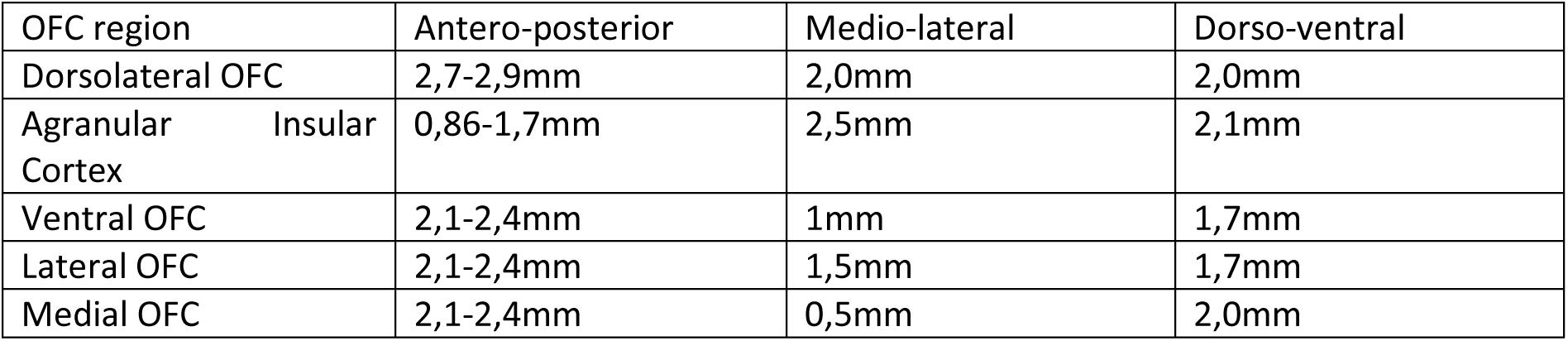
Stereotaxic coordinates for OFC subdivisions, adapted from the mouse brain atlas (Paxinos and Franklin 2001), used in this work to target tracers injections.

| OFC region | Antero-posterior | Medio-lateral | Dorso-ventral |
| --- | --- | --- | --- |
| Dorsolateral OFC | 2,7-2,9mm | 2,0mm | 2,0mm |
| Agranular Insular Cortex | 0,86-1,7mm | 2,5mm | 2,1mm |
| Ventral OFC | 2,1-2,4mm | 1mm | 1,7mm |
| Lateral OFC | 2,1-2,4mm | 1,5mm | 1,7mm |
| Medial OFC | 2,1-2,4mm | 0,5mm | 2,0mm |

Ten days post-surgery, mice were deeply anesthetized (ketamine + xylazine, 0.2 ml/15 g, i.p.) and perfused transcardially with 20 ml of cold saline followed by 50 ml of 4% paraformaldehyde (PFA). Brains were extracted, post-fixed in 4% PFA for 2 hours, then cryoprotected in 30% sucrose at 4°C for 48 hours. Alternate coronal brain sections (60μm) were obtained using a freezing microtome and collected in phosphate-buffered saline (PBS, pH 7.4). Sections were mounted on Permafrost slides, air-dried in the dark, dehydrated, and cover-slipped with Fluoromount-G mounting medium (Invitrogen).

### Image Acquisition

Alternate sections were imaged using a Zeiss Axio Imager M.2 microscope equipped with a wide-band ultraviolet excitation filter (excitation 323 nm, emission 620 nm) for detecting FG. Imaging parameters were kept consistent across all samples. Images were linearly adjusted for brightness and contrast, and superimposed onto corresponding schematics from the Paxinos and Franklin atlas (2001) using Adobe Photoshop. Anatomical alignment was achieved using easily identifiable consistent brain landmarks (e.g., caudate-putamen, hippocampus), as described in Sergejeva, Papp et al. (2015) and further refined following Bjerke, Ovsthus et al. (2023). The table below indicates the brain area and nomenclature used in this work.

**Table 2.**
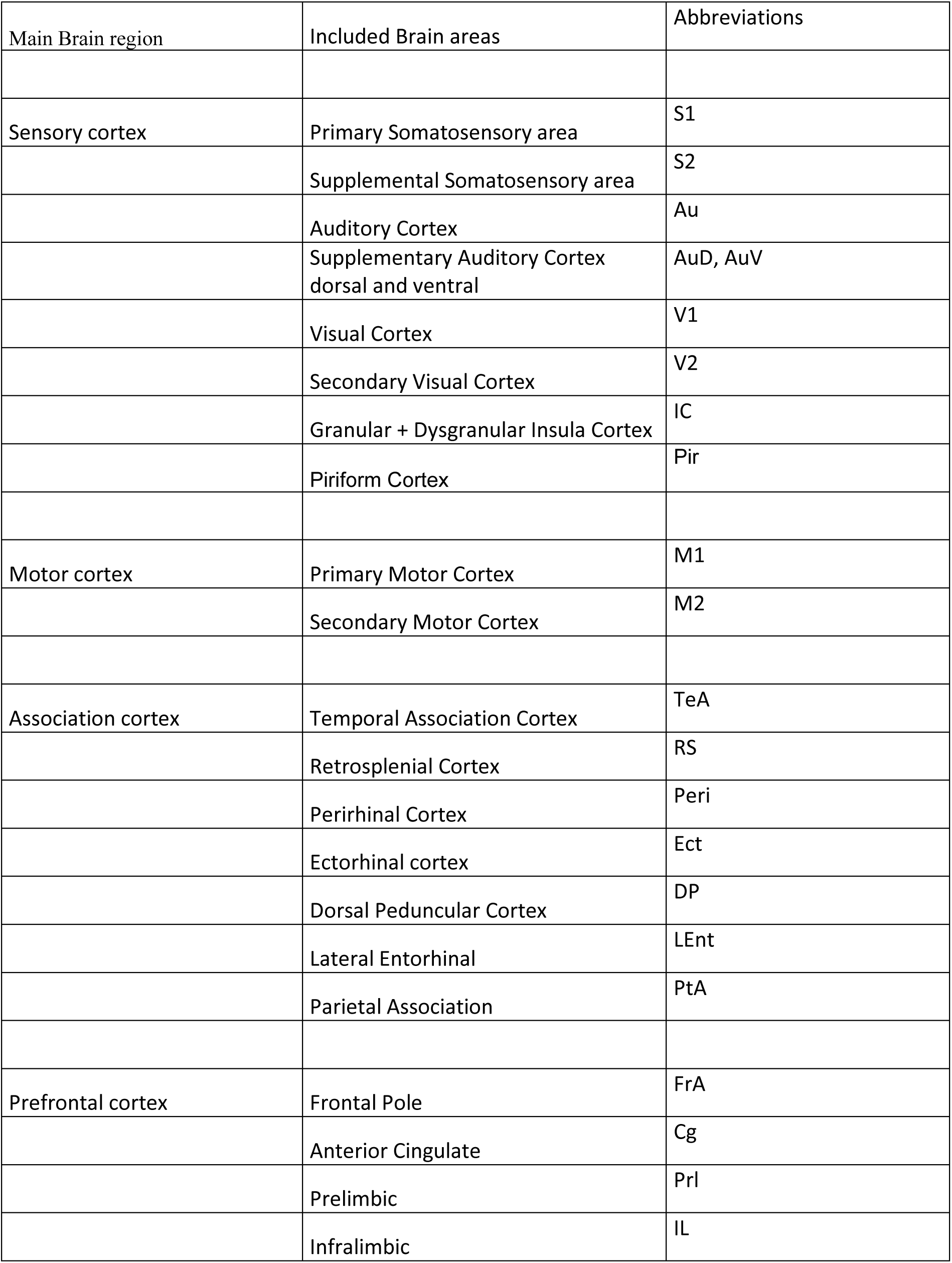
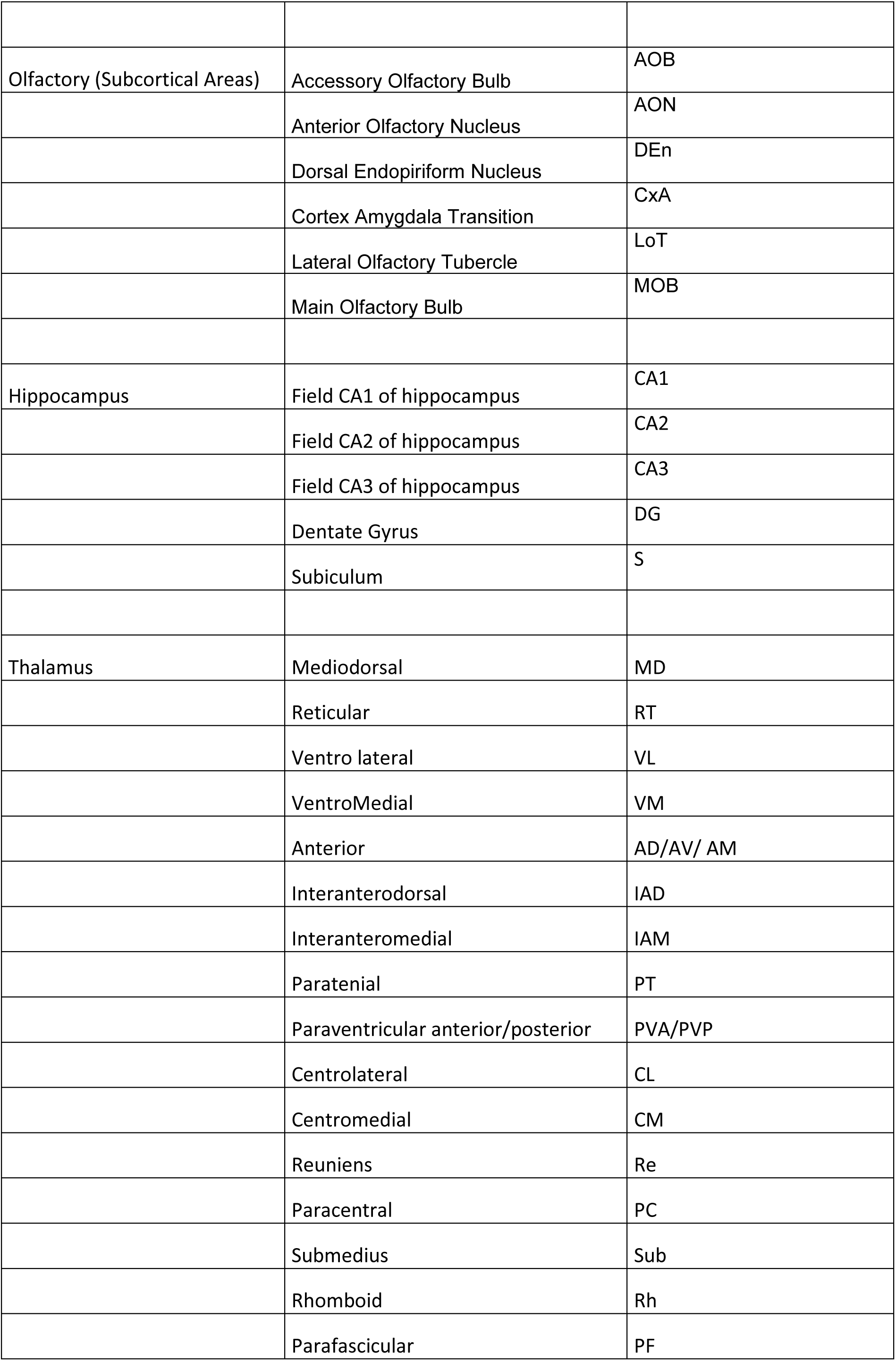
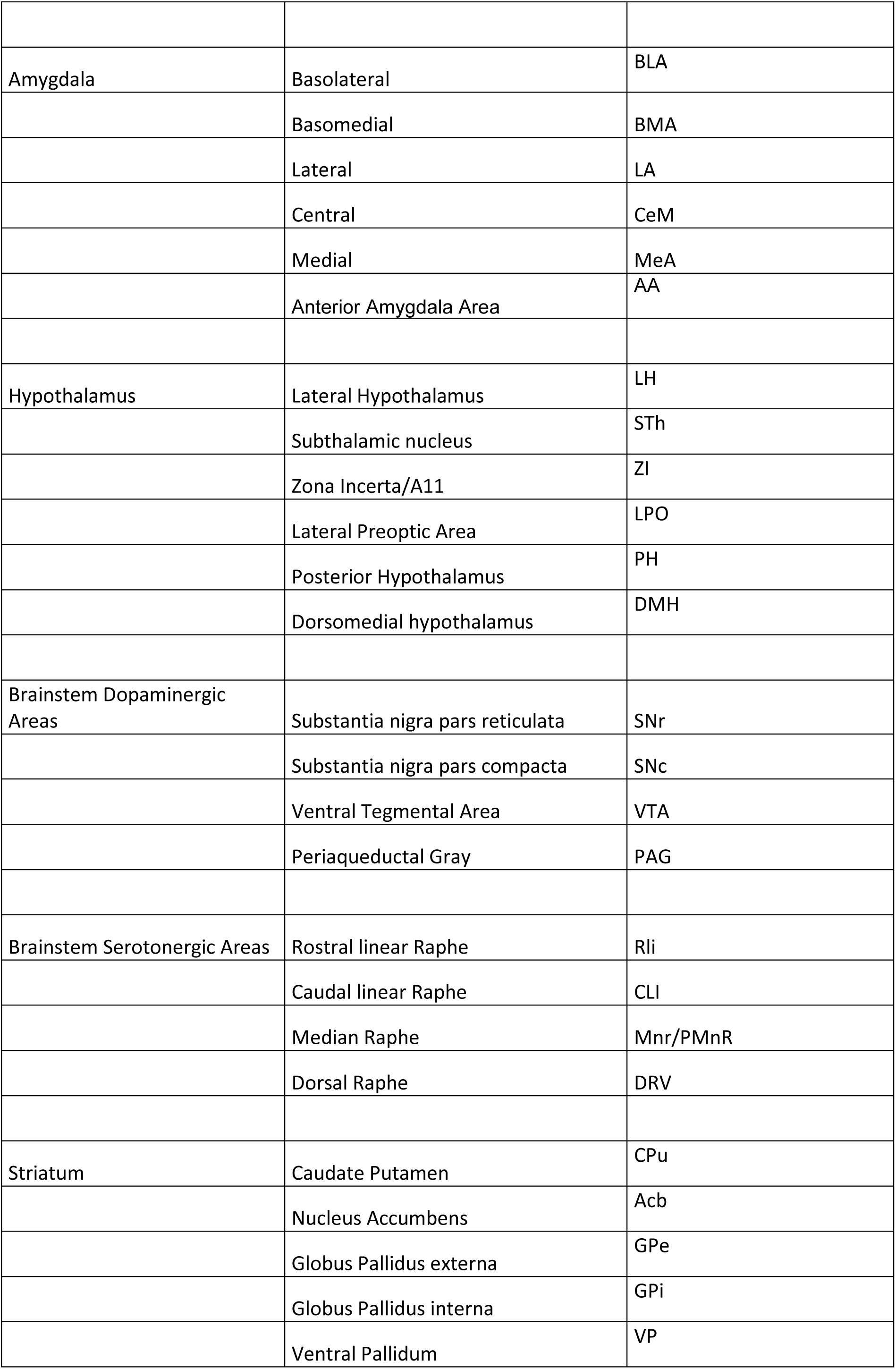
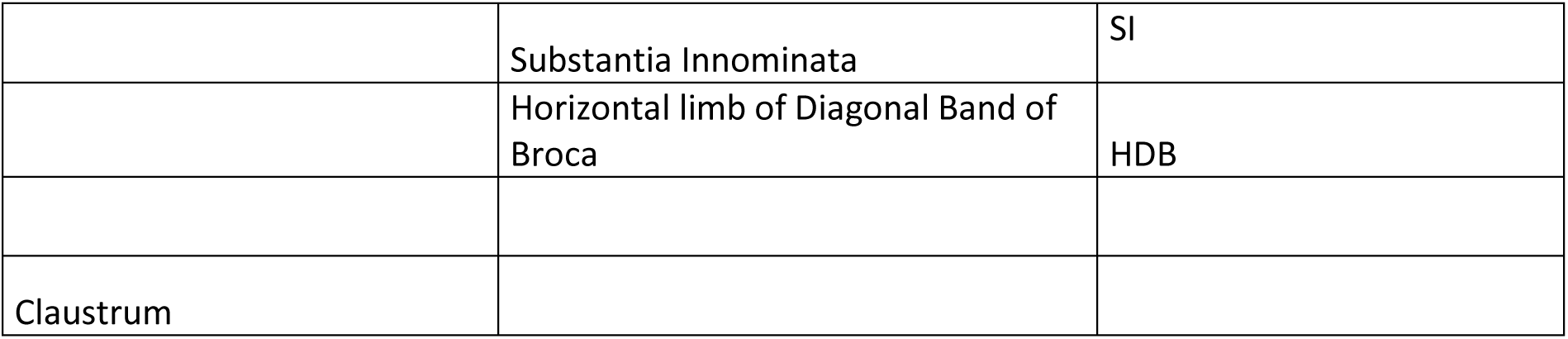
Brain areas nomenclature, abbreviations and groupings as used in this work.

### Image Analysis and Quantification

Superimposed images were first visually inspected for FG-positive cells across brain regions. Cell countings were conducted using Neurolucida 360 and Neurolucida Explorer (MBF Bioscience; workflow in Figure 1). Regions of interest (ROIs) were outlined with color-coded contours, and FG-positive cells within each ROI were marked in distinct colors using the automated proprietary cell detection algorithm of Neurolucida 360 (Hooks et al., 2018). Parameters for cell detection included minimum and maximum cell size and intensity thresholds, optimized automatically using Microbrightfield Neurolucida’s inbuilt proprietary machine learning algorithm. All detected cells were additionally visually verified to eliminate false positives.

Quantitative data, including cell counts and ROI volumes, were extracted with Neurolucida Explorer. Cell density was computed as the number of labeled cells per ROI volume and normalized to the injection volume. To avoid artifacts from tracer spread, data from the injection site were excluded in this analysis. Intra-OFC projections were excluded from the analyses of retrograde input from outside OFC. Final values were expressed as percentage input density, averaged across all animals for each OFC subdivision. Figure 2A shows an overlay of the input tracers injections included in this work.

### Allen Brain Atlas Comparison

Anterograde projection data were obtained from the Allen Mouse Brain Connectivity Atlas (Oh, Harris et al. 2014). Only injections with ≥60% of their volume confined to a specific OFC subdivision (as calculated by the Allen Atlas) were included. This volume threshold was chosen in order to include a reasonable number of injections for analysis while maintaining acceptable confinement of the injection volume within one OFC subdivision. The Allen Brain Atlas does not designate the DLO as a distinct anatomical region. However, as mentioned above, the dorsal portion of the AI (AId), extending rostrally toward the claustrum, corresponds to DLO (Price 2007). This reclassification was supported by its distinct cytoarchitectural features and unique thalamocortical connectivity patterns (Ray and Price 1992, Price 2006). Accordingly, injections within this region were considered as targeting the DLO. To ensure anatomical accuracy, injection coordinates were cross-verified with the Paxinos and Franklin mouse brain atlas prior to analysis. Projection density was calculated as the volume of labeled voxels per ROI divided by the ROI volume, and normalized to injection volume. The final output measure, %output density, was computed by dividing the normalized projection density per ROI by the whole brain innervation density ×100. OFC local outputs and injection sites were excluded from this analysis for consistency with analysis of retrograde data. Figure 2B shows an overlay of the output tracer injections considered.

### Statistical Analysis

Group comparisons were performed using one-way ANOVA followed by Tukey’s post hoc multiple comparisons in GraphPad Prism (v6). In case of striatopallidal output connectivity, paired comparisons were made using Student’s T-test. p-values <0.05 were considered to be significant.

### Input/output correlations

Analyses described here and in the subsequent two sections were performed using custom scripts written for Python 3.10.12, including packages Numpy 2.1.0, Matplotlib 3.9.2 and Scipy 1.14.1.

Average input-input (output-output) correlation matrices between subdivisions were calculated by first constructing, for each injected animal, a 1D sequence of “percentage of total input” (“percentage of total output”) as a function of brain area, in fixed order. Subsequently, the sequences corresponding to all animals injected in the same subdivision were first averaged, resulting in one representative input (output) sequence per subdivision. Then, the Pearson correlation coefficients between the representative input (output) sequences were calculated and reported. Because animal variability is accounted for by pre-averaging, main diagonal elements in the average input-input (output-output) correlation matrices are equal to 1 by definition, while off-diagonal elements represent the similarity between input (output) distributions of distinct subdivisions.

Average input-output correlations were calculated following a similar procedure as described above; the sequences corresponding to all animals injected in the same area were first averaged, resulting in one representative input and one representative output sequence per subdivision. Then, the correlation between the representative input and output sequences of the same subdivision was calculated and reported for each subdivision.

A separate variation was also calculated where, for each area, representative input and output sequences were concatenated into a single longer sequence, and correlation coefficients between concatenated sequences were calculated for all pairs of subdivisions and reported as an average input+ output correlation matrix.

Additionally, input-input (output-output) correlations across all animals (without pre-averaging) were also calculated, and reported in the relevant supplementary materials. These were produced following a similar procedure as described above, but, instead of calculating representative sequences per subdivision, for main diagonal elements, correlation coefficients were calculated between the sequences corresponding to each distinct pair of animals injected in the same subdivision. The mean and standard deviation of the resulting distribution of correlation coefficients was reported, together with the number N of distinct animal pairs, for each subdivision, producing an estimate of the between-animal variability in the distributions of inputs (outputs). For off-diagonal elements, instead, the correlation coefficient was calculated between the sequences corresponding to each distinct pair of animals such that each animal in the pair was injected in a different subdivision. The mean and standard deviation of the resulting distribution of correlation coefficients was reported, together with the number N of distinct animal pairs, for each pair of subdivisions, producing an estimate of the similarity between input (output) distributions of distinct subdivisions that also takes into account between-animal variability.

Additionally, input-output correlations across animals were also calculated and reported in supplementary materials. For each subdivision, the correlation coefficient was calculated between the input and output sequences corresponding to each distinct pair of animals such that one animal was injected with the retrograde tracer (to determine input distributions) and the other with the anterograde tracer (to determine output distributions). The mean and standard deviation of the resulting distribution of correlation coefficients was reported, together with the number N of distinct animal pairs, for each subdivision, producing an estimate of the similarity between the input and output distributions of each subdivision that also takes into account between-animal variability.

### Hierarchical clustering

Hierarchical clustering based on input (output) similarity was performed by calculating an average input (output) distance matrix from the corresponding average input (output) correlation matrix, by replacing each element *r* with *(1-r)*. The average input (output) distance matrix was passed to the Scipy *linkage* method using the *‘average’* parameter for inter-cluster distance calculation, which resulted in the largest correlation between original distance matrix and cophenetic distance matrix among the options tested (*‘single’*, *‘complete’*, *‘average’* and *‘weighted’*) in every case, and the resulting linkage matrix was plotted with the Scipy *dendrogram* method using a fixed but arbitrary threshold of 0.15 for cluster partitioning.

Additionally, hierarchical clustering was also performed on the average concatenated input+ output correlation matrix, following the same procedure as above.

### Intra-OFC connectivity analysis

Intra-OFC putative connectivity strength between subdivisions was calculated by first constructing, for each injected animal, a 1D sequence of “percentage of total input from subdivisions” (“percentage of total output to subdivisions”) as a function of OFC subdivision, in fixed order. The sequences corresponding to all animals injected in the same subdivision were averaged, resulting in one representative intra-OFC input (output) sequence per subdivision. Representative input (output) sequences from all subdivisions were collected and reported in matrix form.

Subsequently, an attempt to recover true connection strengths that respected the input and output matrices simultaneously was made, but the resulting linear system had no solution, suggesting an incompatibility between input-based and output-based measurements. Therefore, a graphical representation of the connectivity structure among all subdivisions was constructed based on the input percentages only. The distribution of connection percentages covered more than one order of magnitude. Therefore, to produce an integrated graphical representation, a thresholding procedure was applied where all connections with strength less than or equal to the median observed strength (coinciding with a clear inflection point in the connection strength histogram) were discarded. The surviving connection strengths and directions were shown as a directed graph.

Additionally, an attempt was made to construct a solvable linear system by repeatedly applying perturbations to the output percentages, in order to obtain matrices compatible with the input results and allow the reconstruction of the true connection strengths between subdivisions. Each element in the output matrix was allowed to vary independently between 10% and 200% of its original value, followed by renormalizing based on column sums, then the linear system of equations combining the input and output percentages was written out, and the smallest singular value of the corresponding matrix was minimized. When the smallest singular value reached zero (within machine precision), the corresponding right singular vector contained the solution to the perturbed linear system, i. e., the reconstructed connection strengths, up to a multiplicative constant. This procedure was repeated over 1000 iterations, the resulting matrices were averaged, and a graphical representation was produced following the same thresholding procedure as above. Results are reported in supplementary materials.

## RESULTS

Here, we present the input and output tracing results, separately for each OFC subdivision, analyzed by quantitative morphometry from retrograde and anterograde injections as specified in the Material and Methods section.

### 1. Whole brain input/ output connectivity from Medial OFC

We analyzed two retrograde and three anterograde tracer injections targeting MO (Figure 2, orange). A representative retrograde injection and the schematic of rostrocaudal distribution of FG-labeled cells are shown in Figure 3A,B.

**Figure 3:**
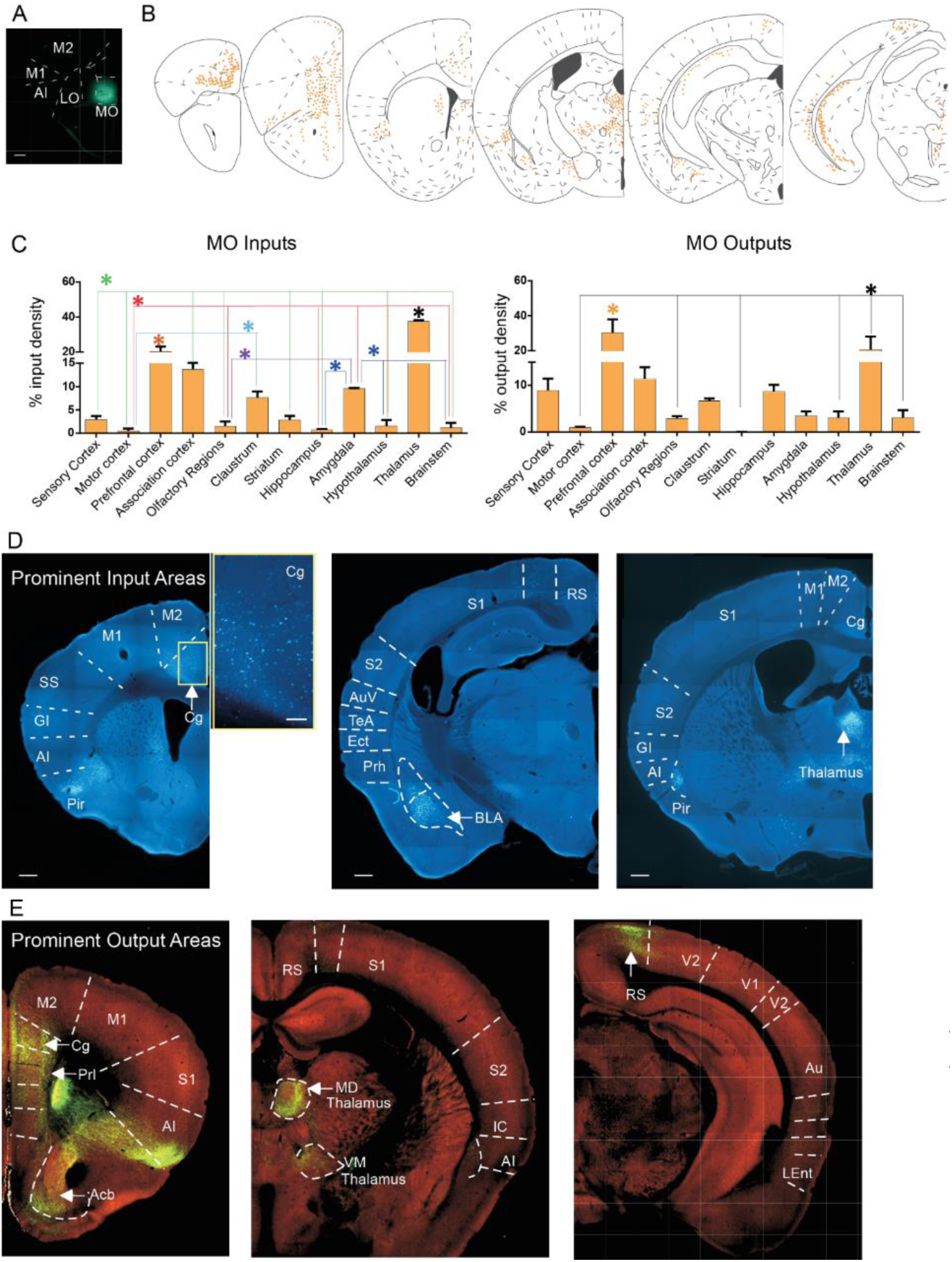
Microphotographs Depicting Inputs to and Outputs from the Medial Orbitofrontal Cortex (MO). **A)** Representative image of a Fluorogold injection site in MO. **B)** Schematic depiction of the rostrocaudal distribution of retrogradely labeled Fluorogold-positive cells across different brain areas. **C)** Histograms showing the percentage of input cell density (left) and output projection density (right) across the main brain areas analyzed in this study. An asterisk above a bar indicates a brain region whose input or output is significantly greater than the others. *p* < 0.05, one-way ANOVA with Tukey’s post hoc test, see also Supplementary 1 for statistics. **D)** Representative images of some prominent input brain areas: anterior cingulate cortex (prefrontal areas; left), basolateral amygdala (BLA; middle), and thalamus (right). The inset shows a magnified image of the retrogradely labelled cells in Cg. **E)** Representative images from the Allen Brain Atlas (http://connectivity.brain-map.org/projection/experiment/272781246) showing some prominent MO output targets. Panels display projections to anterior cingulate and prelimbic cortices (left), mediodorsal (MD) and ventromedial (VM) thalamic nuclei (middle), and retrosplenial association cortex (right). **Scale bars (D):** 200 µm; inset: 100 µm. Arrows indicate the most prominent input/output brain regions.

Retrograde tracing revealed that afferent inputs to MO originated from a broad range of brain regions, without a single dominant source (Figure 3C). Among these, the thalamus (37.38 ± 0.82%) and medial prefrontal cortex (mPFC % input density; 20 ± 2.86%) provided the most substantial projections. These regions were also the primary targets of MO efferents, as demonstrated by anterograde tracing (mPFC % output density: 30.24 ± 7.76%; thalamus: 20.59 ± 7.51%; one-way ANOVA with Tukey post hoc test; see Figure 3C; see also Supplementary 1 for statistics).

Additional substantial input to MO was observed from association cortical areas and the basolateral amygdala (BLA; Figure 3D, middle), which were more prominent compared to projections from for example subcortical olfactory areas or brainstem nuclei (see quantifications in Figure 3C). Similarly, MO projections extended, in addition to mPFC and thalamus, to association cortical areas, hippocampus, and ventral striatum. Interestingly, although quantitative analysis indicated relatively sparse projections to the ventral striatum/nucleus accumbens, qualitative inspection revealed dense and robust axonal labeling in a small confined area of this region (Figure 3E, left).

### 2. Whole brain input/ output connectivity from Ventral OFC

We analyzed four localized retrograde and anterograde tracer injections in VO (Figure 2, blue). A representative retrograde injection and the schematic of rostrocaudal distribution of FG-labeled cells are shown in Figure 4A,B.

**Figure 4:**
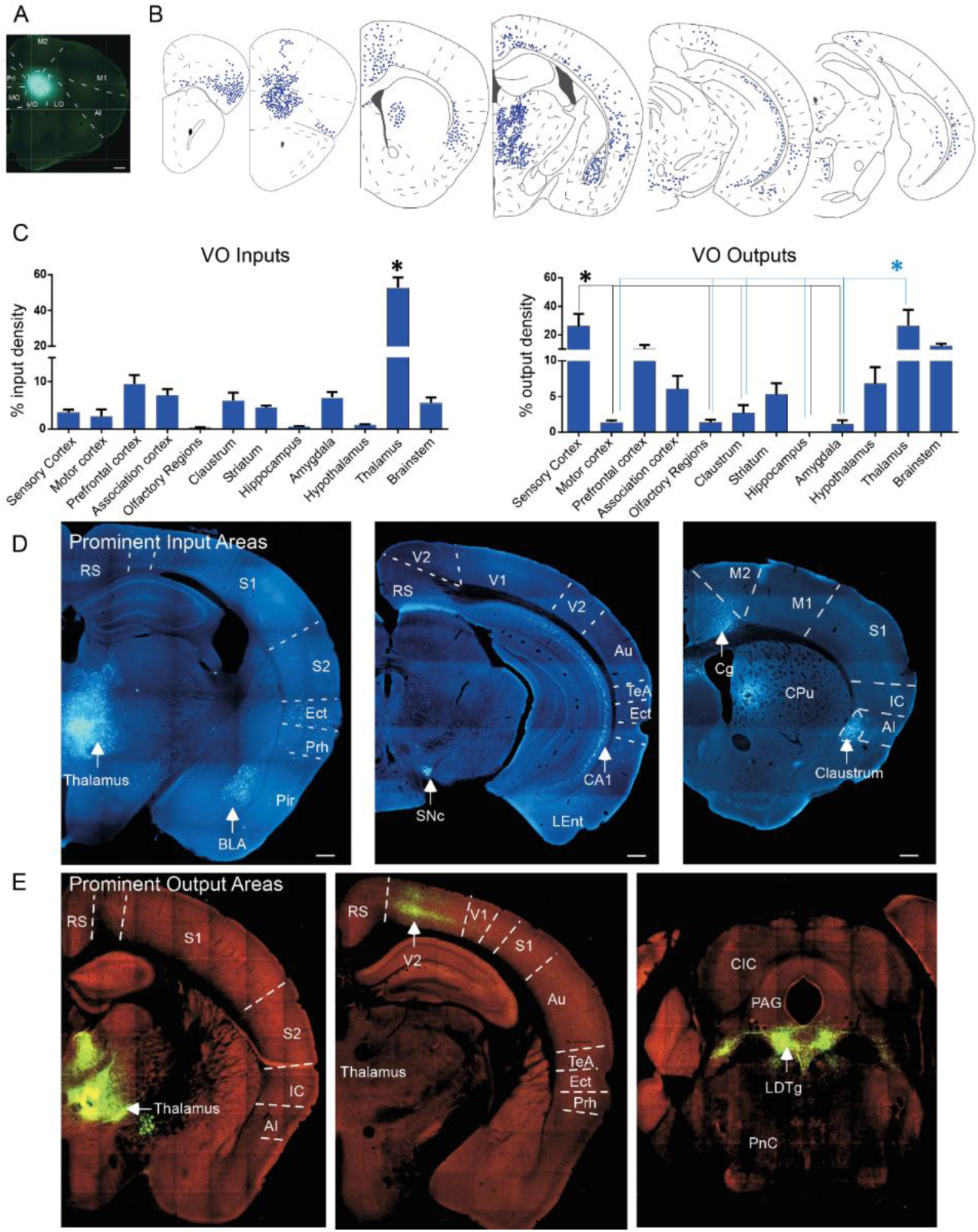
Microphotographs Depicting Inputs to and Outputs from the Ventral Orbitofrontal Cortex (VO). **A)** Representative image of a Fluorogold injection site in VO. **B)** Schematic depiction of the rostrocaudal distribution of retrogradely labeled Fluorogold-positive cells across different brain areas. **C)** Histograms showing the percentage of input cell density (left) and output projection density (right) across the main brain areas analyzed in this study. A single asterisk above a bar indicates a brain region whose input or output is significantly greater than all others. *p* < 0.05, one-way ANOVA with Tukey’s post hoc test, see also Supplementary 1 for statistics **D)** Representative images of some prominent input brain areas: Thalamus (left), substantia nigra pars compacta (SNc) and CA1 region of the hippocampus (middle), and anterior cingulate and claustrum (right)). **E)** Representative images from the Allen Brain Atlas (http://connectivity.brain-map.org/projection/experiment/287769286) showing prominent VO output targets. Panels display projections to the thalamus (left), secondary Visual cortex (V2, middle), and laterodorsal tegmental brainstem nucleus (LDTg, right). **Scale bar (D):** 200 µm. Arrows indicate the most prominent input/output brain regions.

Quantitative analysis of the retrograde tracing demonstrated that the thalamus provided the strongest afferent input to VO (52.60 ± 5.84% input density; Figure 4B), significantly greater than all other brain regions (p < 0.05, one-way ANOVA with Tukey post hoc test). Thalamic projections were reciprocated, as anterograde tracing revealed dense VO efferents to the thalamus (26.42 ± 11.27% output density). In addition, the sensory cortices constituted a major target of VO projections (26.19 ± 8.62%; Figure 4C; see also Supplementary 1 for statistics).

Qualitative examination of retrogradely labeled cells revealed additional strong inputs from the basolateral amygdala (BLA; Figure 4D). Within the hippocampus, FG-positive neurons were observed in the pyramidal cell layer. Retrograde labeling was detected in the claustrum and in the substantia nigra pars compacta (SNc) within the brainstem. On the efferent side, in addition to thalamus, the secondary visual cortex (V2) exhibited dense axonal labeling, while robust VO projections were also observed in several brainstem regions (Figure 4E).

### 3. Whole brain input/ output connectivity from Lateral OFC

We analyzed three retrograde and nine anterograde tracer injections localized within this subdivision (Figure 2, green). A representative retrograde injection and the rostrocaudal distribution of FG-labeled cells are shown in Figure 5A,B.

**Figure 5:**
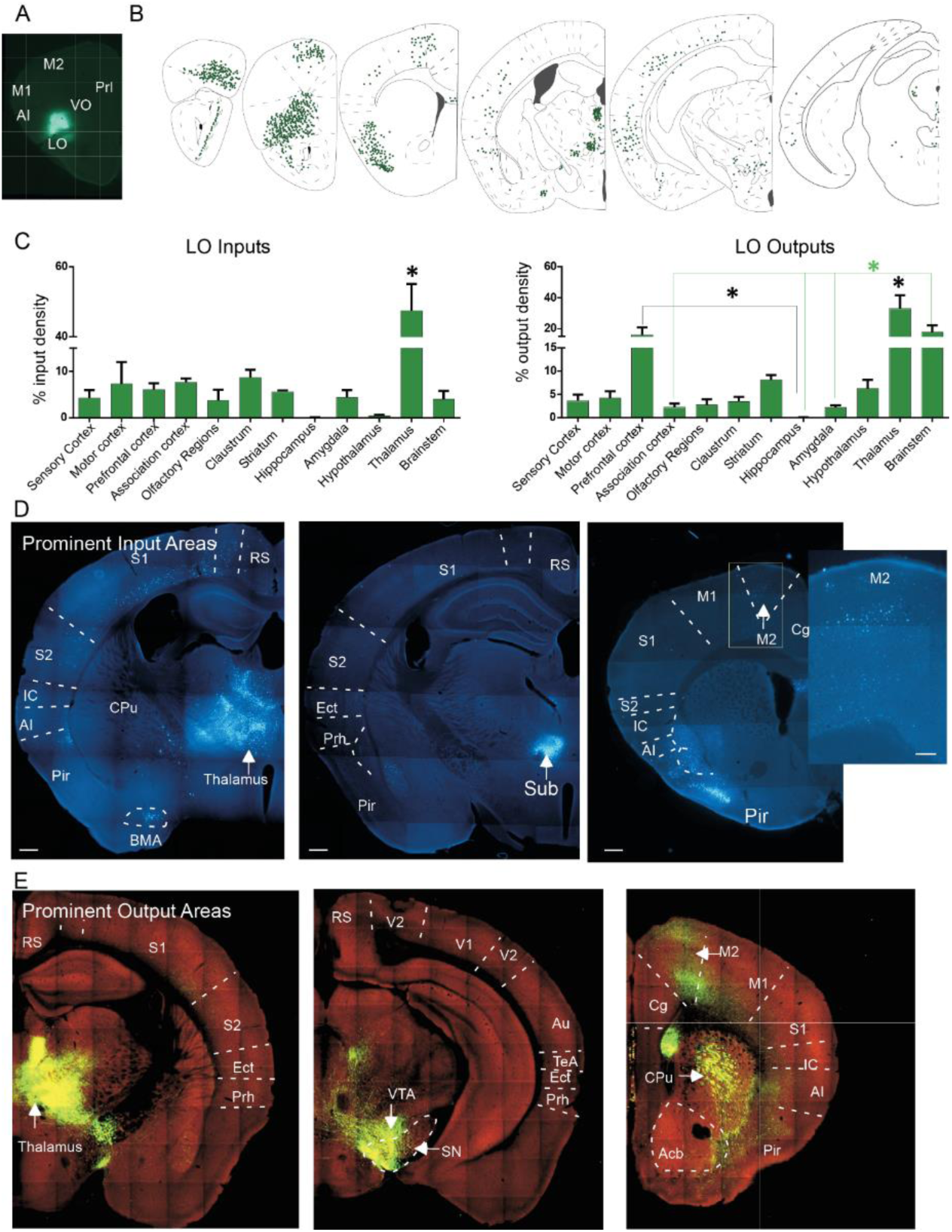
Microphotographs Depicting Inputs to and Outputs from the Lateral Orbitofrontal Cortex (LO). **A)** Representative image of a Fluorogold injection site in LO. **B)** Schematic depiction of the rostrocaudal distribution of retrogradely labeled Fluorogold-positive cells across different brain areas. **C)** Histograms showing the percentage of input cell density (left) and output projection density (right) across the main brain areas analyzed in this study. A single asterisk above a bar indicates a brain region whose input or output is significantly greater than all others. *p* < 0.001, one-way ANOVA with Tukey’s post hoc test, see also Supplementary 1 for statistics **D)** Representative images of some prominent input brain areas: most of thalamus (left) with a strong labelling within submedius thalamic nuclei (Sub; middle), and motor cortex (M2, right). The inset shows a magnified image of the retrogradely labelled cells in M2. **E)** Representative images from the Allen Brain Atlas (http://connectivity.brain-map.org/projection/experiment/112306316) showing some prominent LO output targets. Panels display projections to thalamic nuclei (left), Substantia nigra (SN) and ventral tegmental nucleus (VTA) of the brainstem (middle), and dorsal striatum (CPu) and motor cortex (M2, right). **Scale bars (D):** 200 µm; inset: 100 µm. Arrows indicate the most prominent input/output brain regions.

Quantitative retrograde tracing analysis revealed that, similar to VO, LO exhibited prominent reciprocal connectivity with the thalamus. The thalamus provided the strongest afferent input to LO (47.44 ± 7.72% input density; one-way ANOVA with Tukey post hoc test, p < 0.001), and in turn received robust efferent projections from LO (33.08 ± 8.49% output density; p < 0.001; Figure 5C). Beyond thalamic connections, LO efferents were also strongly directed to the mPFC (16.00±4.96%) and brainstem regions (17.84 ± 4.46%; see also Supplementary 1 for statistics)

Qualitative assessment further highlighted that projections from the thalamus to LO were widespread, with particularly strong labeling in the submedius thalamic nucleus (Sub, Figure 5D, middle). Additional retrograde labeling was detected in the piriform cortex (Pir) and secondary motor cortex (M2; Figure 5D, right). On the output side, dense axonal projections from LO extended to the ventral tegmental area (VTA), substantia nigra (SN), dorsal striatum (caudate-putamen, CPu), and motor cortex (see examples in Figure 5E).

### 4. Whole brain input/ output connectivity from Agranular Insula (AI)

We analyzed three retrograde and six anterograde tracer injections localized AI (Figure 2, purple). A representative retrograde injection and the schematic of rostrocaudal distribution of FG labeled cells are shown in Figure 6A,B. All our injections were localized anterior to bregma so the posterior part of AI was not targeted within this study. In that respect, the functional interpretations of AI input connectivity should be made with caution as there can be functional differences along the anterior-posterior axis in AI (Gehrlach, Weiand et al. 2020).

**Figure 6:**
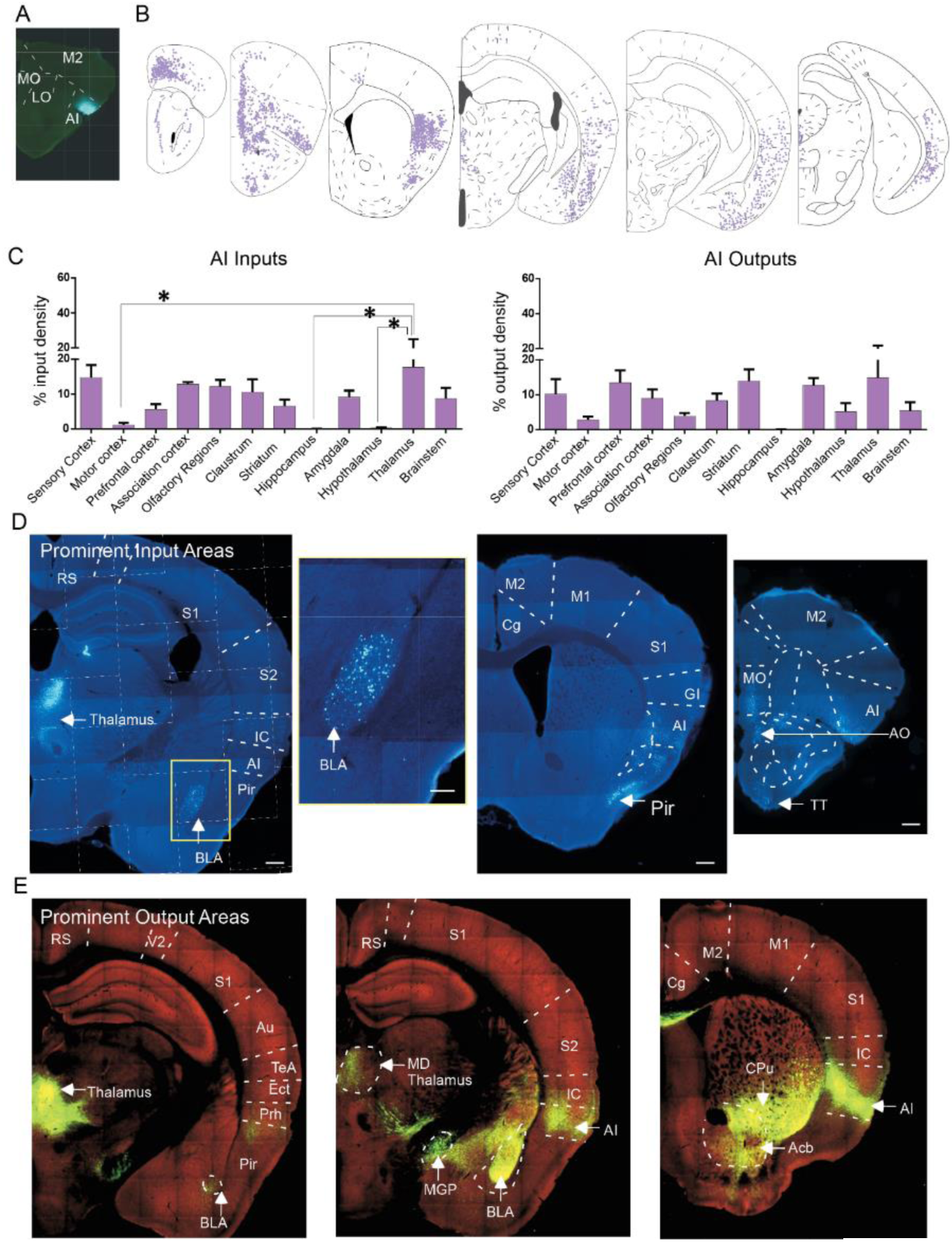
Microphotographs Depicting Inputs to and Outputs from the Agranular Insular Cortex (AI). **A)** Representative image of a Fluorogold injection site in AI. **B)** Schematic depiction of the rostrocaudal distribution of retrogradely labeled Fluorogold-positive cells across different brain areas. **C)** Histograms showing the percentage of input cell density (left) and output projection density (right) across the main brain areas analyzed in this study. *p* < 0.05, one-way ANOVA with Tukey’s post hoc test, see also Supplementary 1 for statistics. **D)** Representative images of some prominent input brain areas: thalamus and basolateral amygdala (BLA, left) with the inset showing a magnified view of retrogradely labelled cells within BLA. Other prominent inputs are from olfactory regions such as Piriform cortex (Pir, middle), tenia tecti (TT) and Accessory olfactory areas (AON, right). **E)** Representative images from the Allen Brain Atlas (http://connectivity.brain-ap.org/projection/experiment/112596790) showing some prominent AI output targets. Panels display projections to thalamic nuclei (left and middle), basolateral amygdala (BLA, middle) and all the striatal areas, medial globus pallidus (MGP, middle), Caudate putamen (CPu) and ventral Accumbens (Acb; right). **Scale bars (D):** 200 µm; inset: 100 µm. Arrows indicate the most prominent input/output brain regions.

Quantitative analysis of retrograde tracing revealed that AI received input from multiple brain regions, without a single dominant afferent source (Figure 6C). Among these, the thalamus (17.77± 7.37% input density) was one of the stronger contributors compared to motor cortex (1.22±0.56%), hippocampus (0.06±0.06%), or hypothalamus (0.27±0.19%). Similarly, quantitative analysis of anterograde tracing did not identify a singularly prominent efferent target. Instead, AI projected broadly, with notable outputs to mPFC (13.51±3.48% output density), striatum (13.94±3.30%), amygdala (12.68±2.09%), and thalamus (14.85±6.81%, see also Supplementary 1 for statistics.

Qualitative examination provided additional findings. The mediodorsal thalamic nucleus (MD) within the different thalamic nuclei and the basolateral amygdala (BLA) within amygdala nuclei exhibited robust retrograde labeling. Sensory cortices, including the Pir, as well as subcortical olfactory regions such as the accessory olfactory area (AO), also contributed to AI inputs (Figure 6D). Anterograde tracing confirmed widespread AI projections to the thalamus, particularly the MD nucleus, as well as to the BLA and the striatopallidal complex. Within the striatum, dense axonal labeling was observed in both the caudate-putamen (CPu) and nucleus accumbens (Acb), as well as in the medial globus pallidus

### 5. Whole brain input/ output connectivity from Dorsolateral OFC (DLO)

We analyzed three retrograde and two anterograde tracer injections targeting DLO (Figure 2, red). A representative retrograde injection and the schematic of rostrocaudal distribution of FG labeled cells are shown in Figure 7A,B.

**Figure 7:**
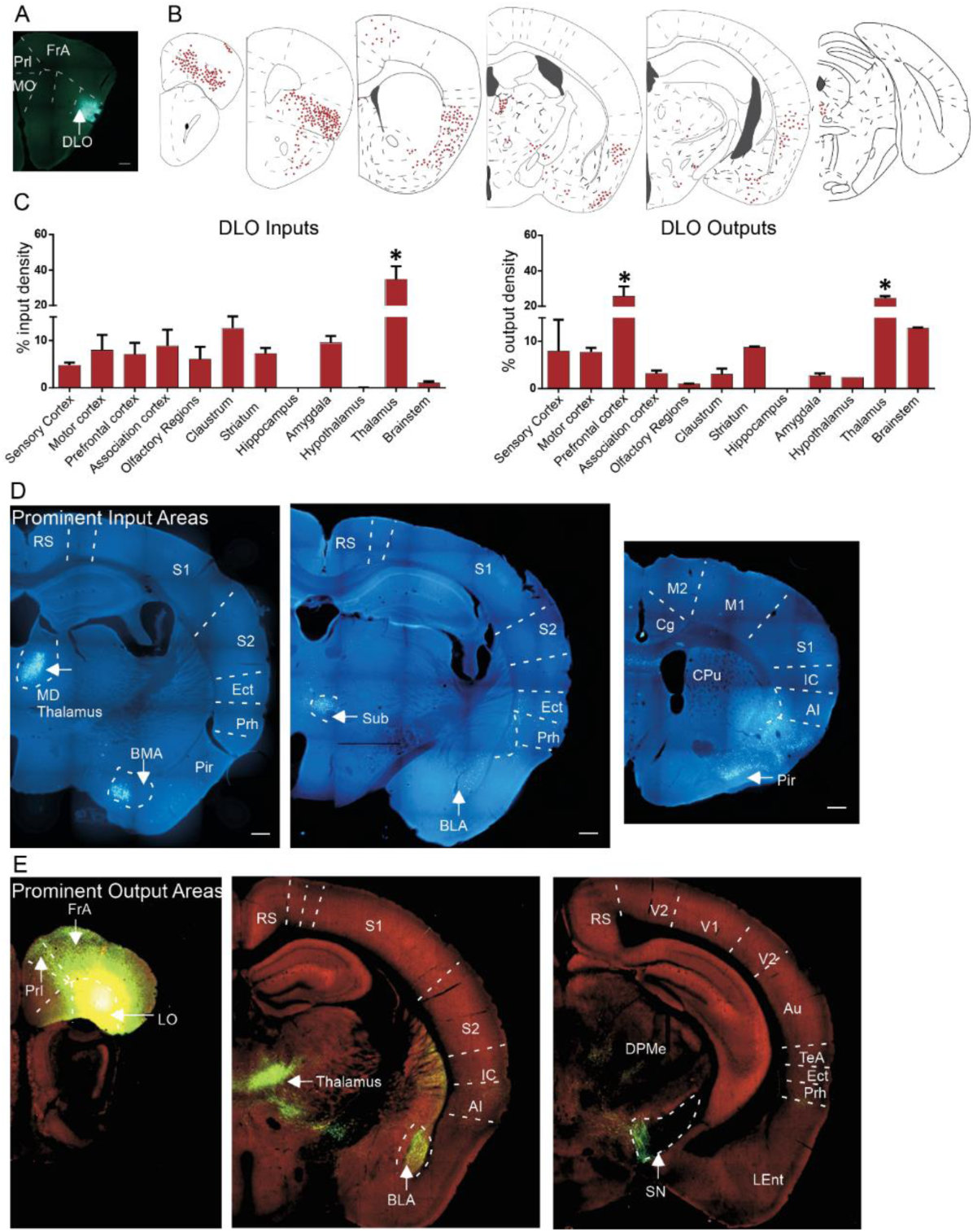
Microphotographs Depicting Inputs to and Outputs from the Dorsolateral Orbitofrontal Cortex (DLO). **A)** Representative image of a Fluorogold injection site in DLO. **B)** Schematic depiction of the rostrocaudal distribution of retrogradely labeled Fluorogold-positive cells across different brain areas. **C)** Histograms showing the percentage of input cell density (left) and output projection density (right) across the main brain areas analyzed in this study. A single asterisk above a bar indicates a brain region whose input or output is significantly greater than all others, *p* < 0.05, one-way ANOVA with Tukey’s post hoc test, see also Supplementary 1 for statistics. **D)** Representative images of some prominent input brain areas: thalamic nuclei mostly MD thalamus (left) and submedius (Sub, middle), basolateral amygdala (BLA, middle) and sensory piriform cortex (right). **E)** Representative images from the Allen Brain Atlas (http://connectivity.brain-ap.org/projection/experiment/170721670) showing some prominent AI output targets. Panels display projections to prefrontal cortical areas (left and middle), thalamus (middle) and Substantia nigra of the brainstem (SN, right). Scale bars (D): 200 μm. Arrows indicate the most prominent input/output brain regions.

Quantitative retrograde tracing analysis revealed that, similar to LO, the thalamus provided the strongest input to DLO (34.73 ± 7.38% input density), significantly greater than all other brain areas (p< 0.0001, one-way ANOVA with Tukey post hoc test). Anterograde analysis confirmed that DLO projected robustly to the thalamus (24.54 ± 1.16% output density), as well as to the mPFC (25.70 ± 5.49%; Figure 7C, see also Supplementary 1 for statistics).

Qualitative assessment further indicated that within the thalamus, both MD and Sub nuclei contributed strong inputs to DLO. Additional retrograde labeling was observed within the basomedial amygdala (BMA) and in olfactory-related cortices, such as the Pir (Figure 7D). On the efferent side, DLO displayed strong intra-OFC projections, particularly to LO, as well as outputs to thalamic nuclei, the BLA, and brainstem regions including the SN (Figure 7E).

Given the widespread connectivity of OFC subdivisions with both cortical and subcortical areas, we further analyzed their connectivity with several key brain regions, using both quantitative analysis and qualitative examination. In some cases, labeling was confined to restricted areas within larger brain region and thus did not reach quantitative significance; nevertheless, these patterns are described to emphasize similarities and differences among OFC subdivisions.

### 6. Comparison of connectivity with sensory cortices among OFC subdivisions

OFC is a multisensory area (Banerjee, Parente et al. 2020, Sharma and Bandyopadhyay 2020, Tripathi, Sato et al. 2021). Most primary and association sensory cortices both sent to and received projections from all OFC subdivisions, though distinct biases in afferent connectivity were observed (Figure 8A). Medial OFC regions (MO and VO) received stronger inputs from visual cortices (see Figure 8, left plot “visual”), whereas central/lateral regions received inputs from somatosensory cortices. In contrast, the most peripheral division, AI, was preferentially targeted by auditory, olfactory, and gustatory cortices. Indeed, AI received the most prominent auditory and chemosensory (gustatory and olfactory) input connectivity among all OFC subdivisions (see ANOVAs followed by post-hoc tests in Figure 8A, and Supplementary 2).

**Figure 8:**
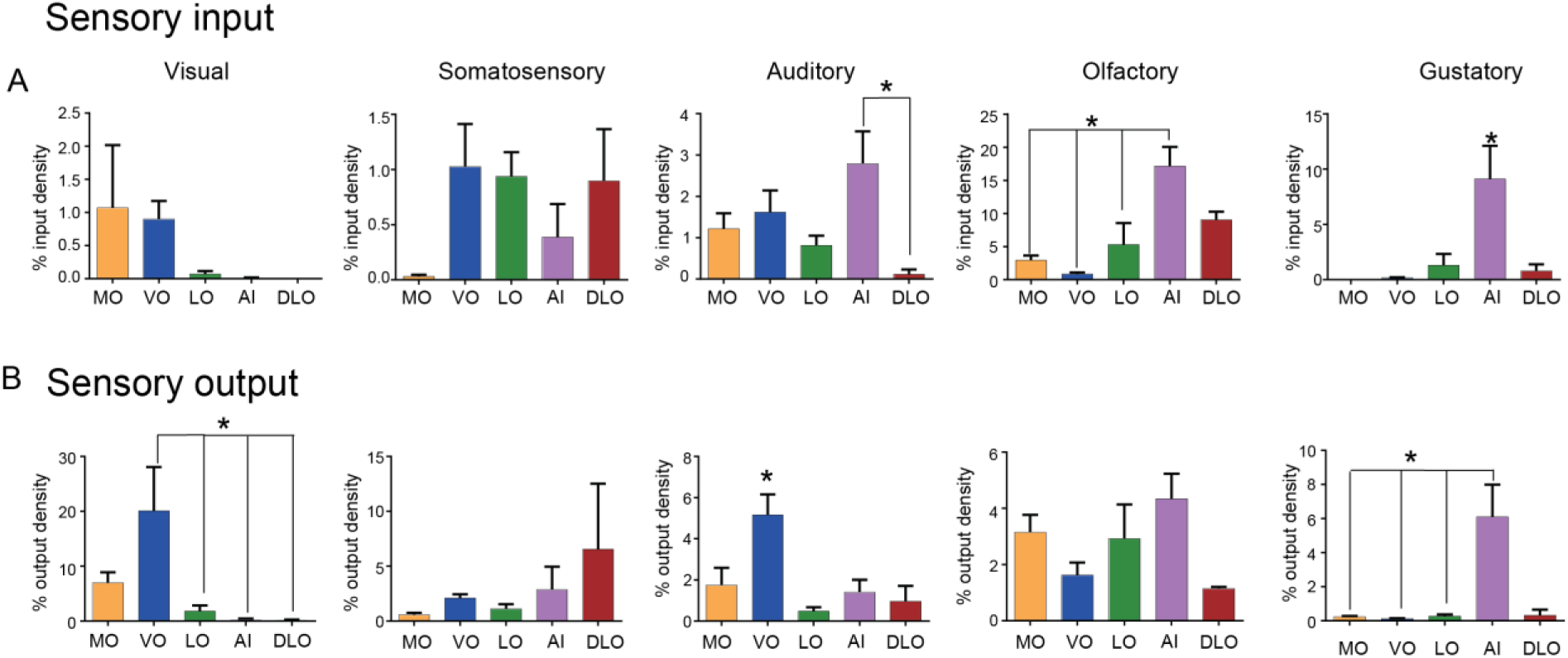
Multisensory Nature of the Orbitofrontal Cortex. **A)** Histograms depict the distribution of inputs from various sensory cortices to orbitofrontal cortex (OFC) subdivisions. Visual cortex projects more to ventral (VO) and medial OFC (MO), whereas somatosensory inputs are broadly distributed across OFC subdivisions, with the exception of MO, which receives minimal somatosensory input. The agranular insular cortex (AI) receives significantly stronger input from auditory, olfactory, and gustatory areas compared to other OFC subdivisions. **(B)** Output patterns from OFC to sensory cortices are shown in the corresponding histograms. VO exhibits the strongest projections to the visual cortex among OFC subdivisions. Overall, output to the somatosensory cortex is minimal. Auditory cortex receives maximal input from VO, while all OFC subdivisions send projections to olfactory areas. The gustatory insula receives its strongest input from AI, while there is minimal output from other OFC subdivisions. p < 0,05, 1-way ANOVA with Tukey, see also Supplementary 2 for statistics.

Efferent connectivity patterns were less segregated (Figure 8B). Visual cortices received their strongest input from VO, while somatosensory cortices received few efferent projections from any OFC subdivision. VO also projected significantly more strongly to auditory cortex, whereas olfactory cortices received inputs from all OFC subdivisions. Notably, AI projected densely to the gustatory region of the insula, paralleling its afferent connectivity (see ANOVAs followed by post-hoc tests in Figure 8B, and Supplementary 2). Together, these findings suggest that although all OFC subdivisions are essentially multisensory in nature, their input–output patterns are biased toward distinct sensory modalities.

The visual comparison between the upper and lower panels of Figure 8 allows to note the pattern similarity between the cortical association inputs and outputs areas, particularly for the most medial MO, connected in a prominent way with acoustic and visual cortices, and the most lateral AI and DLO, receiving predominant acoustic and chemoreceptive inputs.

### 7. OFC connectivity with the striato-pallidal complex

We observed strong but spatially confined inputs from both dorsal and ventral striatum to OFC subdivisions. Closer inspection revealed well-defined clusters of retrogradely labeled cells within the striatopallidal complex, organized in a medio–lateral topography within the dorsal striatum (Figure 9A,B). Specifically, MO received inputs from medial CPu, VO from more medio-central CPu, and LO from lateral CPu.

**Figure 9:**
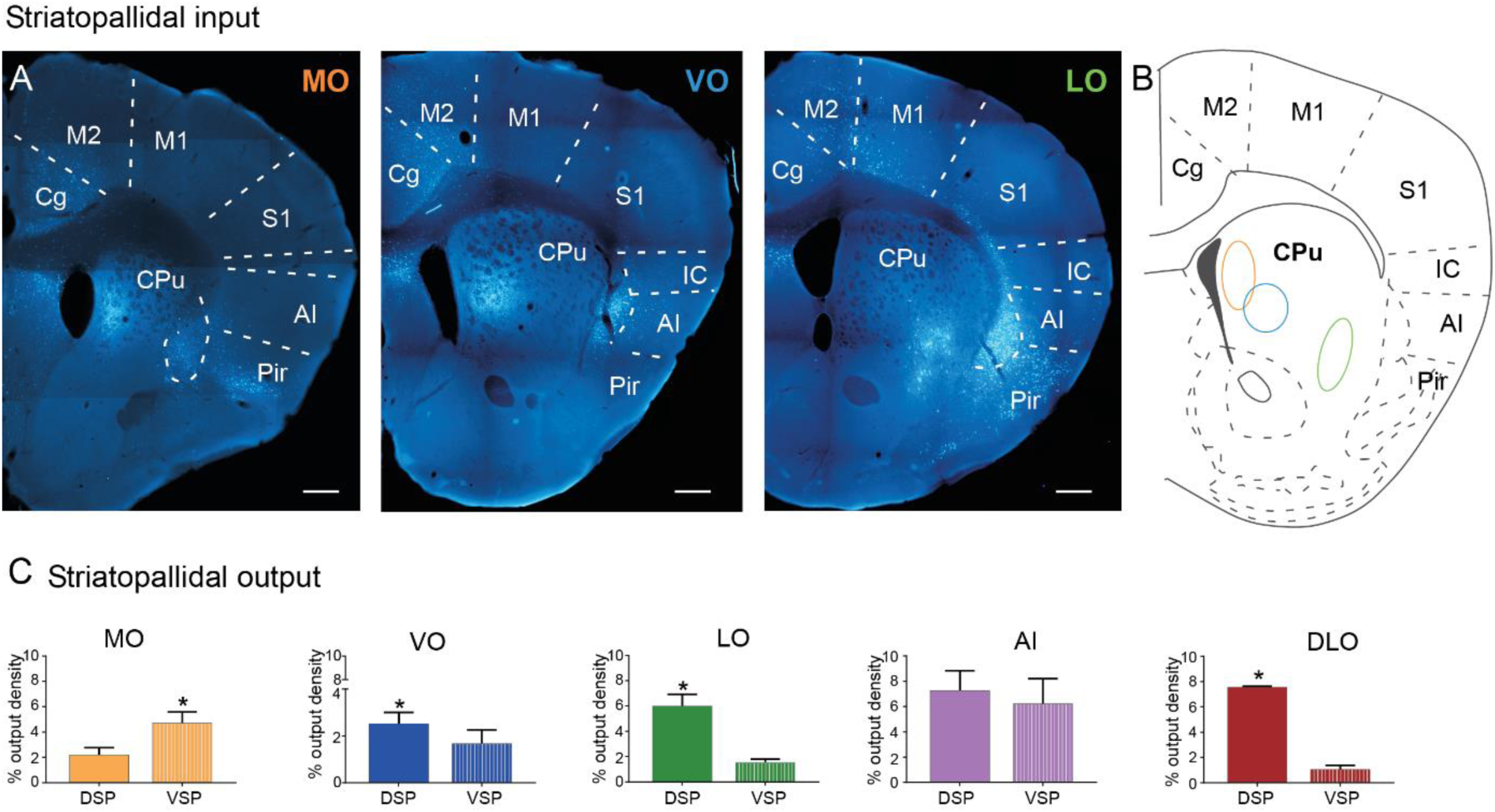
Striatopallidal Connectivity with Orbitofrontal Cortex. **A)** Representative microphotographs demonstrate focal yet dense direct projections from the dorsal striatum to various orbitofrontal cortex (OFC) subdivisions. These projections exhibit a clear topographic organization along the medio-lateral axis, as illustrated in the accompanying schematic (right). The colors of the contours are representative of the respective OFC subdivision. **B)** Quantitative analysis reveals distinct output patterns from OFC subdivisions to the dorsal and ventral striatopallidal complex (DSP and VSP respectively). Starting from the left, Medial OFC (MO) predominantly targets the ventral striatum and pallidum. In contrast, ventral (VO) and lateral OFC (LO) show stronger projections to the dorsal striatopallidal regions. Agranular insular cortex (AI) displays a unique profile with comparable projections to both dorsal and ventral components. Dorsolateral OFC (DLO) resembles LO, with a dominant projection pattern toward the dorsal striatopallidal complex. p<0.05, paired student t-test, see also Supplementary 3 for statistics. **Scale bars (A):** 200 µm.

Quantitative output density analysis (Figure 9C) further supported subdivision specific biases. MO sent significantly stronger output to the ventral striato-pallidal complex (VSP; nucleus accumbens and ventral pallidum), whereas VO and LO sent more substantial output to the dorsal striato-pallidal complex (DSP; CPu and globus pallidus). DLO mirrored VO and LO, showing preferential output to DSP, while AI exhibited relatively similar output to both VSP and DSP. See subdivision specific paired T-tests in Figure 9C and supplementary 3.

### 8. OFC connectivity with the hippocampus

Strong but restricted retrograde labeling was also observed in the hippocampus, despite not being quantitatively significant in comparison with all brain input areas. Specifically, labeled pyramidal neurons in CA1 and the ventral subiculum projected selectively to MO and VO (Figure 10A,B). No hippocampal labeling was observed following tracer injections in LO, DLO, or AI.

**Figure 10:**
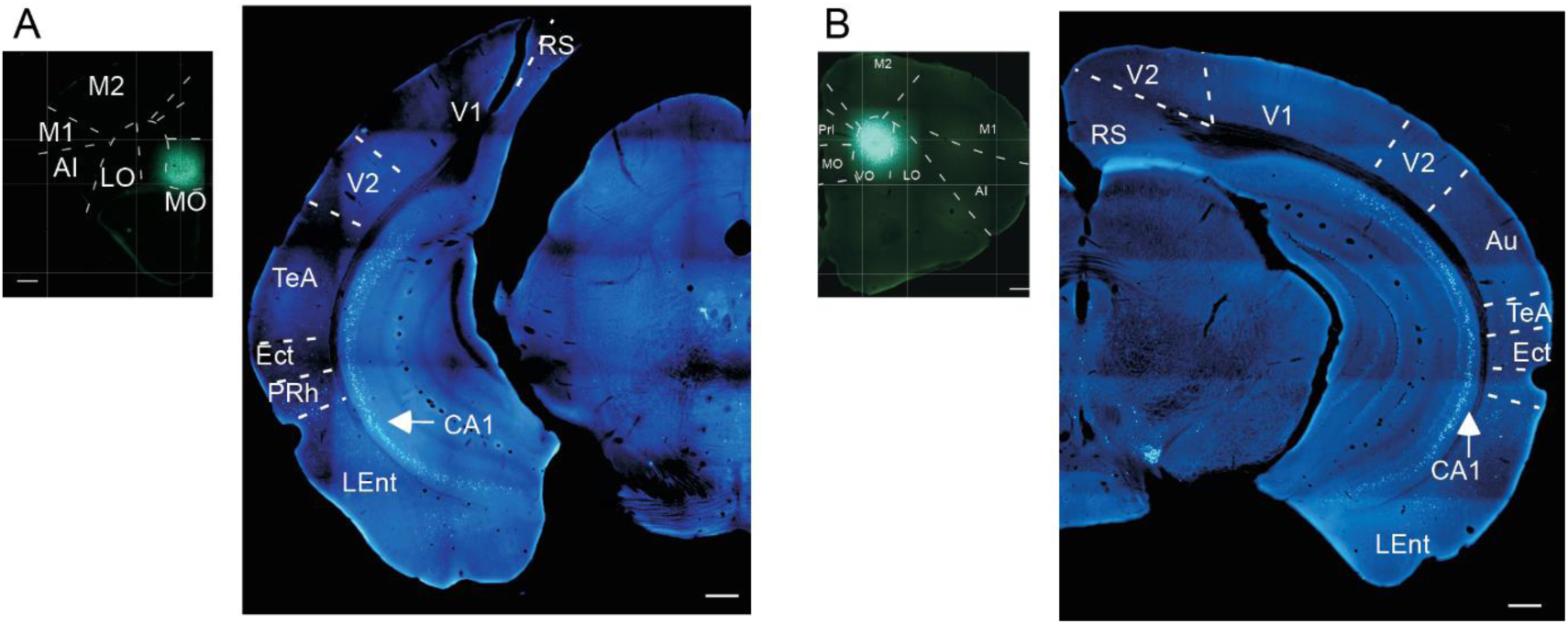
Direct Hippocampal Connectivity to OFC Subdivisions. Retrograde tracer injections confined to medial OFC (MO; **A)** and ventral OFC (VO; **B,**) result in robust but spatially restricted labeling within the CA1 region of the hippocampus, as shown in the corresponding panels on the right. These findings indicate a direct, topographically organized projection from hippocampal CA1 to specific medially located OFC subdivisions. **Scale bar**= 200 µm

### 9. OFC connectivity with brainstem neuromodulatory regions

Clear differences among subdivisions were observed in inputs from dopaminergic and serotonergic brainstem nuclei (Figure 11). VO received strong dopaminergic input from the substantia nigra, including both SNr and SNc (Figure 11A,B), compared with other OFC subdivisions. In contrast, AI was predominantly innervated by serotonergic neurons from the raphe nuclei, in comparison with VO, LO, DLO, and MO (Figure 11C,D). See ANOVAs followed by post-hoc tests in Figure 11A,C and Supplementary 3).

**Figure 11:**
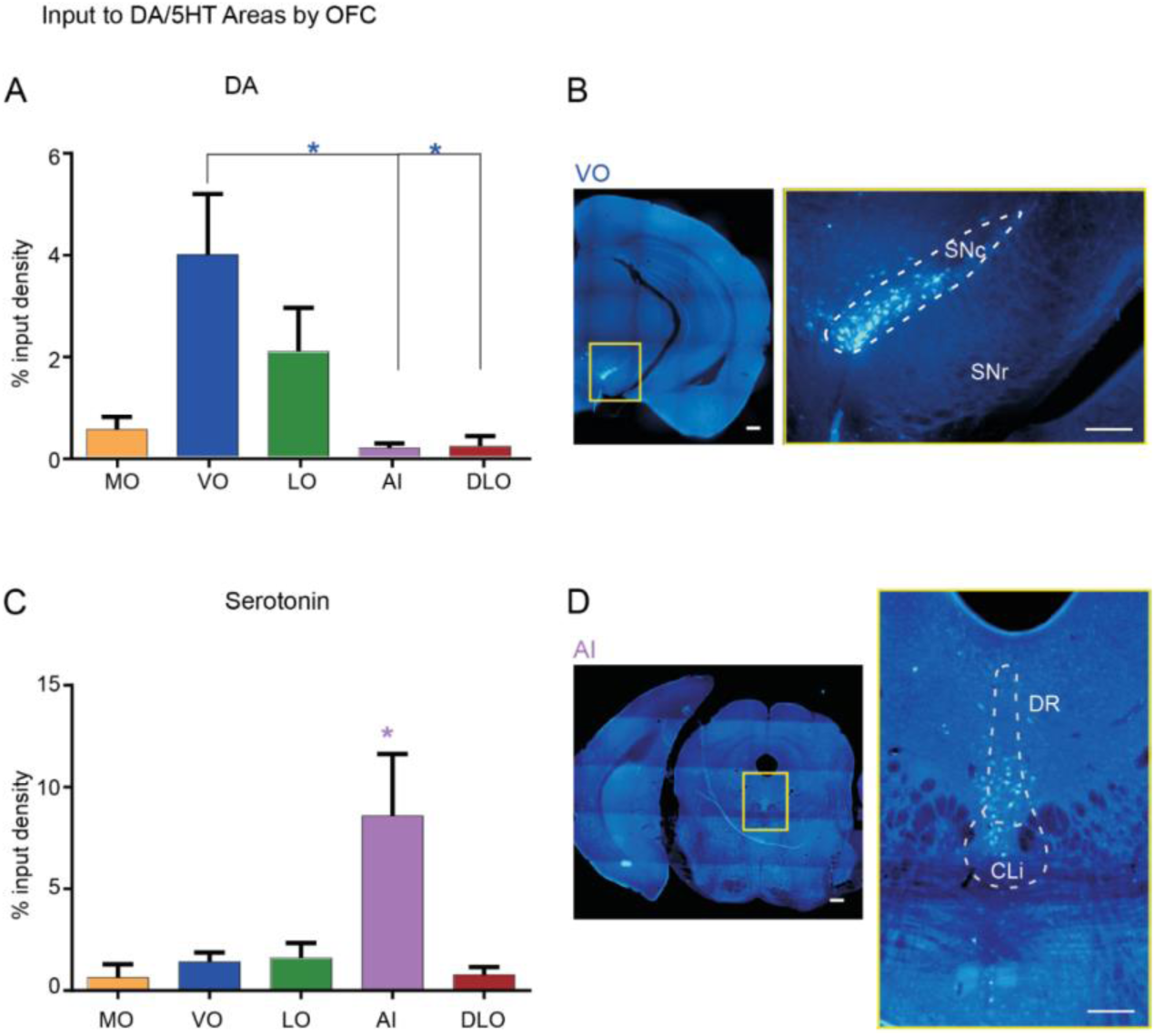
Neuromodulatory Input to OFC Subdivisions. **A)** Histograms reveal significantly greater input to ventral OFC (VO) from dopaminergic brainstem regions. **B)** Representative microphotograph of a coronal brain section following a retrograde tracer injection in VO shows labeled neurons in the substantia nigra, with the inset displaying a magnified view of labeled neurons. **C)** Histograms indicate a significantly higher input from serotonergic brainstem nuclei to the agranular insular cortex (AI) compared to other OFC subdivisions. **D)** Corresponding microphotograph after AI injection shows retrograde labeling in serotonergic areas, including the dorsal raphe (DR) and caudal linear raphe (CLI). The inset provides a magnified view of labeled serotonergic neurons in these regions. p < 0,05, 1-way ANOVA with Tukey, see also Supplementary 3 for statistics **Scale bars (B, D):** 200 µm; inset: 100 µm.

### 10. Comparison among OFC subdivisions of whole brain areas inputs and outputs

To obtain a comprehensive overview of the differences in input and output projection patterns across OFC subdivisions, we compared the retrograde and anterograde connectivity across 12 main brain areas (Figure 12).

**Figure 12.**
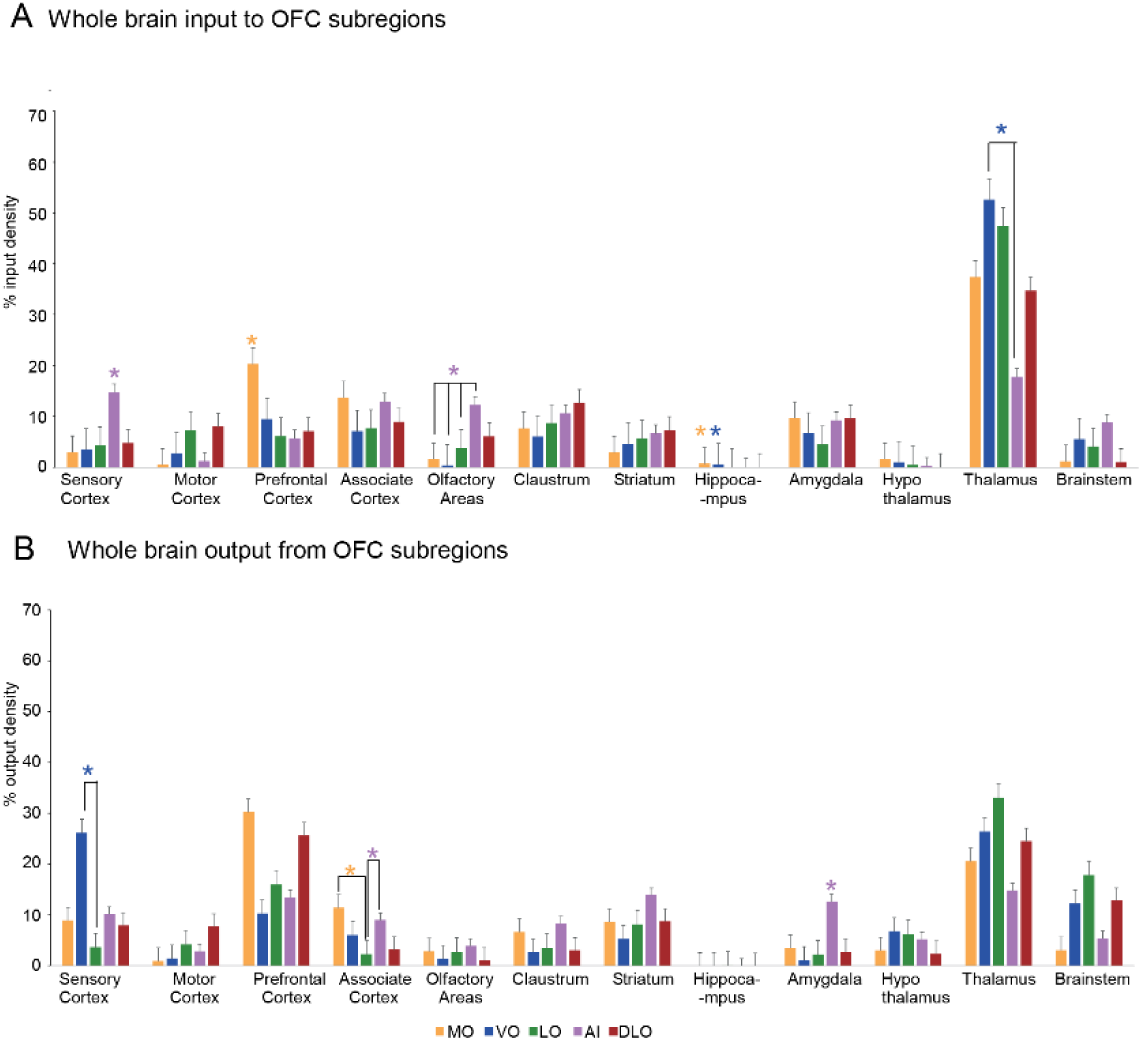
Whole-Brain Mapping of Inputs to and Outputs from OFC Subdivisions. **A)** Quantitative comparison of inputs to all five orbitofrontal cortex (OFC) subdivisions from 12 major brain area groups analyzed in this study. Input strength is represented as the percentage of total input cell density for each OFC subdivision. **B)** Corresponding comparison of output projections from the same OFC subdivisions to the 12 major brain area groups. Output values are also expressed as a percentage of total output projection density, enabling direct comparison of relative connectivity patterns across the brain. A single asterisk above a bar indicates a brain region whose input or output is significantly greater than all others,* *p* < 0.05, one-way ANOVA with Tukey’s post hoc test. See also Supplementary 4 for statistics.

Inputs to OFC exhibited a heterogeneous distribution along the medial–lateral axis (Figure 12A). Medial MO received its strongest projections from the mPFC and hippocampus. The hippocampus also innervated neighboring VO, which in addition received substantial projections from the thalamus. Thalamic input remained the dominant source for more lateral divisions, including LO and DLO. In contrast, the peripheral AI received its primary inputs from sensory and olfactory cortical areas. Note marked projections from basolateral amygdala (BLA) to both MO and AI (see microphotographs in Figures 3D and 6D, respectively).

Analysis of the efferent projections revealed that, although the thalamus was a common target of all OFC subdivisions, clear differences in output patterns could also be delineated (Figure 12B). Moving medial to lateral, MO projected predominantly to the mPFC and association cortical areas, whereas VO sent stronger projections to sensory cortices and brainstem regions. LO projected heavily to the brainstem, while DLO distributed its outputs between the mPFC and brainstem. Notably, AI was the only OFC subdivision to provide strong projections to the amygdala (12.68±2.09% output density, p<0.05, one-way ANOVA with Tukey, distinguishing it from other OFC subdivisions. See also Supplementary 4 for statistics.

### 11. Input/output map similarities reveal distinct connectivity clusters within OFC

We next assessed the degree to which the OFC subdivisions shared similar or distinct origins for their inputs and targets for their outputs, which might reveal the existence of anatomical differences among these subdivisions that are known to be functionally different in humans and non-human primates. For each subdivision, we constructed representative input and output distributions, which we then compared pairwise.

From these results, it was evident that VO and LO, and to a slightly lesser extent DLO and MO, shared strongly similar input distributions, which were dissimilar to the input distribution of AI (Figure 13A), as confirmed by the results of hierarchical clustering (Figure 13B). This dissimilarity is likely driven by the greater proportion of inputs to AI from sensory cortex, olfactory areas and brainstem, and the substantially smaller proportion of inputs from thalamus compared to the other four subdivisions (see also Figure 12A). However, despite the similar inputs, we found notable differences in the output distributions, which set VO, MO and AI apart from each other and from a putative LO and DLO cluster (Figure 13C,D). While output distributions were less similar overall among all subdivisions, the resulting differences were likely due to a strong preference for output to sensory cortex and against output to mPFC in VO compared to all other subdivisions, while AI and MO were less likely to project to thalamus and brainstem and more likely to project to association cortex, as well as amygdala (in the case of AI) and mPFC (in the case of MO), than all others.

**Figure 13.**
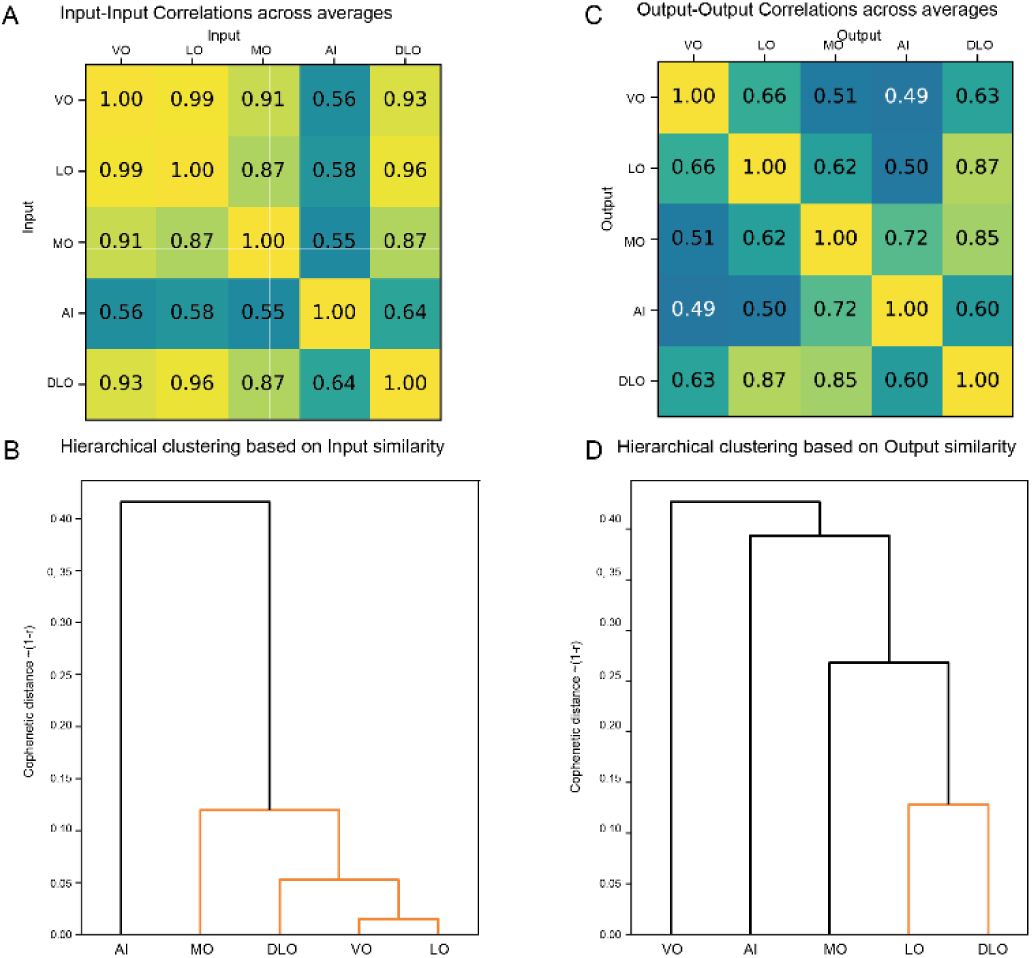
Cluster analysis of the input and output connectivity of the various OFC subdivisions. **A)** Correlation between the distributions of inputs for each pair of subdivisions. **B)** Hierarchical clustering based on the results shown in A. Note the relatively high similarity between inputs to all subdivisions except AI. **C)** Correlation between the distributions of outputs for each pair of subdivisions. **D)** Hierarchical clustering based on the results shown in C. Despite the similarities in input profiles, output profiles reveal greater differences between subdivisions and suggest that VO, AI and MO might each be distinct enough to be considered a separate cluster.

Note that the results of the clustering analysis based on inputs and outputs are independent. For instance, VO and LO have extremely similar input connectivity but divergent output connectivity; nevertheless, one relevant finding observed in both input and output cluster analysis is the relative segregation of AI from the remaining areas.

Finally, we compared each subdivision’s input distribution to its own output distribution, in order to investigate which subdivisions might be more likely to be part of functional loops. The highest input-output correlations were found for LO (*r=0.81*) and MO (*r=0.79*), compared with VO (*r=0.64*), AI (*r=0.56*), and DLO (*r=0.54*), potentially driven by strong reciprocal connections with Thalamus in MO and LO, as well as with mPFC in MO.

Taken together, these observations suggest that, when input connectivity, output connectivity, and putative loop distributions are investigated simultaneously, LO and DLO share rather similar anatomical connectivity, which is differentiated from that of VO by virtue of the latter’s very distinct output profile, while AI is clearly different in both inputs and outputs (see results of the concatenated input and output correlation averages of Figure 14; see also Supplementary Figure 1). These observations point to a potential anatomical substrate for the existence of functional differences among OFC subdivisions also in mice.

**Figure 14.**
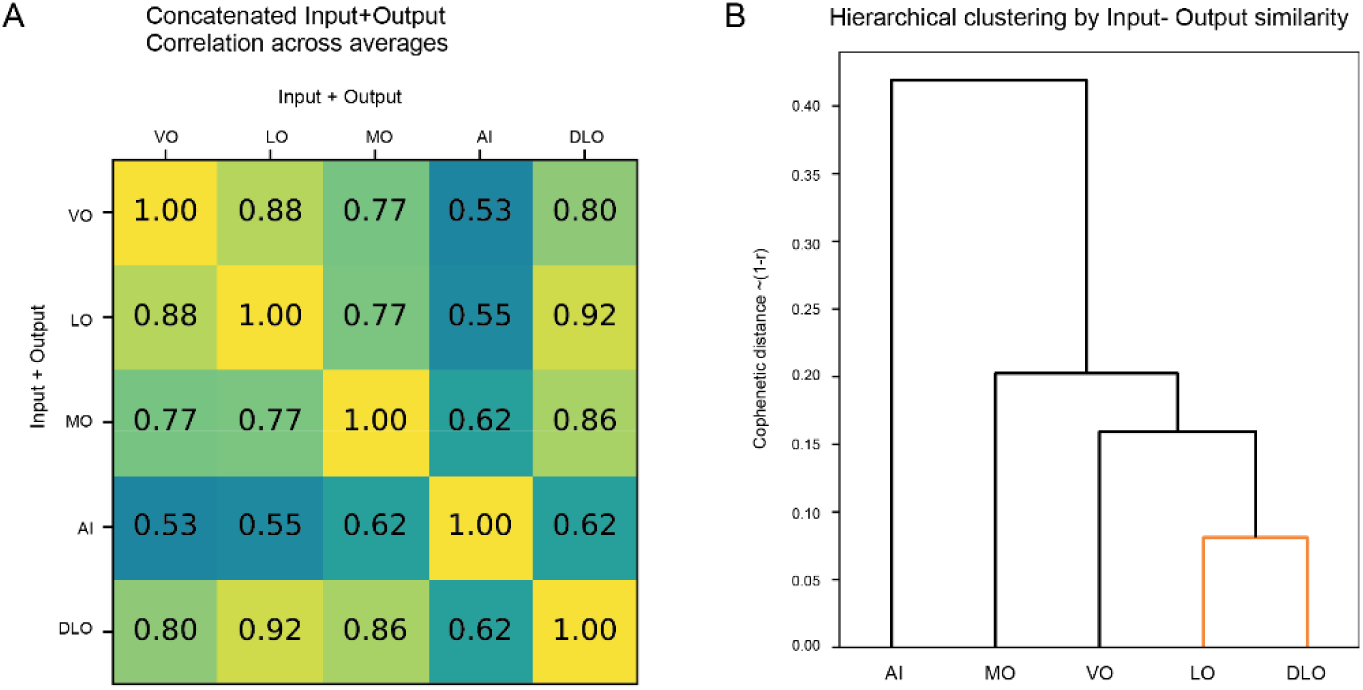
Analysis of the concatenated inputs and outputs. **A)** Correlation between the concatenated distributions of inputs and outputs simultaneously, for each pair of subdivisions. **B)** Hierarchical clustering based on the results shown in A. The results also support a cluster consisting of LO and DLO, while keeping the other subdivisions separate, and are consistent with the results shown in Figure 13.

**Supplementary Figure 1. Related to Figure 13.**
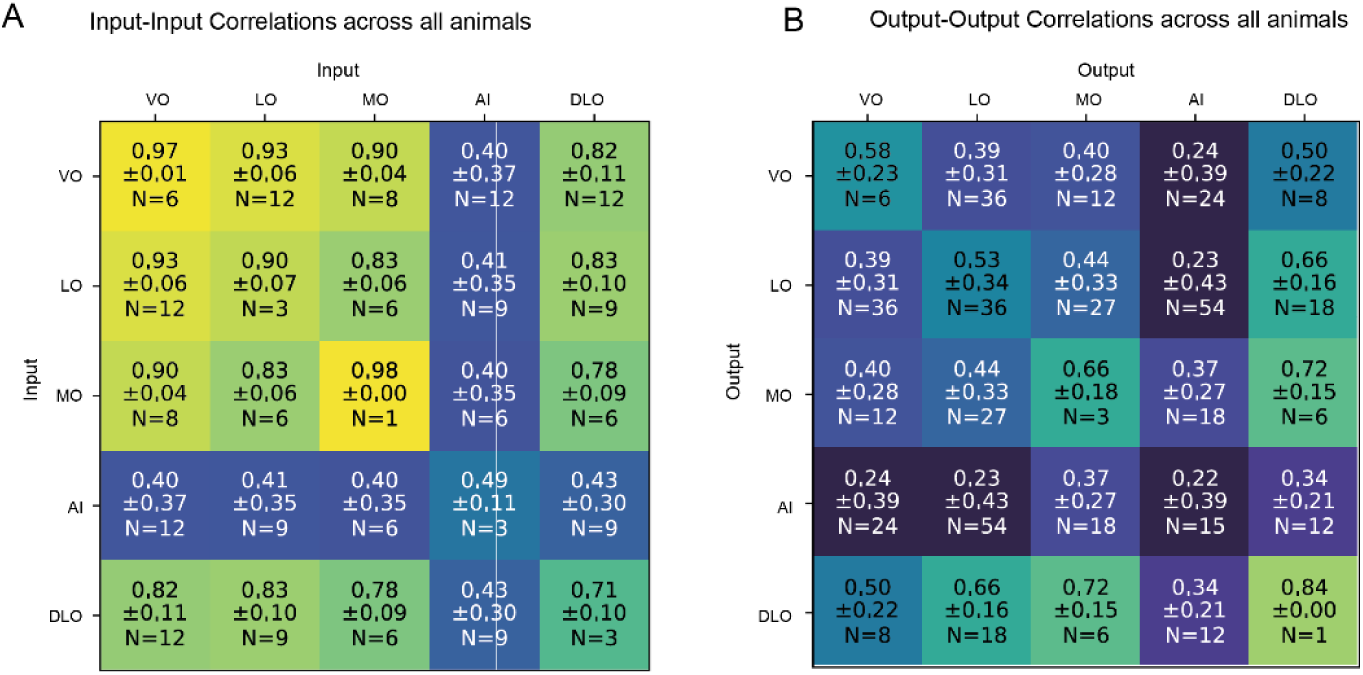
**A)** Correlation between the distributions of inputs for each pair of subdivisions, calculated separately across all suitable pairs of animals and then averaged, instead of being calculated on the representative distributions of each structure as shown in main text. Results do not differ substantially despite the presence of inter-animal variation. **B)** Correlation between the distributions of outputs for each pair of subdivisions, calculated separately across all suitable pairs of animals and then averaged, instead of being calculated on the representative distributions of each structure as shown in main text. Results do not differ substantially despite the presence of inter-animal variation. We also calculated input-output correlations across animals instead of representative averages (not shown), with the following results: MO (*r=0.71±0.19, N=6*), LO (*r=0.53±0.33, N=27*), VO (*r=0.46±0.33, N=16*), DLO (*r=0.46±0.13, N=6*), AI (*r=0.26±0.27, N=18*).

### 12. Thalamic input connectivity patterns of AI and MO are similar

In cortical neuroanatomy, the thalamic innervation pattern is often used as criterion to establish area identities and parcellations (e.g. (Miller-Hansen and Sherman 2022, Howell, Warrington et al. 2025). Moreover, given that the thalamus emerged as both a prominent source of input and a major target of efferent projections for all OFC subdivisions, we further subdivided the thalamus into eight major divisions based on Roy et al. (Nature Neuroscience, 2022) to characterize the OFC-thalamus connectivity matrix in greater detail (Figure 15). Since no anterograde or retrograde labeling was observed in the posterior and lateral subdivisions of the thalamus, they have not been considered in generating the histograms (Figure 15).

**Figure 15.**
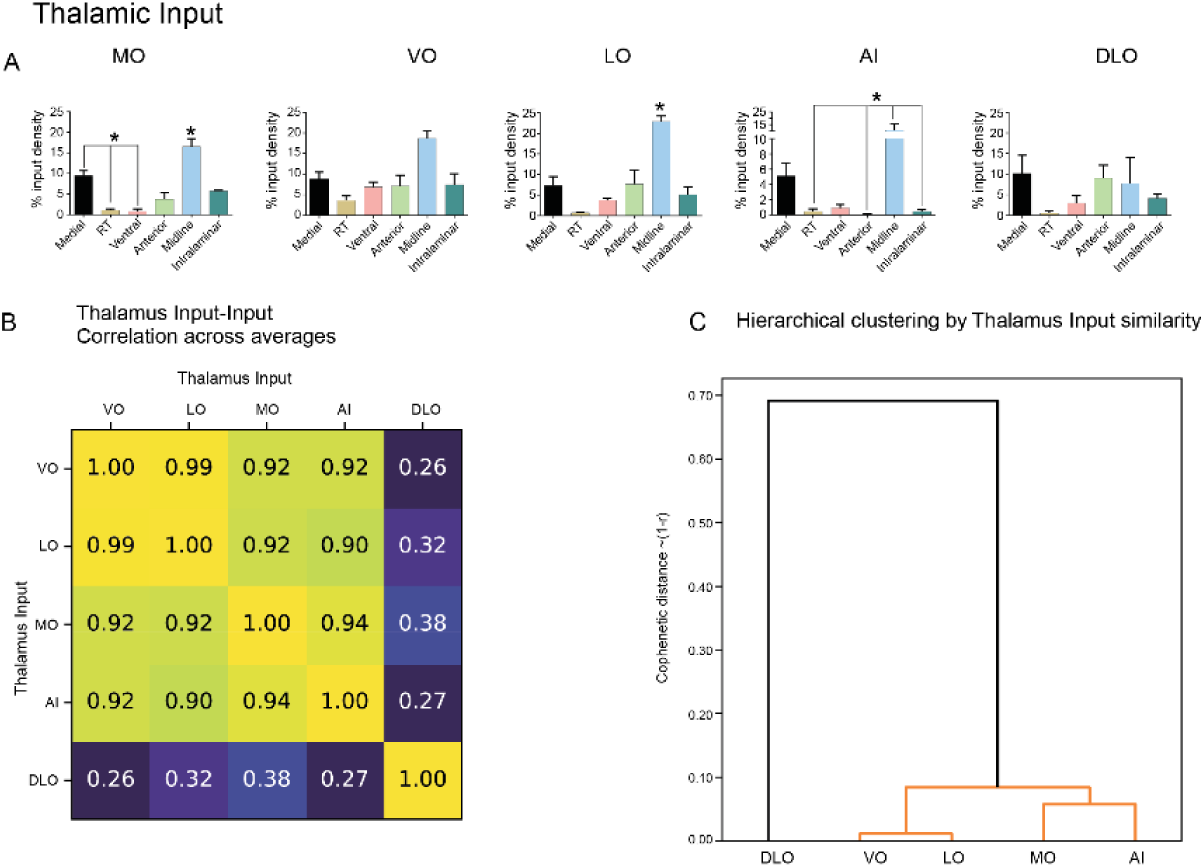
Thalamic input connectivity pattern of OFC subdivisions. **(A)** Histograms show the distribution of thalamic inputs to OFC subdivisions. Midline thalamic nuclei provide strong input to most OFC subdivisions (MO, VO, LO, and AI), with the exception of DLO, which instead receives broad input from multiple thalamic nuclei. **B)** Correlation between the average distributions of thalamic inputs, for each pair of subdivisions. Note the overall high similarity apart from DLO. **C)** Hierarchical clustering based on the results shown in B, highlighting VO/LO and MO/AI as two pairs of structure with highly similar respective thalamic input patterns. A single asterisk above a bar indicates the thalamic nucleus with significantly higher input than the other thalamic nuclei, \**p* < 0.05, one-way ANOVA with Tukey’s post hoc test. See also Supplementary 5 for statistics

**Supplementary Figure 2. Related to Figure 15.**
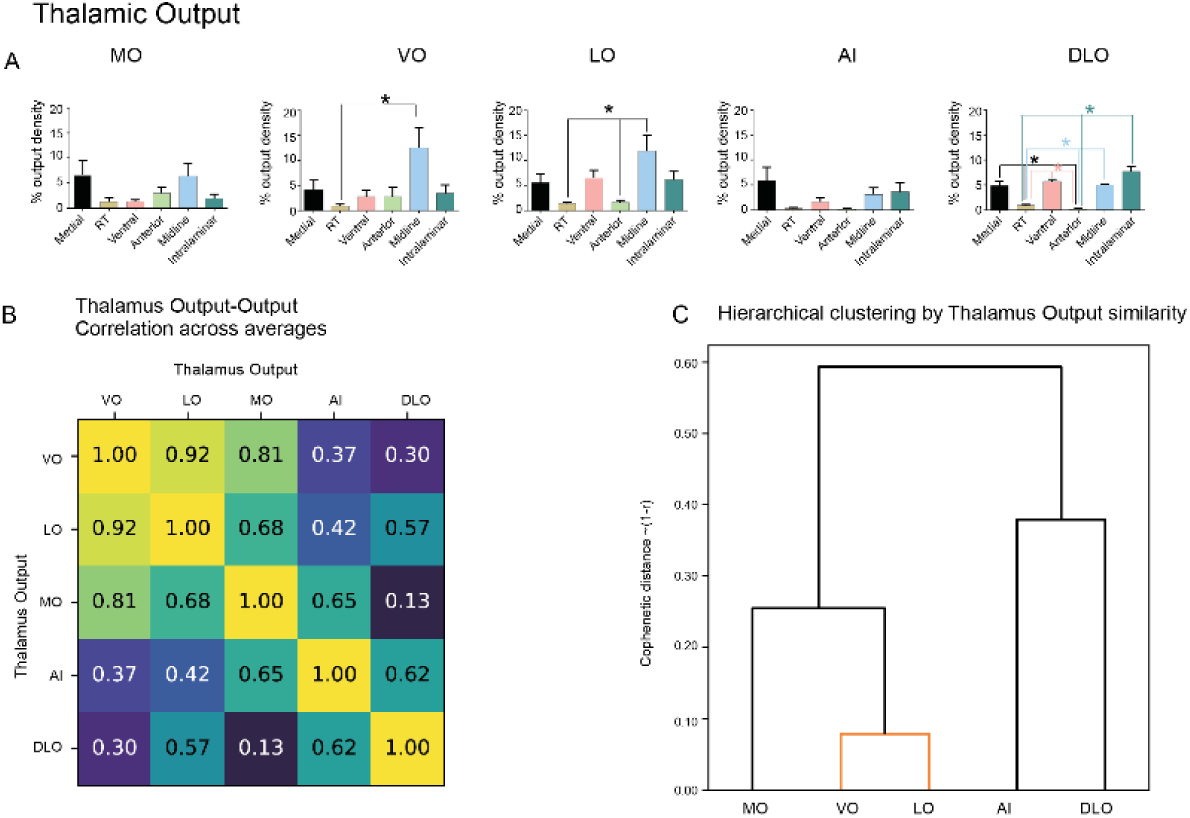
Thalamic outputs from the different OFC subdivisions. **(A)** Histograms illustrate OFC outputs to thalamic nuclei. MO and AI send only sparse projections, whereas VO and LO provide significant output to midline thalamic nuclei. DLO, in contrast, projects broadly across diverse thalamic nuclei. **B)** Correlation between the average distributions of thalamic outputs, for each pair of subdivisions. **C)** Hierarchical clustering based on the results shown in B, revealing relatively diverse output patterns apart from VO and LO which are highly similar. \**p* < 0.05, one-way ANOVA with Tukey’s post hoc test. See also Supplementary 5 for statistics.

Quantitative analysis of retrograde tracing revealed that midline thalamic nuclei provided the strongest input to most OFC subdivisions (Figure 15A). An exception was DLO, which received more widespread innervation across multiple thalamic nuclei. In addition to midline inputs, MO also received projections from medial thalamic nuclei. Visual inspection of the plots in Figure 15A suggests a high similarity between the thalamic innervation pattern of MO and AI on one side, and VO and LO on the other side. See also Supplementary 5 for statistics. In order to quantify the similarities of the patterns of thalamic input to the different OFC subdivisions, we calculated the correlation between the distributions of inputs for each pair of OFC subdivisions (Figure 15B), followed by hierarchical clustering (Figure 15C). The results suggest that VO and LO share a highly similar thalamic innervation pattern, and so do MO and AI.

Anterograde tracing analysis revealed distinct differences among OFC subdivisions in projections to thalamic nuclei (Supplementary Figure 2). The most medial MO and the most peripheral AI, each sent relatively sparse projections to thalamus. In contrast, central VO and LO provided dense reciprocal projections to midline thalamic nuclei, supporting strong OFC-thalamus interconnectivity. DLO again displayed broad projections, innervating multiple thalamic nuclei similarly to its widespread input projections pattern.

### 13. Intra-OFC connectivity graph suggests preferential directions of information flow

We next assessed whether the pattern of intra-OFC connectivity allowed any insights with respect to the flow of information within the structure.

We had access to two independent estimates of the directional connection strength between each pair of OFC subdivisions, labelled here as S_1_ and S_2_. On one hand, we measured the extent to which S_2_’s inputs (expressed as %input density) come from S_1_ relative to other subdivisions, with the retrograde injection localized in S_2_. We report this measurement for all pairs of subdivisions in Figure 16A. On the other hand, we also collected measurements from the datasets given in Alleńs mouse brain connectivity atlas (Atlas. 2011, Oh, Harris et al. 2014) of how much S_1_’s outputs targets S_2_ (expressed as %output density) relative to other subdivisions, investigated via anterograde injections in S_1_. We reproduce these measurements from the anterograde data for all pairs of OFC subdivisions in Figure 16B.

**Figure 16.**
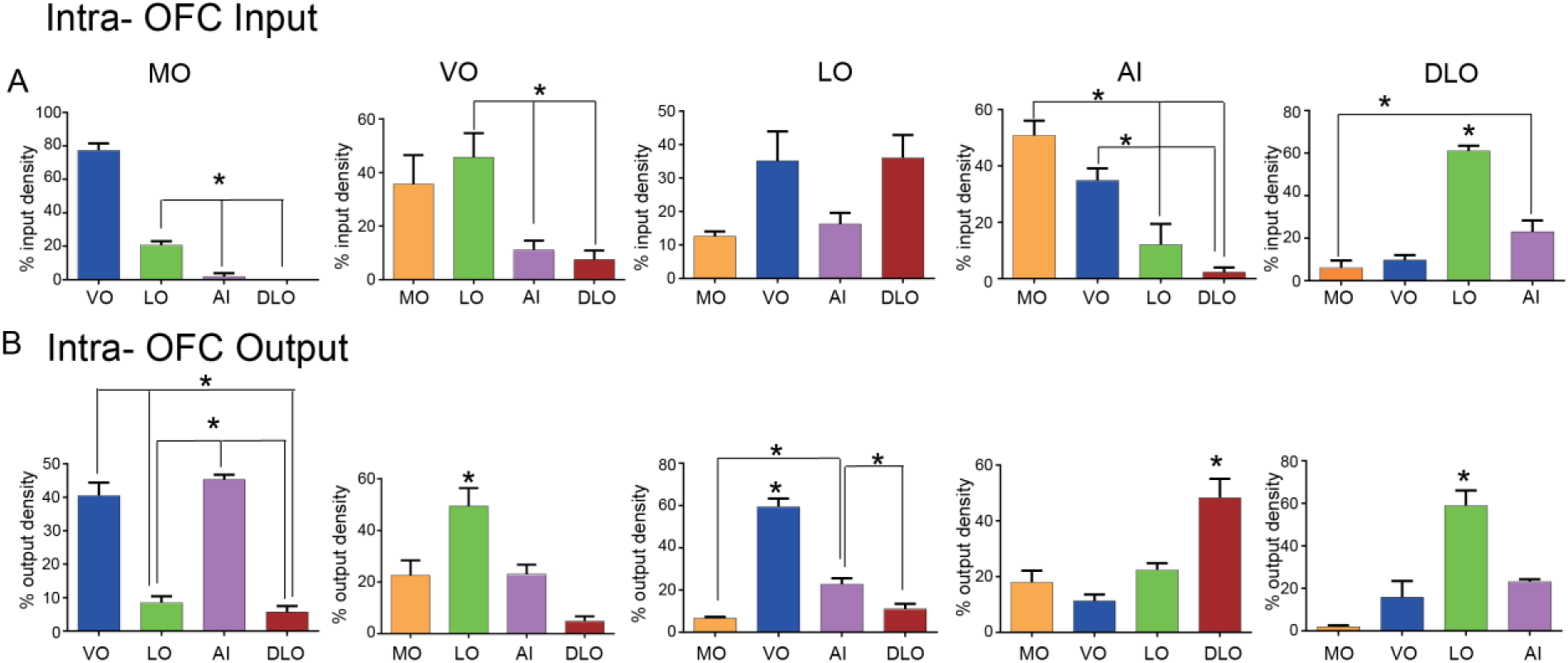
OFC intra-area connectivity. ***A)*** *Histograms show inputs from other OFC subdivisions to MO, VO, LO,, AI and DLO (left to right).. **B)** Corresponding output connectivity histograms of MO, VO, LO, AI and DLO (left to right) with other OFC subdivisions.* A single asterisk above a bar indicates the thalamic nucleus with significantly higher input than the other thalamic nuclei, \**p* < 0.05, one-way ANOVA with Tukey’s post hoc test. See also supplementary 5 for statistics.

For the same (S_1_,S_2_) pair, it is evident that the input-based and output-based descriptions of the S_1_→S_2_ connection are independent, because, for example, it is possible for S_1_ to project exclusively to S_2_ while S_2_ receives its inputs equally from all subdivisions, or for S_1_ to project equally over all subdivisions while S_2_ receives inputs only from S_1_. In each case, the output- or input-based description of S_1_→S_2_ will be equal to 100%, while the opposite description will be equal to 20% (because there are 5 subdivisions). Note, however, that both descriptions must agree on whether a connection exists or not, i.e., the input-based and output-based percentages must simultaneously be either both zero or both positive. Furthermore, the input- and output-based descriptions can be interpreted as simply the outcome of two distinct normalizations of the same matrix that contains the actual numerical strength (e. g. number of projecting axons) of each directional connection; the input-based (output-based) description is obtained by normalizing this matrix such that rows (columns) add up to 100%. With access to both descriptions, it ought to be possible to recover the original matrix of actual connection strengths (up to a multiplicative constant).

However, we note that the two matrices we report are in fact incompatible. Our measurements confirm the absence of a DLO→MO connection (0% of inputs to MO come from DLO), while the measurements reproduced from the Allen atlas anterograde data (Atlas. 2011, Oh, Harris et al. 2014) report that 1.47% of outputs from DLO target MO. The simultaneous presence of a zero and a nonzero estimate for the same connection preclude the recovery of the true connection strength matrix. We note, moreover, that manually zeroing this percentage in the output-based description followed by renormalizing still leads to a linear system with no solution, suggesting that further (more subtle) mathematical incompatibilities exist. In light of these findings, together with the observation that the retrograde labeling was characterized by more confined injections (cf. Figure 2) which allowed greater analytical precision, we elected to initially analyze the pattern of intra-OFC connectivity based exclusively on the input-based description.

Because connection percentages varied over orders of magnitude, we applied a simple thresholding procedure and excluded all connections weaker than the median observed percentage, and constructed a directed weighted graph based on the surviving connections, reported in Figure 17C. A separate attempt was made to recover the true connection strengths by repeatedly applying perturbations to the output percentages in order to obtain compatible matrices and recover estimated connection strengths, which we report in the Supplementary Figure 4, but the resulting connectivity structure and conclusions did not differ appreciably from the one obtained based exclusively on the inputs that we discuss below.

**Figure 17.**
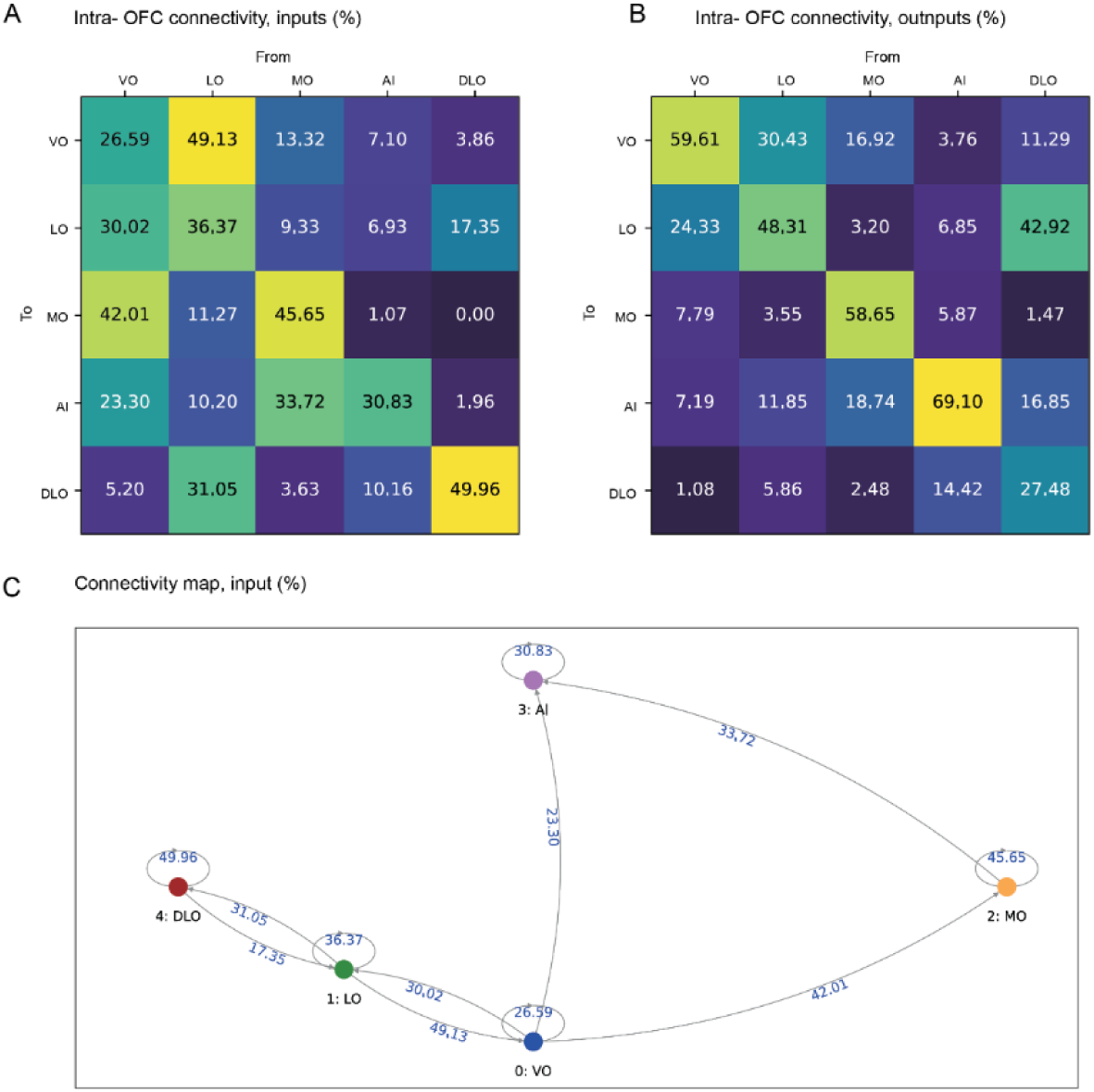
OFC intra-area connectivity and proposed preferential information flow. **A)** Representative distributions of *input* percentages from (columns) to (rows) all pairs of OFC subdivisions, as investigated by retrograde injections, and excluding inputs from the rest of the brain, which constitute a much smaller percentage compared to intra-OFC connections. **B)** Representative distributions of *output* percentages from (columns) to (rows) all pairs of OFC subdivisions, as investigated by anterograde injections, and excluding outputs to the rest of the brain, which constitute a much smaller percentage compared to intra-OFC connections. **C)** Representation of the matrix in A as a directed, weighted graph, after thresholding to exclude all connections with strength less than or equal to the median observed input percentage. Note how DLO and LO lack strong connections to and from the rest of the OFC subdivisions except via VO, and might function as an entry complex receiving inputs mostly from the rest of the brain, while conversely AI lacks intra-OFC outputs, suggesting it might function as an exit node sending outputs mostly to the rest of the brain. See supplementary materials for similar analysis taking into account the matrices in A and B simultaneously.

**Supplementary Figure 3. Related to Figure 17A).**
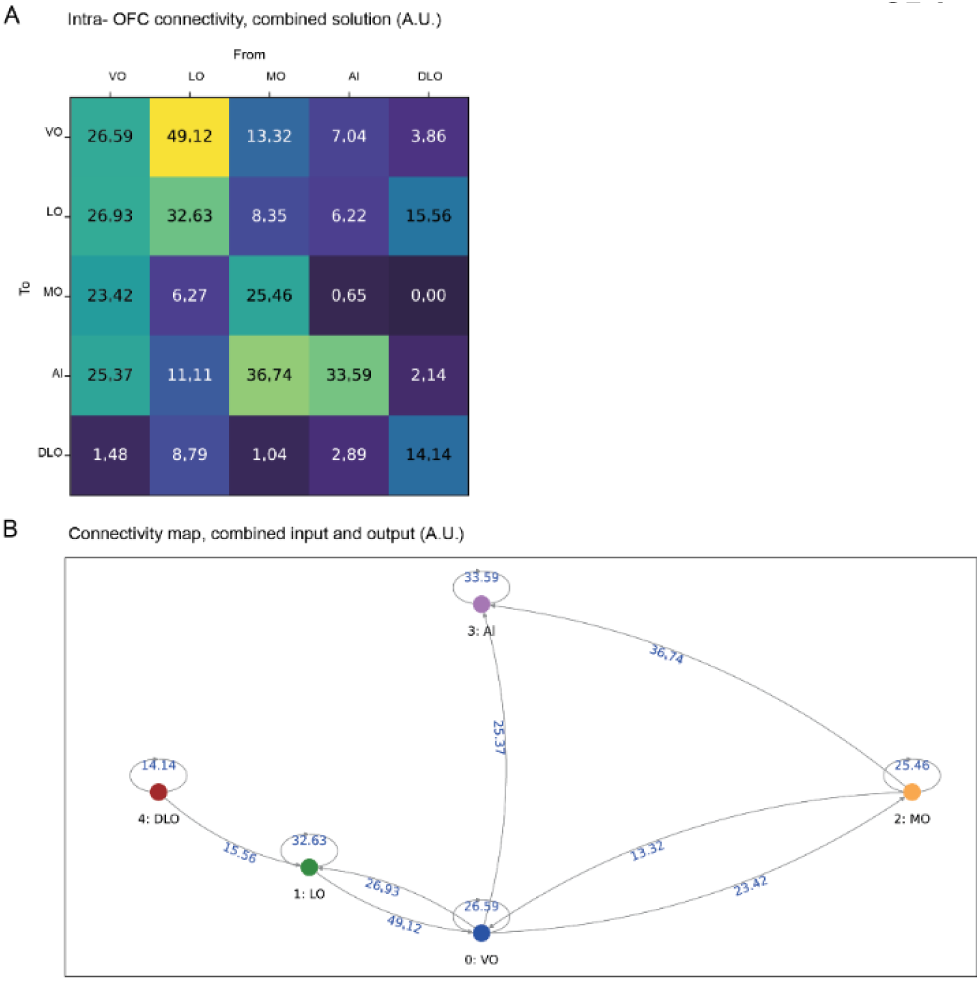
Reconstructed connection strengths obtained by repeatedly perturbing the elements of the output matrix (cf. methods, main text), combining with the original input matrix and solving the corresponding perturbed linear system. Average of 1000 iterations. **B)** Representation of the matrix in A as a directed, weighted graph, after thresholding to exclude all connections with strength less than or equal to the median reconstructed strength. Note the similarity to the graph depicted in Fig. 17C, with minor structural differences; the lack of a connection from LO to DLO emphasizes the latter’s function as an entry node receiving inputs mostly from the rest of the brain, while the presence of a reciprocal connection between MO and VO further emphasizes VO as a central integrative node, and isolates AI as an exit node sending outputs mostly to the rest of the brain.

Interestingly, the empirical topology shown in Figure 17C suggests that DLO and LO do not send or receive important intra-OFC connections to or from the other subdivisions except via VO, having instead the characteristics of an *entry node* or *entry complex* that is mostly driven by external inputs (recall that DLO, LO and VO share a similar pattern of inputs from the rest of the brain, Figure 13B,C). LO has important reciprocal connections with VO and DLO, but the estimated connection strengths suggest that the LO→VO and LO→DLO directions are more important than the opposite directions. Together, this suggests that LO→VO and DLO→LO→VO are potentially important input streams which, given the presence of potentially important shared inputs from Motor Cortex and Striatum to DLO and LO (cf. sections 10, 11), might be related to the animal’s behavioral state or motor plans. VO, on the other hand, has the largest number of intra-OFC targets among the subdivisions, as well as a unique profile of input and output connections to the rest of the brain which emphasizes thalamic, brainstem and prefrontal inputs (cf. sections 2, 10, 11), suggesting that it might play a key integrative role within the OFC. From VO, strong outputs reach both MO and AI, and AI in particular receives connections from VO and MO but does not send important intra-OFC outputs to any subdivision, having instead the characteristics of an *exit* node. In that sense, it is notable that, compared to other OFC subdivisions (cf. sections 10, 11), AI sends the most outputs to cognitive and limbic structures such as Amygdala and the Striatum, as well as (together with other subdivisions) Thalamus, Prefrontal Cortex, and Association Cortex, but without particularly emphasizing outputs to the Motor Cortex or Brainstem.

## DISCUSSION

We have quantified, compared and put in quantitative relation the normalized input densities, output densities and the densities of intra-area connections for the different OFC subdivisions in mouse OFC. Our results are synthesized in Figure 18A, which shows the hierarchical clustering of similarities among subdivisions according to the patterns of input, output, and by looking at their concatenation (Figure 18A, left, middle and right, respectively). VO and (D)LO share a similar, prominent thalamic input (see also Yang, Yang et al. (2025)), and cluster together with MO, but VO is set apart at output level, possibly for its strong efferent connectivity to association cortical areas. AI is positioned peripherally in both input and output dendrograms, suggesting that this subdivision has a distinct character from neighboring ones. On the other hand, AI is equally distinct from the remaining part of the insular cortex, according to Gehrlach, Weiand et al. (2020), where the anterior insular cortex (mostly agranular) clusters differently from the middle and posterior divisions. One could thus wonder whether it is more justified to incorporate AI in OFC (Price 2006, Zimmermann, Yamin et al. 2017) and not, as the name suggests, in the insular cortex. Indeed, recent input tracing work in mice did not include AI (Yang, Yang et al. 2025), while in rats some authors do include it (Izquierdo 2017) and others do not (Mathiasen, Aggleton et al. 2023). Our data suggest that it is legitimate to consider AI part of the mouse OFC circuit. First, the data of Figure 15 show the impressive similarity between the most medial and most lateral OFC subdivisions (MO and AI, respectively) in terms of thalamic innervation pattern, a feature often used as a criterion for areal identity (Iurilli, Ghezzi et al. 2012, Miller-Hansen and Sherman 2022, Howell, Warrington et al. 2025). Second, MO and AI are strongly interconnected, as AI is the main target of MO (in the efferent series analysis) and MO is the main input to AI (in the afferent series analysis). Our computation of the information flow within OFC also reaffirms a prominent connectivity from MO to AI, while suggesting that whereas (D)LO function mostly as an entry point for information within OFC, AI has the characteristics of an output node of OFC.

**Figure 18:**
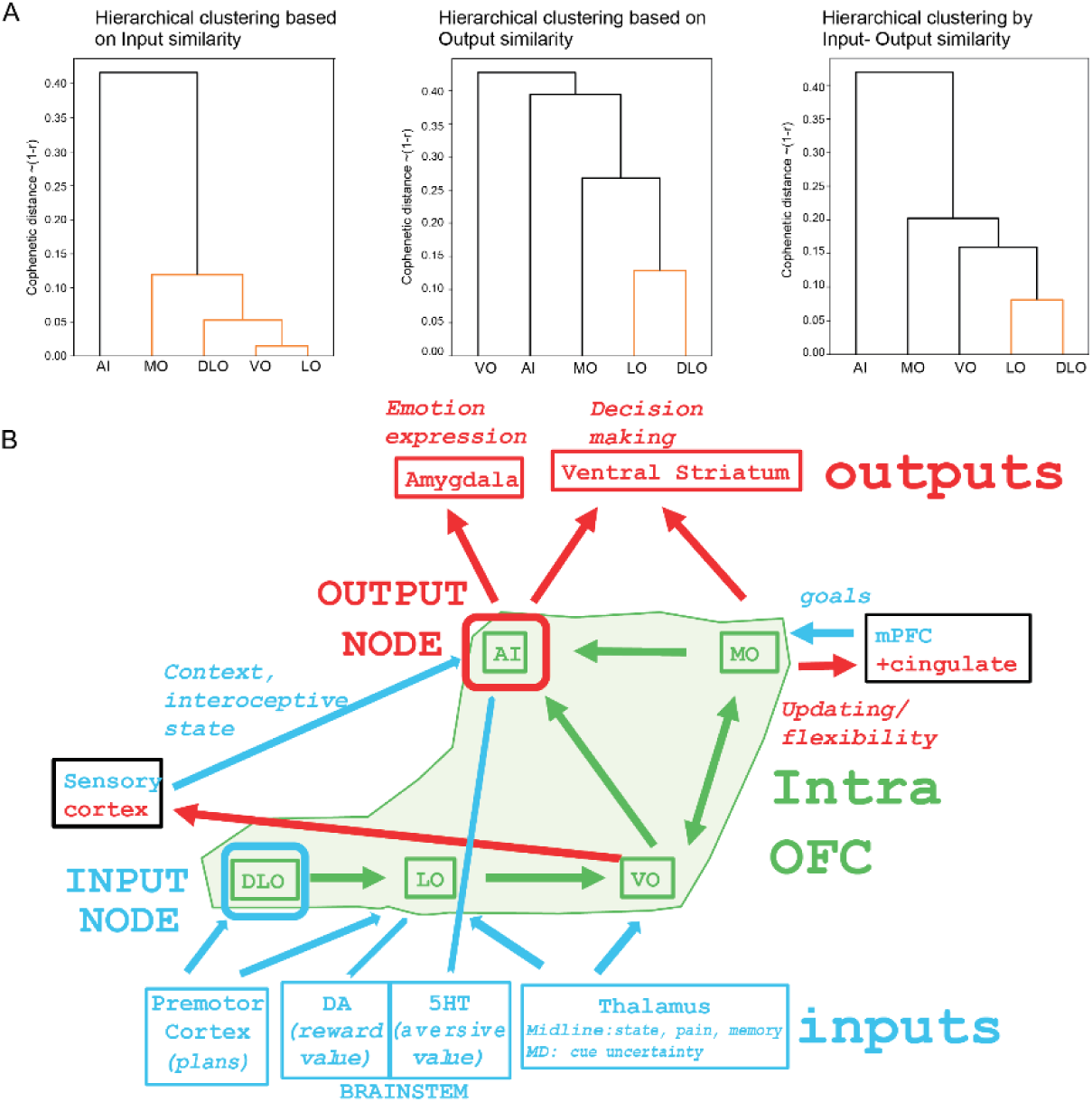
Clustering results and proposed input integration model within OFC necessary for the output target activation. **A)** Cluster dendrograms for inputs, outputs, and concatenated inputs and outputs. **B)** Proposed anatomo-functional stream where information flows based on the combination of OFC subdivisionś inputs, outputs and intra-area connectivity. Information on the animal motor plan is conveyed via DLO (input node) and LO, that also integrate information on the stimulus reward value (brainstem dopaminergic modulation). Thalamic inputs on animal state and memories as well as cue uncertainty enters mainly in LO and VO. Area VO sends mainly information to MO, highly interconnected with mPFC and cingulate, thus integrating and updating animaĺs current goals. AI, that receives dominant inputs from MO, has characteristics of an output node innervating mainly the amygdala, responsible for emotional expression, and the ventral striatum, responsible for stimulus value-related decision making.

Our data on intra-OFC connectivity allows the proposal of a sketch of the putative information flow within OFC. This model foresees a flow from (D)LO (entry complex of the circuit) to VO and from there to MO, with MO finally projecting to AI. This view, together with our data about the main external inputs to the various OFC subdivisions, allows us to propose a model in which the intra-OFC circuits gradually integrate information about the animal motor plans (premotor cortex inputs to (D)LO), the reward value of stimulus contingencies (brainstem dopaminergic inputs to LO), and various types of thalamic inputs on the animal status, also in relation to stimulus features (to LO/VO) – see Figure 18B. More specifically, midline thalamus conveys information on nociceptive stimuli (Chen 2009) and memories (given the prominent projections from hippocampus (Griffin 2015), whereas the medio-dorsal conveys information about stimulus cue uncertainty (Mukherjee, Lam et al. 2021). At MO, information on the animaĺs goals from medial prefrontal cortex (Yang, Hu et al. 2022) is also integrated, and the output from MO to mPFC might be well needed to update the current goal setting. MO circuitry is indeed reciprocally interlinked to the medial prefrontal cortex according to ours and other recent data (Yang, Yang et al. 2025). Within OFC, the main target of MO is AI, which has comparatively unimportant intra-OFC outputs. Note that among the main target structures of AI are the amygdala and the ventral striatum, critical structures for emotional reactivity and reward-driven, motivated decision making.

OFC receives multiple sensory inputs from a variety of cortical association areas (Figure 8A) (Sharma and Bandyopadhyay 2020, Tripathi, Sato et al. 2021), where visual inputs tends to be more dominant in the medial subdivisions (MO and VO), whereas the integration between acoustic and chemical senses tends to be dominant in AI. The latter observation is relevant as the integration of acoustic inputs and odours from pups is enhanced in auditory cortex after pregnancy (Cohen, Rothschild et al. 2011), and that this input is transferred to OFC via connections very sensitive to key neuromodulators after pregnancy such as neurosteroids (Tripathi, Sato et al. 2021). Recent observation documented OFC neuronal response related to anticipatory pup-oriented retrieval movements, responsible for brainstem dopaminergic activation (Tasaka, Hagihara et al. 2025), possibly via the marked connections from LO to the ventral tegmental area (see our Figure 5E). It is not a surprise that gustatory inputs, largely from posterior insula (Livneh, Sugden et al. 2020) are further processed within OFC. The fundamental works of Rolls lab (Rolls 2019) demonstrating value encoding in primate OFC was indeed performed using food as stimuli, and showed that whereas insular neurons respond to food irrespective of the animal satiety, OFC neurons respond to food reward only when hungry (Rolls 2019), and malfunction of OFC inhibitory neurons is involved in disturbances of eating behavior (Li, Chen et al. 2024). Indeed, reduced response of OFC inhibitory interneurons to rewarding stimuli such as food accounts for the increased responsiveness and hence the lack of devaluation to food stimuli in obese mice (Seabrook, Naef et al. 2023). In relation to OFC role in encoding stimulus reward salience it is important to remember that the main OFC output nodes (AI) project preferentially to the ventral striatum, critical for motivation-driven behavior (Enriquez-Traba, Arenivar et al. 2025). Communication between the OFC, ventral striatum and pallidum would likely be rapid as our work describes a relatively new finding of direct pallidal and striatal connections to the cortex. This establishes a monosynaptic reciprocal loop between the OFC and striatopallidal basal ganglia system. A few other studies have begun to investigate such direct pallidal input connectivity to the cortex (Saunders, Oldenburg et al. 2015, Ferenczi, Wang et al. 2025). However, to the best of our knowledge, we report for the first time the existence of topographically organized, direct, striato-cortical projections, whose physiological role will have to be further investigated in the near future.

One of the motivations for this study was also the fundamental observation by Rolls and colleagues that the medial and lateral OFC preferentially process rewarding and aversive stimuli, respectively (Rolls, Cheng et al. 2020). Importantly, this functional segregation is disturbed in major depression so as to favour processing of aversive stimuli, with profound implications for our understanding and for designing potential new therapeutic approaches for mood disorders (Zhang, Rolls et al. 2024). The main reservation for extending these observations to rodents is given by the two fundamental facts that reward coding is to a significant extent happening already at subcortical level in rodents on the one hand (Rolls 2019), and by the fact that rodent OFC encodes not only stimulus value, but also action value (reviewed in Rolls, Cheng et al. (2020)). No clear evidence for such a functional segregation can be found in the existing literature, even if the topic has not been systematically investigated. For example, processing of aversive gustatory stimuli does not seem to be affected by functional inactivation confined to lateral OFC (Panayi and Killcross 2018). However, we found that brainstem dopaminergic innervation – potentially conveying reward-related aspects of stimuli - is significantly more pronounced in the more medial VO, compared to serotoninergic innervation – related to aversive stimuli -, the latter being instead dominant in the more lateral AI. These observations should prompt studies investigating more systematically the question of the phylogenetically conserved spatial segregation of rewarding vs aversive stimuli within OFC in rodents.

In conclusion, our work is more in line with an intra-OFC serial integration of the information necessary to update goals, regulate emotional state and drive motivational behaviour, rather than pointing to the existence of parallel, distinct information streams (a scheme that may have become more dominant in primates (Ben Shalom 2024). This is in line with decision-making, the ultimate function of OFC (together with the corresponding striato-thalamic circuit), requiring more serial than parallel information processing and integration. Future studies will investigate the functional significance of the intra-OFC paths we identified, as well as of the prominent loops emanating from VO/LO (with thalamus), MO (with mPFC) and AI (with amygdala), possibly via pathway specific, viral based manipulation of identified connections (Tripathi, Sato et al. 2021).

## REFERENCES

Atlas., A. M. B. (2011). Available from mouse.brain-map.org.

Barreiros, I. V., H. Ishii, M. E. Walton and M. C. Panayi (2021). "Defining an orbitofrontal compass: Functional and anatomical heterogeneity across anterior-posterior and medial-lateral axes." Behav Neurosci 135(2): 165–173.

Barreiros, I. V., M. C. Panayi and M. E. Walton (2021). "Organization of Afferents along the Anterior-posterior and Medial-lateral Axes of the Rat Orbitofrontal Cortex." Neuroscience 460: 53–68.

Ben Shalom, D. (2024). "Editorial: The four streams of the prefrontal cortex." Front Neuroanat 18: 1487947.

Bjerke, I., M. Ovsthus, M. Checinska and T. Leergaard (2023). An illustrated guide to landmarks in histological rat and mouse brain images, Zenodo.

Cazares, C., D. C. Schreiner, M. L. Valencia and C. M. Gremel (2022). "Orbitofrontal cortex populations are differentially recruited to support actions." Curr Biol 32(21): 4675–4687 e4675.

Chen, A. C. (2009). "Higher cortical modulation of pain perception in the human brain: Psychological determinant." Neurosci Bull 25(5): 267–276.

Cohen, L., G. Rothschild and A. Mizrahi (2011). "Multisensory integration of natural odors and sounds in the auditory cortex." Neuron 72(2): 357–369.

Enriquez-Traba, J., M. Arenivar, H. E. Yarur-Castillo, C. Noh, R. J. Flores, T. Weil, S. Roy, T. B. Usdin, C. T. LaGamma, H. Wang, V. S. Tsai, D. Kerspern, A. E. Moritz, D. R. Sibley, A. Lutas, R. Moratalla, Z. Freyberg and H. A. Tejeda (2025). "Dissociable control of motivation and reinforcement by distinct ventral striatal dopamine receptors." Nat Neurosci 28(1): 105–121.

Ferenczi, E. A., W. Wang, A. Biswas, T. Pottala, Y. Dong, A. K. Chan, M. A. Albanese, R. S. Sohur, T. Jia, K. J. Mastro and B. L. Sabatini (2025). "Reciprocal projections between the globus pallidus externa and cortex span motor and nonmotor regions." Proc Natl Acad Sci U S A 122(23): e2423367122.

Furuyashiki, T. and M. Gallagher (2007). "Neural encoding in the orbitofrontal cortex related to goal-directed behavior." Ann N Y Acad Sci 1121: 193–215.

Gehrlach, D. A., C. Weiand, T. N. Gaitanos, E. Cho, A. S. Klein, A. A. Hennrich, K. K. Conzelmann and N. Gogolla (2020). "A whole-brain connectivity map of mouse insular cortex." Elife 9.

Gourley, S. L., K. S. Zimmermann, A. G. Allen and J. R. Taylor (2016). "The Medial Orbitofrontal Cortex Regulates Sensitivity to Outcome Value." J Neurosci 36(16): 4600–4613.

Griffin, A. L. (2015). "Role of the thalamic nucleus reuniens in mediating interactions between the hippocampus and medial prefrontal cortex during spatial working memory." Front Syst Neurosci 9: 29.

Howell, A. M., S. Warrington, C. Fonteneau, Y. T. Cho, S. N. Sotiropoulos, J. D. Murray and A. Anticevic (2025). "The spatial extent of anatomical connections within the thalamus varies across the cortical hierarchy in humans and macaques." bioRxiv.

Iurilli, G., D. Ghezzi, U. Olcese, G. Lassi, C. Nazzaro, R. Tonini, V. Tucci, F. Benfenati and P. Medini (2012). "Sound-driven synaptic inhibition in primary visual cortex." Neuron 73(4): 814–828.

Izquierdo, A. (2017). "Functional Heterogeneity within Rat Orbitofrontal Cortex in Reward Learning and Decision Making." J Neurosci 37(44): 10529–10540.

Krettek, J. E. and J. L. Price (1977). "The cortical projections of the mediodorsal nucleus and adjacent thalamic nuclei in the rat." J Comp Neurol 171(2): 157–191.

Li, W., X. Chen, Y. Luo, M. Xiao, Y. Liu and H. Chen (2024). "Altered connectivity patterns of medial and lateral orbitofrontal cortex underlie the severity of bulimic symptoms." Int J Clin Health Psychol 24(1): 100439.

Livneh, Y., A. U. Sugden, J. C. Madara, R. A. Essner, V. I. Flores, L. A. Sugden, J. M. Resch, B. B. Lowell and M. L. Andermann (2020). "Estimation of Current and Future Physiological States in Insular Cortex." Neuron 105(6): 1094–1111 e1010.

Mathiasen, M. L., J. P. Aggleton and M. P. Witter (2023). "Projections of the insular cortex to orbitofrontal and medial prefrontal cortex: A tracing study in the rat." Front Neuroanat 17: 1131167.

Miller-Hansen, A. J. and S. M. Sherman (2022). "Conserved patterns of functional organization between cortex and thalamus in mice." Proc Natl Acad Sci U S A 119(21): e2201481119.

Mukherjee, A., N. H. Lam, R. D. Wimmer and M. M. Halassa (2021). "Thalamic circuits for independent control of prefrontal signal and noise." Nature 600(7887): 100–104.

Oh, S. W., J. A. Harris, L. Ng, B. Winslow, N. Cain, S. Mihalas, Q. Wang, C. Lau, L. Kuan, A. M. Henry, M. T. Mortrud, B. Ouellette, T. N. Nguyen, S. A. Sorensen, C. R. Slaughterbeck, W. Wakeman, Y. Li, D. Feng, A. Ho, E. Nicholas, K. E. Hirokawa, P. Bohn, K. M. Joines, H. Peng, M. J. Hawrylycz, J. W. Phillips, J. G. Hohmann, P. Wohnoutka, C. R. Gerfen, C. Koch, A. Bernard, C. Dang, A. R. Jones and H. Zeng (2014). "A mesoscale connectome of the mouse brain." Nature 508(7495): 207–214.

Panayi, M. C. and S. Killcross (2018). "Functional heterogeneity within the rodent lateral orbitofrontal cortex dissociates outcome devaluation and reversal learning deficits." Elife 7.

Paxinos, G. and K. Franklin (2001). The Mouse Brain in Stereotaxic Coordinates. USA, Academic Press.

Price, J. L. (2006). Architectonic structure of the orbital and medial prefrontal cortex. The Orbitofrontal Cortex, Oxford University Press: 3-18.

Price, J. L. (2007). "Definition of the orbital cortex in relation to specific connections with limbic and visceral structures and other cortical regions." Ann N Y Acad Sci 1121: 54–71.

Ray, J. P. and J. L. Price (1992). "The organization of the thalamocortical connections of the mediodorsal thalamic nucleus in the rat, related to the ventral forebrain-prefrontal cortex topography." J Comp Neurol 323(2): 167–197.

Rolls, E. T. (2019). The Rodent Orbitofrontal Cortex. The Orbitofrontal Cortex, Oxford Academic: 228–236.

Rolls, E. T., W. Cheng and J. Feng (2020). "The orbitofrontal cortex: reward, emotion and depression." Brain Commun 2(2): fcaa196.

Rudebeck, P. H. and E. L. Rich (2018). "Orbitofrontal cortex." Curr Biol 28(18): R1083–R1088.

Seabrook, L. T., L. Naef, C. Baimel, A. K. Judge, T. Kenney, M. Ellis, T. Tayyab, M. Armstrong, M. Qiao, S. B. Floresco and S. L. Borgland (2023). "Disinhibition of the orbitofrontal cortex biases decision-making in obesity." Nat Neurosci 26(1): 92–106.

Sergejeva, M., E. A. Papp, R. Bakker, M. A. Gaudnek, Y. Okamura-Oho, J. Boline, J. G. Bjaalie and A. Hess (2015). "Anatomical landmarks for registration of experimental image data to volumetric rodent brain atlasing templates." J Neurosci Methods 240: 161–169.

Sharma, S. and S. Bandyopadhyay (2020). "Differential Rapid Plasticity in Auditory and Visual Responses in the Primarily Multisensory Orbitofrontal Cortex." eNeuro 7(3).

Tasaka, G. I., M. Hagihara, S. Irie, H. Kobayashi, K. Inada, K. Kobayashi, S. Kato, K. Kobayashi and K. Miyamichi (2025). "Orbitofrontal cortex influences dopamine dynamics associated with alloparental behavioral acquisition in female mice." Sci Adv 11(27): eadr4620.

Tripathi, A., S. S. Sato and P. Medini (2021). "Cortico-cortical connectivity behind acoustic information transfer to mouse orbitofrontal cortex is sensitive to neuromodulation and displays local sensory gating: relevance in disorders with auditory hallucinations?" J Psychiatry Neurosci 46(3): E371–E387.

Ward, R. D., V. Winiger, E. R. Kandel, P. D. Balsam and E. H. Simpson (2015). "Orbitofrontal cortex mediates the differential impact of signaled-reward probability on discrimination accuracy." Front Neurosci 9: 230.

Yang, C., Y. Hu, A. D. Talishinsky, C. T. Potter, C. B. Calva, L. A. Ramsey, A. J. Kesner, R. F. Don, S. Junn, A. Tan, A. F. Pierce, C. Nicolas, Y. Arima, S. C. Lee, C. Su, J. M. Coudriet, C. A. Mejia-Aponte, D. V. Wang, H. Lu, Y. Yang and S. Ikemoto (2022). "Medial prefrontal cortex and anteromedial thalamus interaction regulates goal-directed behavior and dopaminergic neuron activity." Nat Commun 13(1): 1386.

Yang, M., H. Yang, L. Shen and T. Xu (2025). "Anatomical mapping of whole-brain monosynaptic inputs to the orbitofrontal cortex." Front Neural Circuits 19: 1567036.

Zhang, B., E. T. Rolls, X. Wang, C. Xie, W. Cheng and J. Feng (2024). "Roles of the medial and lateral orbitofrontal cortex in major depression and its treatment." Mol Psychiatry 29(4): 914–928.

Zimmermann, K. S., J. A. Yamin, D. G. Rainnie, K. J. Ressler and S. L. Gourley (2017). "Connections of the Mouse Orbitofrontal Cortex and Regulation of Goal-Directed Action Selection by Brain-Derived Neurotrophic Factor." Biol Psychiatry 81(4): 366–377.

Banerjee, A., G. Parente, J. Teutsch, C. Lewis, F. F. Voigt and F. Helmchen (2020). "Value-guided remapping of sensory cortex by lateral orbitofrontal cortex." Nature 585(7824): 245–250.

Saunders, A., I. A. Oldenburg, V. K. Berezovskii, C. A. Johnson, N. D. Kingery, H. L. Elliott, T. Xie, C. R. Gerfen and B. L. Sabatini (2015). "A direct GABAergic output from the basal ganglia to frontal cortex." Nature 521(7550): 85–89.

