## Supplementary 5 for "Anatomical input-output streams within mouse orbitofrontal cortex subdivisions"

**Projection from OFC subdivision to Thalamic nuclei**

**Table Analyzed** **MO**

**ANOVA summary**

| | |
|---|---|
| F | 26,19 |
| P value | 0,0005 |
| P value summary | *** |
| Are differences among means statistic | Yes |
| R square | 0,9562 |

**Tukey's multiple comparisons test**

| | Mean 1 | Mean 2 | Mean Diff, | SE of diff, | n1 | n2 | q | DF | Significant | Summary | Adjusted P Value |
|---|---|---|---|---|---|---|---|---|---|---|---|
| Medial vs. RT(Inhibitory) | 9,445 | 1,081 | 8,364 | 1,655 | 2 | 2 | 7,146 | 6 | Yes | * | 0,017 |
| Medial vs. ventral | 9,445 | 0,7792 | 8,666 | 1,655 | 2 | 2 | 7,404 | 6 | Yes | * | 0,0144 |
| Medial vs. anterior | 9,445 | 3,76 | 5,685 | 1,655 | 2 | 2 | 4,858 | 6 | No | ns | 0,0902 |
| Medial vs. midline | 9,445 | 16,55 | -7,104 | 1,655 | 2 | 2 | 6,069 | 6 | Yes | * | 0,0361 |
| Medial vs. intralaminar | 9,445 | 5,766 | 3,678 | 1,655 | 2 | 2 | 3,143 | 6 | No | ns | 0,3456 |
| RT(Inhibitory) vs. ventral | 1,081 | 0,7792 | 0,3018 | 1,655 | 2 | 2 | 0,2579 | 6 | No | ns | > 0,9999 |
| RT(Inhibitory) vs. anterior | 1,081 | 3,76 | -2,679 | 1,655 | 2 | 2 | 2,289 | 6 | No | ns | 0,6167 |
| RT(Inhibitory) vs. midline | 1,081 | 16,55 | -15,47 | 1,655 | 2 | 2 | 13,22 | 6 | Yes | *** | 0,0007 |
| RT(Inhibitory) vs. intralaminar | 1,081 | 5,766 | -4,685 | 1,655 | 2 | 2 | 4,003 | 6 | No | ns | 0,1772 |
| ventral vs. anterior | 0,7792 | 3,76 | -2,98 | 1,655 | 2 | 2 | 2,547 | 6 | No | ns | 0,5258 |
| ventral vs. midline | 0,7792 | 16,55 | -15,77 | 1,655 | 2 | 2 | 13,47 | 6 | Yes | *** | 0,0006 |
| ventral vs. intralaminar | 0,7792 | 5,766 | -4,987 | 1,655 | 2 | 2 | 4,261 | 6 | No | ns | 0,1444 |
| anterior vs. midline | 3,76 | 16,55 | -12,79 | 1,655 | 2 | 2 | 10,93 | 6 | Yes | ** | 0,0019 |
| anterior vs. intralaminar | 3,76 | 5,766 | -2,007 | 1,655 | 2 | 2 | 1,715 | 6 | No | ns | 0,8179 |
| midline vs. intralaminar | 16,55 | 5,766 | 10,78 | 1,655 | 2 | 2 | 9,212 | 6 | Yes | ** | 0,0048 |

**Table Analyzed** **VO**

**ANOVA summary**

| | |
|---|---|
| F | 7,144 |
| P value | 0,0008 |
| P value summary | *** |
| Are differences among means statistic | Yes |
| R square | 0,6649 |

**Tukey's multiple comparisons test**

| | Mean 1 | Mean 2 | Mean Diff, | SE of diff, | n1 | n2 | q | DF | Significant | Summary | Adjusted P Value |
|---|---|---|---|---|---|---|---|---|---|---|---|
| Medial vs. RT(Inhibitory) | 8,712 | 3,609 | 5,103 | 2,732 | 4 | 4 | 2,642 | 18 | No | ns | 0,451 |
| Medial vs. ventral | 8,712 | 6,882 | 1,83 | 2,732 | 4 | 4 | 0,9473 | 18 | No | ns | 0,9831 |
| Medial vs. anterior | 8,712 | 7,183 | 1,529 | 2,732 | 4 | 4 | 0,7915 | 18 | No | ns | 0,9925 |
| Medial vs. midline | 8,712 | 18,72 | -10 | 2,732 | 4 | 4 | 5,18 | 18 | Yes | * | 0,0187 |
| Medial vs. intralaminar | 8,712 | 7,498 | 1,214 | 2,732 | 4 | 4 | 0,6284 | 18 | No | ns | 0,9974 |
| RT(Inhibitory) vs. ventral | 3,609 | 6,882 | -3,273 | 2,732 | 4 | 4 | 1,694 | 18 | No | ns | 0,8321 |
| RT(Inhibitory) vs. anterior | 3,609 | 7,183 | -3,574 | 2,732 | 4 | 4 | 1,85 | 18 | No | ns | 0,7769 |
| RT(Inhibitory) vs. midline | 3,609 | 18,72 | -15,11 | 2,732 | 4 | 4 | 7,821 | 18 | Yes | *** | 0,0004 |
| RT(Inhibitory) vs. intralaminar | 3,609 | 7,498 | -3,889 | 2,732 | 4 | 4 | 2,013 | 18 | No | ns | 0,7131 |
| ventral vs. anterior | 6,882 | 7,183 | -0,3011 | 2,732 | 4 | 4 | 0,1559 | 18 | No | ns | > 0,9999 |
| ventral vs. midline | 6,882 | 18,72 | -11,83 | 2,732 | 4 | 4 | 6,127 | 18 | Yes | ** | 0,0046 |
| ventral vs. intralaminar | 6,882 | 7,498 | -0,616 | 2,732 | 4 | 4 | 0,3189 | 18 | No | ns | > 0,9999 |
| anterior vs. midline | 7,183 | 18,72 | -11,53 | 2,732 | 4 | 4 | 5,971 | 18 | Yes | ** | 0,0058 |
| anterior vs. intralaminar | 7,183 | 7,498 | -0,3149 | 2,732 | 4 | 4 | 0,163 | 18 | No | ns | > 0,9999 |
| midline vs. intralaminar | 18,72 | 7,498 | 11,22 | 2,732 | 4 | 4 | 5,808 | 18 | Yes | ** | 0,0074 |

**Table Analyzed** **LO**

**ANOVA summary**

| | |
|---|---|
| F | 15,98 |
| P value | < 0,0001 |
| P value summary | **** |
| Are differences among means statistic | Yes |
| R square | 0,8694 |

**Tukey's multiple comparisons test**

| | Mean 1 | Mean 2 | Mean Diff, | SE of diff, | n1 | n2 | q | DF | Significant | Summary | Adjusted P Value |
|---|---|---|---|---|---|---|---|---|---|---|---|
| Medial vs. RT(Inhibitory) | 7,293 | 0,7601 | 6,533 | 2,745 | 3 | 3 | 3,365 | 12 | No | ns | 0,2369 |
| Medial vs. ventral | 7,293 | 3,695 | 3,598 | 2,745 | 3 | 3 | 1,854 | 12 | No | ns | 0,7744 |
| Medial vs. anterior | 7,293 | 7,672 | -0,3792 | 2,745 | 3 | 3 | 0,1954 | 12 | No | ns | > 0,9999 |
| Medial vs. midline | 7,293 | 22,88 | -15,58 | 2,745 | 3 | 3 | 8,028 | 12 | Yes | ** | 0,0011 |
| Medial vs. intralaminar | 7,293 | 5,139 | 2,154 | 2,745 | 3 | 3 | 1,109 | 12 | No | ns | 0,965 |
| RT(Inhibitory) vs. ventral | 0,7601 | 3,695 | -2,935 | 2,745 | 3 | 3 | 1,512 | 12 | No | ns | 0,8845 |

| | | | | | | | | | | | |
|---|---|---|---|---|---|---|---|---|---|---|---|
| RT(Inhibitory) vs. anterior | 0,7601 | 7,672 | -6,912 | 2,745 | 3 | 3 | 3,561 | 12 | No | ns | 0,1932 |
| RT(Inhibitory) vs. midline | 0,7601 | 22,88 | -22,12 | 2,745 | 3 | 3 | 11,39 | 12 | Yes | **** | < 0,0001 |
| RT(Inhibitory) vs. intralaminar | 0,7601 | 5,139 | -4,379 | 2,745 | 3 | 3 | 2,256 | 12 | No | ns | 0,6162 |
| ventral vs. anterior | 3,695 | 7,672 | -3,978 | 2,745 | 3 | 3 | 2,049 | 12 | No | ns | 0,6996 |
| ventral vs. midline | 3,695 | 22,88 | -19,18 | 2,745 | 3 | 3 | 9,882 | 12 | Yes | *** | 0,0002 |
| ventral vs. intralaminar | 3,695 | 5,139 | -1,445 | 2,745 | 3 | 3 | 0,7443 | 12 | No | ns | 0,9939 |
| anterior vs. midline | 7,672 | 22,88 | -15,21 | 2,745 | 3 | 3 | 7,833 | 12 | Yes | ** | 0,0014 |
| anterior vs. intralaminar | 7,672 | 5,139 | 2,533 | 2,745 | 3 | 3 | 1,305 | 12 | No | ns | 0,9331 |
| midline vs. intralaminar | 22,88 | 5,139 | 17,74 | 2,745 | 3 | 3 | 9,138 | 12 | Yes | *** | 0,0003 |

**Table Analyzed** **AI**

**ANOVA summary**

| | |
|---|---|
| F | 3,963 |
| P value | 0,0235 |
| P value summary | * |
| Are differences among means statistic | Yes |
| R square | 0,6228 |

**Tukey's multiple comparisons test**

| | Mean 1 | Mean 2 | Mean Diff, | SE of diff, | n1 | n2 | q | DF | Significant | Summary | Adjusted P Value |
|---|---|---|---|---|---|---|---|---|---|---|---|
| Medial vs. RT(Inhibitory) | 5,119 | 0,4703 | 4,649 | 3,022 | 3 | 3 | 2,175 | 12 | No | ns | 0,649 |
| Medial vs. ventral | 5,119 | 0,9021 | 4,217 | 3,022 | 3 | 3 | 1,973 | 12 | No | ns | 0,7293 |
| Medial vs. anterior | 5,119 | 0,06641 | 5,053 | 3,022 | 3 | 3 | 2,364 | 12 | No | ns | 0,5723 |
| Medial vs. midline | 5,119 | 10,76 | -5,637 | 3,022 | 3 | 3 | 2,638 | 12 | No | ns | 0,4648 |
| Medial vs. intralaminar | 5,119 | 0,458 | 4,661 | 3,022 | 3 | 3 | 2,181 | 12 | No | ns | 0,6467 |
| RT(Inhibitory) vs. ventral | 0,4703 | 0,9021 | -0,4318 | 3,022 | 3 | 3 | 0,2021 | 12 | No | ns | > 0,9999 |
| RT(Inhibitory) vs. anterior | 0,4703 | 0,06641 | 0,4038 | 3,022 | 3 | 3 | 0,189 | 12 | No | ns | > 0,9999 |
| RT(Inhibitory) vs. midline | 0,4703 | 10,76 | -10,29 | 3,022 | 3 | 3 | 4,813 | 12 | Yes | * | 0,0464 |
| RT(Inhibitory) vs. intralaminar | 0,4703 | 0,458 | 0,01225 | 3,022 | 3 | 3 | 0,005732 | 12 | No | ns | > 0,9999 |
| ventral vs. anterior | 0,9021 | 0,06641 | 0,8357 | 3,022 | 3 | 3 | 0,391 | 12 | No | ns | 0,9997 |
| ventral vs. midline | 0,9021 | 10,76 | -9,854 | 3,022 | 3 | 3 | 4,611 | 12 | No | ns | 0,059 |
| ventral vs. intralaminar | 0,9021 | 0,458 | 0,4441 | 3,022 | 3 | 3 | 0,2078 | 12 | No | ns | > 0,9999 |
| anterior vs. midline | 0,06641 | 10,76 | -10,69 | 3,022 | 3 | 3 | 5,002 | 12 | Yes | * | 0,0371 |
| anterior vs. intralaminar | 0,06641 | 0,458 | -0,3916 | 3,022 | 3 | 3 | 0,1832 | 12 | No | ns | > 0,9999 |
| midline vs. intralaminar | 10,76 | 0,458 | 10,3 | 3,022 | 3 | 3 | 4,818 | 12 | Yes | * | 0,0461 |

**Table Analyzed** **DLO**

**ANOVA summary**

| | |
|---|---|
| F | 1,16 |
| P value | 0,383 |
| P value summary | ns |
| Are differences among means statistic | No |
| R square | 0,3258 |

**Tukey's multiple comparisons test**

| | Mean 1 | Mean 2 | Mean Diff, | SE of diff, | n1 | n2 | q | DF | Significant | Summary | Adjusted P Value |
|---|---|---|---|---|---|---|---|---|---|---|---|
| Medial vs. RT(Inhibitory) | 10,17 | 0,5724 | 9,598 | 4,945 | 3 | 3 | 2,745 | 12 | No | ns | 0,425 |
| Medial vs. ventral | 10,17 | 3,056 | 7,114 | 4,945 | 3 | 3 | 2,035 | 12 | No | ns | 0,7053 |
| Medial vs. anterior | 10,17 | 9,028 | 1,142 | 4,945 | 3 | 3 | 0,3266 | 12 | No | ns | 0,9999 |
| Medial vs. midline | 10,17 | 7,75 | 2,421 | 4,945 | 3 | 3 | 0,6923 | 12 | No | ns | 0,9957 |
| Medial vs. intralaminar | 10,17 | 4,154 | 6,016 | 4,945 | 3 | 3 | 1,721 | 12 | No | ns | 0,821 |
| RT(Inhibitory) vs. ventral | 0,5724 | 3,056 | -2,484 | 4,945 | 3 | 3 | 0,7104 | 12 | No | ns | 0,9951 |
| RT(Inhibitory) vs. anterior | 0,5724 | 9,028 | -8,456 | 4,945 | 3 | 3 | 2,418 | 12 | No | ns | 0,5504 |
| RT(Inhibitory) vs. midline | 0,5724 | 7,75 | -7,177 | 4,945 | 3 | 3 | 2,053 | 12 | No | ns | 0,6981 |
| RT(Inhibitory) vs. intralaminar | 0,5724 | 4,154 | -3,582 | 4,945 | 3 | 3 | 1,024 | 12 | No | ns | 0,975 |
| ventral vs. anterior | 3,056 | 9,028 | -5,972 | 4,945 | 3 | 3 | 1,708 | 12 | No | ns | 0,8252 |
| ventral vs. midline | 3,056 | 7,75 | -4,694 | 4,945 | 3 | 3 | 1,342 | 12 | No | ns | 0,9254 |
| ventral vs. intralaminar | 3,056 | 4,154 | -1,098 | 4,945 | 3 | 3 | 0,314 | 12 | No | ns | > 0,9999 |
| anterior vs. midline | 9,028 | 7,75 | 1,279 | 4,945 | 3 | 3 | 0,3657 | 12 | No | ns | 0,9998 |
| anterior vs. intralaminar | 9,028 | 4,154 | 4,874 | 4,945 | 3 | 3 | 1,394 | 12 | No | ns | 0,914 |
| midline vs. intralaminar | 7,75 | 4,154 | 3,596 | 4,945 | 3 | 3 | 1,028 | 12 | No | ns | 0,9746 |

**Projection from Thalamic Nuclei to OFC Subdivisions**

**Table Analyzed** **MO**

**ANOVA summary**

| | |
|---|---|
| F | 2,191 |
| P value | 0,1233 |
| P value summary | ns |
| Are differences among means s | No |
| R square | 0,4773 |

**Tukey's multiple comparisons test**

| | Mean 1 | Mean 2 | Mean Diff, | SE of diff, | n1 | n2 | q | DF | Significant? | Summary | Adjusted P Value |
|---|---|---|---|---|---|---|---|---|---|---|---|
| Medial vs. RT(Inhibitory) | 6,628 | 1,269 | 5,36 | 2,377 | 3 | 3 | 3,189 | 12 | No | ns | 0,2827 |
| Medial vs. ventral | 6,628 | 1,345 | 5,283 | 2,377 | 3 | 3 | 3,143 | 12 | No | ns | 0,2956 |
| Medial vs. anterior | 6,628 | 2,993 | 3,635 | 2,377 | 3 | 3 | 2,163 | 12 | No | ns | 0,6541 |
| Medial vs. midline | 6,628 | 6,459 | 0,1689 | 2,377 | 3 | 3 | 0,1005 | 12 | No | ns | > 0,9999 |
| Medial vs. intralaminar | 6,628 | 1,9 | 4,728 | 2,377 | 3 | 3 | 2,813 | 12 | No | ns | 0,4009 |
| RT(Inhibitory) vs. ventral | 1,269 | 1,345 | -0,07691 | 2,377 | 3 | 3 | 0,04576 | 12 | No | ns | > 0,9999 |
| RT(Inhibitory) vs. anterior | 1,269 | 2,993 | -1,725 | 2,377 | 3 | 3 | 1,026 | 12 | No | ns | 0,9748 |
| RT(Inhibitory) vs. midline | 1,269 | 6,459 | -5,191 | 2,377 | 3 | 3 | 3,088 | 12 | No | ns | 0,3116 |
| RT(Inhibitory) vs. intralaminar | 1,269 | 1,9 | -0,6315 | 2,377 | 3 | 3 | 0,3757 | 12 | No | ns | 0,9998 |
| ventral vs. anterior | 1,345 | 2,993 | -1,648 | 2,377 | 3 | 3 | 0,9804 | 12 | No | ns | 0,9793 |
| ventral vs. midline | 1,345 | 6,459 | -5,114 | 2,377 | 3 | 3 | 3,043 | 12 | No | ns | 0,3254 |
| ventral vs. intralaminar | 1,345 | 1,9 | -0,5546 | 2,377 | 3 | 3 | 0,33 | 12 | No | ns | 0,9999 |
| anterior vs. midline | 2,993 | 6,459 | -3,466 | 2,377 | 3 | 3 | 2,062 | 12 | No | ns | 0,6944 |
| anterior vs. intralaminar | 2,993 | 1,9 | 1,093 | 2,377 | 3 | 3 | 0,6504 | 12 | No | ns | 0,9968 |
| midline vs. intralaminar | 6,459 | 1,9 | 4,559 | 2,377 | 3 | 3 | 2,713 | 12 | No | ns | 0,4369 |

**Table Analyzed** **VO**

**ANOVA summary**

| | |
|---|---|
| F | 3,47 |
| P value | 0,0227 |
| P value summary | * |
| Are differences among means s | Yes |
| R square | 0,4908 |

**Tukey's multiple comparisons test**

| | Mean 1 | Mean 2 | Mean Diff, | SE of diff, | n1 | n2 | q | DF | Significant? | Summary | Adjusted P Value |
|---|---|---|---|---|---|---|---|---|---|---|---|
| Medial vs. RT(Inhibitory) | 3,992 | 0,8965 | 3,095 | 3,112 | 4 | 4 | 1,407 | 18 | No | ns | 0,9138 |
| Medial vs. ventral | 3,992 | 2,787 | 1,204 | 3,112 | 4 | 4 | 0,5473 | 18 | No | ns | 0,9987 |
| Medial vs. anterior | 3,992 | 2,793 | 1,199 | 3,112 | 4 | 4 | 0,5447 | 18 | No | ns | 0,9987 |
| Medial vs. midline | 3,992 | 12,49 | -8,502 | 3,112 | 4 | 4 | 3,864 | 18 | No | ns | 0,1168 |
| Medial vs. intralaminar | 3,992 | 3,455 | 0,5367 | 3,112 | 4 | 4 | 0,2439 | 18 | No | ns | > 0,9999 |
| RT(Inhibitory) vs. ventral | 0,8965 | 2,787 | -1,891 | 3,112 | 4 | 4 | 0,8593 | 18 | No | ns | 0,9891 |
| RT(Inhibitory) vs. anterior | 0,8965 | 2,793 | -1,897 | 3,112 | 4 | 4 | 0,8619 | 18 | No | ns | 0,9889 |
| RT(Inhibitory) vs. midline | 0,8965 | 12,49 | -11,6 | 3,112 | 4 | 4 | 5,27 | 18 | Yes | * | 0,0164 |
| RT(Inhibitory) vs. intralaminar | 0,8965 | 3,455 | -2,558 | 3,112 | 4 | 4 | 1,163 | 18 | No | ns | 0,9595 |
| ventral vs. anterior | 2,787 | 2,793 | -0,00572 | 3,112 | 4 | 4 | 0,002598 | 18 | No | ns | > 0,9999 |
| ventral vs. midline | 2,787 | 12,49 | -9,707 | 3,112 | 4 | 4 | 4,411 | 18 | No | ns | 0,0562 |
| ventral vs. intralaminar | 2,787 | 3,455 | -0,6676 | 3,112 | 4 | 4 | 0,3034 | 18 | No | ns | > 0,9999 |
| anterior vs. midline | 2,793 | 12,49 | -9,701 | 3,112 | 4 | 4 | 4,408 | 18 | No | ns | 0,0564 |
| anterior vs. intralaminar | 2,793 | 3,455 | -0,6619 | 3,112 | 4 | 4 | 0,3008 | 18 | No | ns | > 0,9999 |
| midline vs. intralaminar | 12,49 | 3,455 | 9,039 | 3,112 | 4 | 4 | 4,108 | 18 | No | ns | 0,0849 |

**Table Analyzed** **LO**

**ANOVA summary**

| | |
|---|---|
| F | 4,607 |
| P value | 0,0016 |
| P value summary | ** |
| Are differences among means s | Yes |
| R square | 0,3243 |

**Tukey's multiple comparisons test**

| | Mean 1 | Mean 2 | Mean Diff, | SE of diff, | n1 | n2 | q | DF | Significant? | Summary | Adjusted P Value |
|---|---|---|---|---|---|---|---|---|---|---|---|
| Medial vs. RT(Inhibitory) | 5,487 | 1,412 | 4,075 | 2,525 | 9 | 9 | 2,282 | 48 | No | ns | 0,5938 |
| Medial vs. ventral | 5,487 | 6,523 | -1,036 | 2,525 | 9 | 9 | 0,5799 | 48 | No | ns | 0,9984 |

| | | | | | | | | | | | |
|---|---|---|---|---|---|---|---|---|---|---|---|
| Medial vs. anterior | 5,487 | 1,623 | 3,863 | 2,525 | 9 | 9 | 2,164 | 48 | No | ns | 0,6471 |
| Medial vs. midline | 5,487 | 11,84 | -6,351 | 2,525 | 9 | 9 | 3,557 | 48 | No | ns | 0,1401 |
| Medial vs. intralaminar | 5,487 | 6,197 | -0,7096 | 2,525 | 9 | 9 | 0,3974 | 48 | No | ns | 0,9997 |
| RT(Inhibitory) vs. ventral | 1,412 | 6,523 | -5,111 | 2,525 | 9 | 9 | 2,862 | 48 | No | ns | 0,3445 |
| RT(Inhibitory) vs. anterior | 1,412 | 1,623 | -0,2116 | 2,525 | 9 | 9 | 0,1185 | 48 | No | ns | > 0,9999 |
| RT(Inhibitory) vs. midline | 1,412 | 11,84 | -10,43 | 2,525 | 9 | 9 | 5,839 | 48 | Yes | ** | 0,0019 |
| RT(Inhibitory) vs. intralaminar | 1,412 | 6,197 | -4,785 | 2,525 | 9 | 9 | 2,68 | 48 | No | ns | 0,4178 |
| ventral vs. anterior | 6,523 | 1,623 | 4,899 | 2,525 | 9 | 9 | 2,744 | 48 | No | ns | 0,3913 |
| ventral vs. midline | 6,523 | 11,84 | -5,315 | 2,525 | 9 | 9 | 2,977 | 48 | No | ns | 0,3022 |
| ventral vs. intralaminar | 6,523 | 6,197 | 0,3259 | 2,525 | 9 | 9 | 0,1825 | 48 | No | ns | > 0,9999 |
| anterior vs. midline | 1,623 | 11,84 | -10,21 | 2,525 | 9 | 9 | 5,72 | 48 | Yes | ** | 0,0025 |
| anterior vs. intralaminar | 1,623 | 6,197 | -4,573 | 2,525 | 9 | 9 | 2,561 | 48 | No | ns | 0,4686 |
| midline vs. intralaminar | 11,84 | 6,197 | 5,641 | 2,525 | 9 | 9 | 3,159 | 48 | No | ns | 0,2417 |

**Table Analyzed** **AI**

**ANOVA summary**

| | |
|---|---|
| F | 2,044 |
| P value | 0,1007 |
| P value summary | ns |
| Are differences among means s | No |
| R square | 0,2541 |

**Tukey's multiple comparisons test**

| | Mean 1 | Mean 2 | Mean Diff, | SE of diff, | n1 | n2 | q | DF | Significant? | Summary | Adjusted P Value |
|---|---|---|---|---|---|---|---|---|---|---|---|
| Medial vs. RT(Inhibitory) | 5,804 | 0,3671 | 5,437 | 2,11 | 6 | 6 | 3,645 | 30 | No | ns | 0,1344 |
| Medial vs. ventral | 5,804 | 1,618 | 4,186 | 2,11 | 6 | 6 | 2,806 | 30 | No | ns | 0,3743 |
| Medial vs. anterior | 5,804 | 0,3068 | 5,498 | 2,11 | 6 | 6 | 3,685 | 30 | No | ns | 0,1269 |
| Medial vs. midline | 5,804 | 3,103 | 2,702 | 2,11 | 6 | 6 | 1,811 | 30 | No | ns | 0,793 |
| Medial vs. intralaminar | 5,804 | 3,653 | 2,152 | 2,11 | 6 | 6 | 1,443 | 30 | No | ns | 0,9075 |
| RT(Inhibitory) vs. ventral | 0,3671 | 1,618 | -1,251 | 2,11 | 6 | 6 | 0,8386 | 30 | No | ns | 0,9907 |
| RT(Inhibitory) vs. anterior | 0,3671 | 0,3068 | 0,06035 | 2,11 | 6 | 6 | 0,04045 | 30 | No | ns | > 0,9999 |
| RT(Inhibitory) vs. midline | 0,3671 | 3,103 | -2,736 | 2,11 | 6 | 6 | 1,834 | 30 | No | ns | 0,7844 |
| RT(Inhibitory) vs. intralaminar | 0,3671 | 3,653 | -3,285 | 2,11 | 6 | 6 | 2,202 | 30 | No | ns | 0,6315 |
| ventral vs. anterior | 1,618 | 0,3068 | 1,311 | 2,11 | 6 | 6 | 0,879 | 30 | No | ns | 0,9885 |
| ventral vs. midline | 1,618 | 3,103 | -1,485 | 2,11 | 6 | 6 | 0,9954 | 30 | No | ns | 0,98 |
| ventral vs. intralaminar | 1,618 | 3,653 | -2,034 | 2,11 | 6 | 6 | 1,364 | 30 | No | ns | 0,9256 |
| anterior vs. midline | 0,3068 | 3,103 | -2,796 | 2,11 | 6 | 6 | 1,874 | 30 | No | ns | 0,7689 |
| anterior vs. intralaminar | 0,3068 | 3,653 | -3,346 | 2,11 | 6 | 6 | 2,243 | 30 | No | ns | 0,6137 |
| midline vs. intralaminar | 3,103 | 3,653 | -0,5496 | 2,11 | 6 | 6 | 0,3685 | 30 | No | ns | 0,9998 |

**Table Analyzed** **DLO**

**ANOVA summary**

| | |
|---|---|
| F | 26,02 |
| P value | 0,0005 |
| P value summary | *** |
| Are differences among means s | Yes |
| R square | 0,9559 |

**Tukey's multiple comparisons test**

| | Mean 1 | Mean 2 | Mean Diff, | SE of diff, | n1 | n2 | q | DF | Significant? | Summary | Adjusted P Value |
|---|---|---|---|---|---|---|---|---|---|---|---|
| Medial vs. RT(Inhibitory) | 4,853 | 0,9477 | 3,905 | 0,7997 | 2 | 2 | 6,907 | 6 | Yes | * | 0,02 |
| Medial vs. ventral | 4,853 | 5,694 | -0,8411 | 0,7997 | 2 | 2 | 1,488 | 6 | No | ns | 0,884 |
| Medial vs. anterior | 4,853 | 0,2983 | 4,555 | 0,7997 | 2 | 2 | 8,055 | 6 | Yes | ** | 0,0095 |
| Medial vs. midline | 4,853 | 5,006 | -0,1525 | 0,7997 | 2 | 2 | 0,2697 | 6 | No | ns | > 0,9999 |
| Medial vs. intralaminar | 4,853 | 7,744 | -2,891 | 0,7997 | 2 | 2 | 5,113 | 6 | No | ns | 0,074 |
| RT(Inhibitory) vs. ventral | 0,9477 | 5,694 | -4,747 | 0,7997 | 2 | 2 | 8,394 | 6 | Yes | ** | 0,0077 |
| RT(Inhibitory) vs. anterior | 0,9477 | 0,2983 | 0,6493 | 0,7997 | 2 | 2 | 1,148 | 6 | No | ns | 0,9548 |
| RT(Inhibitory) vs. midline | 0,9477 | 5,006 | -4,058 | 0,7997 | 2 | 2 | 7,176 | 6 | Yes | * | 0,0167 |
| RT(Inhibitory) vs. intralaminar | 0,9477 | 7,744 | -6,797 | 0,7997 | 2 | 2 | 12,02 | 6 | Yes | ** | 0,0011 |
| ventral vs. anterior | 5,694 | 0,2983 | 5,396 | 0,7997 | 2 | 2 | 9,543 | 6 | Yes | ** | 0,004 |
| ventral vs. midline | 5,694 | 5,006 | 0,6887 | 0,7997 | 2 | 2 | 1,218 | 6 | No | ns | 0,9433 |
| ventral vs. intralaminar | 5,694 | 7,744 | -2,05 | 0,7997 | 2 | 2 | 3,625 | 6 | No | ns | 0,2387 |
| anterior vs. midline | 0,2983 | 5,006 | -4,707 | 0,7997 | 2 | 2 | 8,325 | 6 | Yes | ** | 0,008 |
| anterior vs. intralaminar | 0,2983 | 7,744 | -7,446 | 0,7997 | 2 | 2 | 13,17 | 6 | Yes | *** | 0,0007 |
| midline vs. intralaminar | 5,006 | 7,744 | -2,739 | 0,7997 | 2 | 2 | 4,843 | 6 | No | ns | 0,0912 |

### Intra OFC Input

**Table Analyzed** **MO**

**ANOVA summary**

| | |
|---|---|
| F | 196,1 |
| P value | < 0,0001 |
| P value summary | **** |
| Are differences among mea | Yes |
| R square | 0,9932 |

**Tukey's multiple comparisons test**

| | Mean 1 | Mean 2 | Mean Diff, | SE of diff, | n1 | n2 | q | DF | Significant? | Summary | Adjusted P Value |
|---|---|---|---|---|---|---|---|---|---|---|---|
| DLO vs. VO | 0 | 77,35 | -77,35 | 3,649 | 2 | 2 | 29,98 | 4 | Yes | *** | 0,0001 |
| DLO vs. LO | 0 | 20,72 | -20,72 | 3,649 | 2 | 2 | 8,028 | 4 | Yes | * | 0,0162 |
| DLO vs. AI | 0 | 1,936 | -1,936 | 3,649 | 2 | 2 | 0,7503 | 4 | No | ns | 0,9471 |
| VO vs. LO | 77,35 | 20,72 | 56,63 | 3,649 | 2 | 2 | 21,95 | 4 | Yes | *** | 0,0004 |
| VO vs. AI | 77,35 | 1,936 | 75,41 | 3,649 | 2 | 2 | 29,23 | 4 | Yes | *** | 0,0001 |
| LO vs. AI | 20,72 | 1,936 | 18,78 | 3,649 | 2 | 2 | 7,278 | 4 | Yes | * | 0,0228 |

**Table Analyzed** **VO**

**ANOVA summary**

| | |
|---|---|
| F | 6,255 |
| P value | 0,0084 |
| P value summary | ** |
| Are differences among mea | Yes |
| R square | 0,61 |

**Tukey's multiple comparisons test**

| | Mean 1 | Mean 2 | Mean Diff, | SE of diff, | n1 | n2 | q | DF | Significant? | Summary | Adjusted P Value |
|---|---|---|---|---|---|---|---|---|---|---|---|
| DLO vs. LO | 7,485 | 45,69 | -38,2 | 10,54 | 4 | 4 | 5,125 | 12 | Yes | * | 0,0159 |
| DLO vs. MO | 7,485 | 35,72 | -28,23 | 10,54 | 4 | 4 | 3,788 | 12 | No | ns | 0,0821 |
| DLO vs. AI | 7,485 | 11,11 | -3,623 | 10,54 | 4 | 4 | 0,4861 | 12 | No | ns | 0,9854 |
| LO vs. MO | 45,69 | 35,72 | 9,97 | 10,54 | 4 | 4 | 1,338 | 12 | No | ns | 0,7814 |
| LO vs. AI | 45,69 | 11,11 | 34,58 | 10,54 | 4 | 4 | 4,639 | 12 | Yes | * | 0,0291 |
| MO vs. AI | 35,72 | 11,11 | 24,61 | 10,54 | 4 | 4 | 3,302 | 12 | No | ns | 0,1444 |

**Table Analyzed** **LO**

**ANOVA summary**

| | |
|---|---|
| F | 4,47 |
| P value | 0,0401 |
| P value summary | * |
| Are differences among mea | Yes |
| R square | 0,6264 |

**Tukey's multiple comparisons test**

| | Mean 1 | Mean 2 | Mean Diff, | SE of diff, | n1 | n2 | q | DF | Significant? | Summary | Adjusted P Value |
|---|---|---|---|---|---|---|---|---|---|---|---|
| DLO vs. VO | 36,11 | 35,15 | 0,9653 | 8,274 | 3 | 3 | 0,165 | 8 | No | ns | 0,9994 |
| DLO vs. MO | 36,11 | 12,57 | 23,55 | 8,274 | 3 | 3 | 4,024 | 8 | No | ns | 0,0827 |
| DLO vs. AI | 36,11 | 16,17 | 19,95 | 8,274 | 3 | 3 | 3,409 | 8 | No | ns | 0,1521 |
| VO vs. MO | 35,15 | 12,57 | 22,58 | 8,274 | 3 | 3 | 3,859 | 8 | No | ns | 0,0975 |
| VO vs. AI | 35,15 | 16,17 | 18,98 | 8,274 | 3 | 3 | 3,244 | 8 | No | ns | 0,1785 |
| MO vs. AI | 12,57 | 16,17 | -3,598 | 8,274 | 3 | 3 | 0,6149 | 8 | No | ns | 0,9707 |

**Table Analyzed** **AI**

**ANOVA summary**

| | |
|---|---|
| F | 18,82 |
| P value | 0,0006 |
| P value summary | *** |
| Are differences among mea | Yes |
| R square | 0,8759 |

**Tukey's multiple comparisons test**

| | Mean 1 | Mean 2 | Mean Diff, | SE of diff, | n1 | n2 | q | DF | Significant? | Summary | Adjusted P Value |
|---|---|---|---|---|---|---|---|---|---|---|---|
| DLO vs. VO | 2,278 | 34,87 | -32,59 | 7,164 | 3 | 3 | 6,434 | 8 | Yes | ** | 0,0081 |
| DLO vs. LO | 2,278 | 12,03 | -9,751 | 7,164 | 3 | 3 | 1,925 | 8 | No | ns | 0,554 |
| DLO vs. MO | 2,278 | 50,82 | -48,54 | 7,164 | 3 | 3 | 9,583 | 8 | Yes | *** | 0,0006 |
| VO vs. LO | 34,87 | 12,03 | 22,84 | 7,164 | 3 | 3 | 4,509 | 8 | No | ns | 0,051 |
| VO vs. MO | 34,87 | 50,82 | -15,95 | 7,164 | 3 | 3 | 3,149 | 8 | No | ns | 0,1957 |
| LO vs. MO | 12,03 | 50,82 | -38,79 | 7,164 | 3 | 3 | 7,658 | 8 | Yes | ** | 0,0028 |

**Table Analyzed** **DLO**

**ANOVA summary**

| | |
|---|---|
| F | 50,21 |
| P value | < 0,0001 |
| P value summary | **** |
| Are differences among mea | Yes |
| R square | 0,9496 |

**Tukey's multiple comparisons test**

| | Mean 1 | Mean 2 | Mean Diff, | SE of diff, | n1 | n2 | q | DF | Significant? | Summary | Adjusted P Value |
|---|---|---|---|---|---|---|---|---|---|---|---|
| VO vs. LO | 9,692 | 61,05 | -51,36 | 5,012 | 3 | 3 | 14,49 | 8 | Yes | **** | < 0,0001 |
| VO vs. MO | 9,692 | 6,191 | 3,501 | 5,012 | 3 | 3 | 0,988 | 8 | No | ns | 0,8948 |
| VO vs. AI | 9,692 | 23,06 | -13,37 | 5,012 | 3 | 3 | 3,774 | 8 | No | ns | 0,1062 |
| LO vs. MO | 61,05 | 6,191 | 54,86 | 5,012 | 3 | 3 | 15,48 | 8 | Yes | **** | < 0,0001 |
| LO vs. AI | 61,05 | 23,06 | 37,99 | 5,012 | 3 | 3 | 10,72 | 8 | Yes | *** | 0,0003 |
| MO vs. AI | 6,191 | 23,06 | -16,87 | 5,012 | 3 | 3 | 4,761 | 8 | Yes | * | 0,0397 |

### Intra OFC Output

**Table Analyzed** **MO**

**ANOVA summary**

| | |
|---|---|
| F | 72,32 |
| P value | < 0,0001 |
| P value summary | **** |
| Are differences among mear | Yes |
| R square | 0,9644 |

**Tukey's multiple comparisons test**

| | Mean 1 | Mean 2 | Mean Diff, | SE of diff, | n1 | n2 | q | DF | Significant | Summary | Adjusted P Value |
|---|---|---|---|---|---|---|---|---|---|---|---|
| DLO vs. VO | 5,674 | 40,52 | -34,84 | 3,459 | 3 | 3 | 14,25 | 8 | Yes | **** | < 0,0001 |
| DLO vs. LO | 5,674 | 8,517 | -2,842 | 3,459 | 3 | 3 | 1,162 | 8 | No | ns | 0,8428 |
| DLO vs. AI | 5,674 | 45,29 | -39,62 | 3,459 | 3 | 3 | 16,2 | 8 | Yes | **** | < 0,0001 |
| VO vs. LO | 40,52 | 8,517 | 32 | 3,459 | 3 | 3 | 13,08 | 8 | Yes | **** | < 0,0001 |
| VO vs. AI | 40,52 | 45,29 | -4,775 | 3,459 | 3 | 3 | 1,952 | 8 | No | ns | 0,5435 |
| LO vs. AI | 8,517 | 45,29 | -36,78 | 3,459 | 3 | 3 | 15,04 | 8 | Yes | **** | < 0,0001 |

**Table Analyzed** **VO**

**ANOVA summary**

| | |
|---|---|
| F | 13,45 |
| P value | 0,0004 |
| P value summary | *** |
| Are differences among mear | Yes |
| R square | 0,7707 |

**Tukey's multiple comparisons test**

| | Mean 1 | Mean 2 | Mean Diff, | SE of diff, | n1 | n2 | q | DF | Significant | Summary | Adjusted P Value |
|---|---|---|---|---|---|---|---|---|---|---|---|
| DLO vs. LO | 4,925 | 49,42 | -44,5 | 7,073 | 4 | 4 | 8,897 | 12 | Yes | *** | 0,0002 |
| DLO vs. AI | 4,925 | 23,03 | -18,1 | 7,073 | 4 | 4 | 3,619 | 12 | No | ns | 0,1002 |
| DLO vs. MO | 4,925 | 22,63 | -17,7 | 7,073 | 4 | 4 | 3,539 | 12 | No | ns | 0,11 |
| LO vs. AI | 49,42 | 23,03 | 26,4 | 7,073 | 4 | 4 | 5,278 | 12 | Yes | * | 0,0132 |
| LO vs. MO | 49,42 | 22,63 | 26,8 | 7,073 | 4 | 4 | 5,358 | 12 | Yes | * | 0,0119 |
| AI vs. MO | 23,03 | 22,63 | 0,4 | 7,073 | 4 | 4 | 0,07998 | 12 | No | ns | > 0,9999 |

**Table Analyzed** **LO**

**ANOVA summary**

| | |
|---|---|
| F | 80,45 |
| P value | < 0,0001 |
| P value summary | **** |
| Are differences among mear | Yes |
| R square | 0,8829 |

**Tukey's multiple comparisons test**

| | Mean 1 | Mean 2 | Mean Diff, | SE of diff, | n1 | n2 | q | DF | Significant | Summary | Adjusted P Value |
|---|---|---|---|---|---|---|---|---|---|---|---|
| DLO vs. VO | 11,18 | 59,44 | -48,26 | 3,775 | 9 | 9 | 18,08 | 32 | Yes | **** | < 0,0001 |
| DLO vs. AI | 11,18 | 22,77 | -11,59 | 3,775 | 9 | 9 | 4,342 | 32 | Yes | * | 0,0214 |
| DLO vs. MO | 11,18 | 6,618 | 4,56 | 3,775 | 9 | 9 | 1,708 | 32 | No | ns | 0,6265 |
| VO vs. AI | 59,44 | 22,77 | 36,67 | 3,775 | 9 | 9 | 13,74 | 32 | Yes | **** | < 0,0001 |
| VO vs. MO | 59,44 | 6,618 | 52,82 | 3,775 | 9 | 9 | 19,79 | 32 | Yes | **** | < 0,0001 |
| AI vs. MO | 22,77 | 6,618 | 16,15 | 3,775 | 9 | 9 | 6,05 | 32 | Yes | *** | 0,0009 |

**Table Analyzed** **AI**

**ANOVA summary**

| | |
|---|---|
| F | 14,03 |
| P value | < 0,0001 |
| P value summary | **** |
| Are differences among mear | Yes |
| R square | 0,6779 |

**Tukey's multiple comparisons test**

| | Mean 1 | Mean 2 | Mean Diff, | SE of diff, | n1 | n2 | q | DF | Significant | Summary | Adjusted P Value |
|---|---|---|---|---|---|---|---|---|---|---|---|
| DLO vs. VO | 48,33 | 11,29 | 37,04 | 6,121 | 6 | 6 | 8,559 | 20 | Yes | **** | < 0,0001 |
| DLO vs. LO | 48,33 | 22,41 | 25,92 | 6,121 | 6 | 6 | 5,989 | 20 | Yes | ** | 0,0021 |
| DLO vs. MO | 48,33 | 17,97 | 30,36 | 6,121 | 6 | 6 | 7,016 | 20 | Yes | *** | 0,0004 |
| VO vs. LO | 11,29 | 22,41 | -11,12 | 6,121 | 6 | 6 | 2,57 | 20 | No | ns | 0,2947 |
| VO vs. MO | 11,29 | 17,97 | -6,679 | 6,121 | 6 | 6 | 1,543 | 20 | No | ns | 0,6988 |
| LO vs. MO | 22,41 | 17,97 | 4,445 | 6,121 | 6 | 6 | 1,027 | 20 | No | ns | 0,8855 |

**Table Analyzed** **DLO**

| ANOVA summary | |
|---|---|
| F | 21,24 |
| P value | 0,0064 |
| P value summary | ** |
| Are differences among mear | Yes |
| R square | 0,9409 |

**Tukey's multiple comparisons test**

| | Mean 1 | Mean 2 | Mean Diff, | SE of diff, | n1 | n2 | q | DF | Significant | Summary | Adjusted P Value |
|---|---|---|---|---|---|---|---|---|---|---|---|
| VO vs. LO | 15,86 | 58,91 | -43,05 | 7,439 | 2 | 2 | 8,184 | 4 | Yes | * | 0,0151 |
| VO vs. AI | 15,86 | 23,19 | -7,334 | 7,439 | 2 | 2 | 1,394 | 4 | No | ns | 0,7656 |
| VO vs. MO | 15,86 | 2,044 | 13,81 | 7,439 | 2 | 2 | 2,626 | 4 | No | ns | 0,3692 |
| LO vs. AI | 58,91 | 23,19 | 35,72 | 7,439 | 2 | 2 | 6,79 | 4 | Yes | * | 0,029 |
| LO vs. MO | 58,91 | 2,044 | 56,86 | 7,439 | 2 | 2 | 10,81 | 4 | Yes | ** | 0,0054 |
| AI vs. MO | 23,19 | 2,044 | 21,15 | 7,439 | 2 | 2 | 4,02 | 4 | No | ns | 0,1439 |
