## Supplementary 1 for "Anatomical input-output streams within mouse orbitofrontal cortex subdivisions"

| Table Analyzed | MO inputs |
|---|---|
| **ANOVA summary** | |
| F | 77,86 |
| P value | < 0,0001 |
| P value summary | **** |
| Are differences among means statistically s | Yes |
| R square | 0,9862 |

**Tukey's multiple comparisons test**

| | Mean 1 | Mean 2 | Mean Diff, | SE of diff, | n1 | n2 | q | DF | Significant | Summary | Adjusted P Value |
|---|---|---|---|---|---|---|---|---|---|---|---|
| Sensory Cortex vs. Motor cortex | 2,949 | 0,5037 | 2,445 | 1,768 | 2 | 2 | 1,956 | 12 | No | ns | 0,9465 |
| Sensory Cortex vs. Association cortex | 2,949 | 13,71 | -10,76 | 1,768 | 2 | 2 | 8,604 | 12 | Yes | ** | 0,0019 |
| Sensory Cortex vs. Prefrontal cortex | 2,949 | 20,29 | -17,34 | 1,768 | 2 | 2 | 13,87 | 12 | Yes | **** | < 0,0001 |
| Sensory Cortex vs. Olfactory regions | 2,949 | 1,543 | 1,406 | 1,768 | 2 | 2 | 1,124 | 12 | No | ns | 0,9992 |
| Sensory Cortex vs. Hippocampus | 2,949 | 0,7232 | 2,226 | 1,768 | 2 | 2 | 1,78 | 12 | No | ns | 0,9705 |
| Sensory Cortex vs. Thalamus | 2,949 | 37,38 | -34,43 | 1,768 | 2 | 2 | 27,54 | 12 | Yes | **** | < 0,0001 |
| Sensory Cortex vs. Amygdala | 2,949 | 9,629 | -6,68 | 1,768 | 2 | 2 | 5,344 | 12 | No | ns | 0,0676 |
| Sensory Cortex vs. Hypothalamus | 2,949 | 1,578 | 1,371 | 1,768 | 2 | 2 | 1,096 | 12 | No | ns | 0,9993 |
| Sensory Cortex vs. Brainstem | 2,949 | 1,183 | 1,766 | 1,768 | 2 | 2 | 1,413 | 12 | No | ns | 0,9945 |
| Sensory Cortex vs. Striatum | 2,949 | 2,873 | 0,0756 | 1,768 | 2 | 2 | 0,0605 | 12 | No | ns | > 0,9999 |
| Sensory Cortex vs. Claustrum | 2,949 | 7,645 | -4,696 | 1,768 | 2 | 2 | 3,757 | 12 | No | ns | 0,3447 |
| Motor cortex vs. Association cortex | 0,5037 | 13,71 | -13,2 | 1,768 | 2 | 2 | 10,56 | 12 | Yes | *** | 0,0003 |
| Motor cortex vs. Prefrontal cortex | 0,5037 | 20,29 | -19,78 | 1,768 | 2 | 2 | 15,83 | 12 | Yes | **** | < 0,0001 |
| Motor cortex vs. Olfactory regions | 0,5037 | 1,543 | -1,04 | 1,768 | 2 | 2 | 0,8315 | 12 | No | ns | > 0,9999 |
| Motor cortex vs. Hippocampus | 0,5037 | 0,7232 | -0,2194 | 1,768 | 2 | 2 | 0,1755 | 12 | No | ns | > 0,9999 |
| Motor cortex vs. Thalamus | 0,5037 | 37,38 | -36,88 | 1,768 | 2 | 2 | 29,5 | 12 | Yes | **** | < 0,0001 |
| Motor cortex vs. Amygdala | 0,5037 | 9,629 | -9,125 | 1,768 | 2 | 2 | 7,299 | 12 | Yes | ** | 0,0077 |
| Motor cortex vs. Hypothalamus | 0,5037 | 1,578 | -1,074 | 1,768 | 2 | 2 | 0,8595 | 12 | No | ns | > 0,9999 |
| Motor cortex vs. Brainstem | 0,5037 | 1,183 | -0,679 | 1,768 | 2 | 2 | 0,5431 | 12 | No | ns | > 0,9999 |
| Motor cortex vs. Striatum | 0,5037 | 2,873 | -2,369 | 1,768 | 2 | 2 | 1,895 | 12 | No | ns | 0,9559 |
| Motor cortex vs. Claustrum | 0,5037 | 7,645 | -7,142 | 1,768 | 2 | 2 | 5,713 | 12 | Yes | * | 0,0448 |
| Association cortex vs. Prefrontal cortex | 13,71 | 20,29 | -6,583 | 1,768 | 2 | 2 | 5,266 | 12 | No | ns | 0,0736 |
| Association cortex vs. Olfactory regions | 13,71 | 1,543 | 12,16 | 1,768 | 2 | 2 | 9,729 | 12 | Yes | *** | 0,0006 |
| Association cortex vs. Hippocampus | 13,71 | 0,7232 | 12,98 | 1,768 | 2 | 2 | 10,38 | 12 | Yes | *** | 0,0003 |
| Association cortex vs. Thalamus | 13,71 | 37,38 | -23,67 | 1,768 | 2 | 2 | 18,94 | 12 | Yes | **** | < 0,0001 |
| Association cortex vs. Amygdala | 13,71 | 9,629 | 4,076 | 1,768 | 2 | 2 | 3,261 | 12 | No | ns | 0,5178 |
| Association cortex vs. Hypothalamus | 13,71 | 1,578 | 12,13 | 1,768 | 2 | 2 | 9,701 | 12 | Yes | *** | 0,0006 |
| Association cortex vs. Brainstem | 13,71 | 1,183 | 12,52 | 1,768 | 2 | 2 | 10,02 | 12 | Yes | *** | 0,0005 |
| Association cortex vs. Striatum | 13,71 | 2,873 | 10,83 | 1,768 | 2 | 2 | 8,665 | 12 | Yes | ** | 0,0018 |
| Association cortex vs. Claustrum | 13,71 | 7,645 | 6,06 | 1,768 | 2 | 2 | 4,847 | 12 | No | ns | 0,1161 |
| Prefrontal cortex vs. Olfactory regions | 20,29 | 1,543 | 18,75 | 1,768 | 2 | 2 | 14,99 | 12 | Yes | **** | < 0,0001 |
| Prefrontal cortex vs. Hippocampus | 20,29 | 0,7232 | 19,57 | 1,768 | 2 | 2 | 15,65 | 12 | Yes | **** | < 0,0001 |
| Prefrontal cortex vs. Thalamus | 20,29 | 37,38 | -17,09 | 1,768 | 2 | 2 | 13,67 | 12 | Yes | **** | < 0,0001 |
| Prefrontal cortex vs. Amygdala | 20,29 | 9,629 | 10,66 | 1,768 | 2 | 2 | 8,527 | 12 | Yes | ** | 0,0021 |
| Prefrontal cortex vs. Hypothalamus | 20,29 | 1,578 | 18,71 | 1,768 | 2 | 2 | 14,97 | 12 | Yes | **** | < 0,0001 |
| Prefrontal cortex vs. Brainstem | 20,29 | 1,183 | 19,11 | 1,768 | 2 | 2 | 15,28 | 12 | Yes | **** | < 0,0001 |
| Prefrontal cortex vs. Striatum | 20,29 | 2,873 | 17,42 | 1,768 | 2 | 2 | 13,93 | 12 | Yes | **** | < 0,0001 |
| Prefrontal cortex vs. Claustrum | 20,29 | 7,645 | 12,64 | 1,768 | 2 | 2 | 10,11 | 12 | Yes | *** | 0,0004 |
| Olfactory regions vs. Hippocampus | 1,543 | 0,7232 | 0,8201 | 1,768 | 2 | 2 | 0,656 | 12 | No | ns | > 0,9999 |
| Olfactory regions vs. Thalamus | 1,543 | 37,38 | -35,84 | 1,768 | 2 | 2 | 28,67 | 12 | Yes | **** | < 0,0001 |
| Olfactory regions vs. Amygdala | 1,543 | 9,629 | -8,086 | 1,768 | 2 | 2 | 6,468 | 12 | Yes | * | 0,0193 |
| Olfactory regions vs. Hypothalamus | 1,543 | 1,578 | -0,03496 | 1,768 | 2 | 2 | 0,028 | 12 | No | ns | > 0,9999 |
| Olfactory regions vs. Brainstem | 1,543 | 1,183 | 0,3605 | 1,768 | 2 | 2 | 0,2884 | 12 | No | ns | > 0,9999 |
| Olfactory regions vs. Striatum | 1,543 | 2,873 | -1,33 | 1,768 | 2 | 2 | 1,064 | 12 | No | ns | 0,9995 |
| Olfactory regions vs. Claustrum | 1,543 | 7,645 | -6,102 | 1,768 | 2 | 2 | 4,881 | 12 | No | ns | 0,112 |
| Hippocampus vs. Thalamus | 0,7232 | 37,38 | -36,66 | 1,768 | 2 | 2 | 29,32 | 12 | Yes | **** | < 0,0001 |
| Hippocampus vs. Amygdala | 0,7232 | 9,629 | -8,906 | 1,768 | 2 | 2 | 7,124 | 12 | Yes | ** | 0,0093 |
| Hippocampus vs. Hypothalamus | 0,7232 | 1,578 | -0,855 | 1,768 | 2 | 2 | 0,684 | 12 | No | ns | > 0,9999 |
| Hippocampus vs. Brainstem | 0,7232 | 1,183 | -0,4595 | 1,768 | 2 | 2 | 0,3676 | 12 | No | ns | > 0,9999 |
| Hippocampus vs. Striatum | 0,7232 | 2,873 | -2,15 | 1,768 | 2 | 2 | 1,72 | 12 | No | ns | 0,9766 |
| Hippocampus vs. Claustrum | 0,7232 | 7,645 | -6,922 | 1,768 | 2 | 2 | 5,537 | 12 | No | ns | 0,0545 |
| Thalamus vs. Amygdala | 37,38 | 9,629 | 27,75 | 1,768 | 2 | 2 | 22,2 | 12 | Yes | **** | < 0,0001 |
| Thalamus vs. Hypothalamus | 37,38 | 1,578 | 35,8 | 1,768 | 2 | 2 | 28,64 | 12 | Yes | **** | < 0,0001 |
| Thalamus vs. Brainstem | 37,38 | 1,183 | 36,2 | 1,768 | 2 | 2 | 28,95 | 12 | Yes | **** | < 0,0001 |
| Thalamus vs. Striatum | 37,38 | 2,873 | 34,51 | 1,768 | 2 | 2 | 27,6 | 12 | Yes | **** | < 0,0001 |
| Thalamus vs. Claustrum | 37,38 | 7,645 | 29,73 | 1,768 | 2 | 2 | 23,79 | 12 | Yes | **** | < 0,0001 |
| Amygdala vs. Hypothalamus | 9,629 | 1,578 | 8,051 | 1,768 | 2 | 2 | 6,44 | 12 | Yes | * | 0,0199 |
| Amygdala vs. Brainstem | 9,629 | 1,183 | 8,446 | 1,768 | 2 | 2 | 6,756 | 12 | Yes | * | 0,014 |
| Amygdala vs. Striatum | 9,629 | 2,873 | 6,756 | 1,768 | 2 | 2 | 5,404 | 12 | No | ns | 0,0632 |
| Amygdala vs. Claustrum | 9,629 | 7,645 | 1,984 | 1,768 | 2 | 2 | 1,587 | 12 | No | ns | 0,9867 |
| Hypothalamus vs. Brainstem | 1,578 | 1,183 | 0,3955 | 1,768 | 2 | 2 | 0,3164 | 12 | No | ns | > 0,9999 |
| Hypothalamus vs. Striatum | 1,578 | 2,873 | -1,295 | 1,768 | 2 | 2 | 1,036 | 12 | No | ns | 0,9996 |
| Hypothalamus vs. Claustrum | 1,578 | 7,645 | -6,067 | 1,768 | 2 | 2 | 4,853 | 12 | No | ns | 0,1154 |
| Brainstem vs. Striatum | 1,183 | 2,873 | -1,69 | 1,768 | 2 | 2 | 1,352 | 12 | No | ns | 0,9961 |
| Brainstem vs. Claustrum | 1,183 | 7,645 | -6,463 | 1,768 | 2 | 2 | 5,17 | 12 | No | ns | 0,0818 |
| Striatum vs. Claustrum | 2,873 | 7,645 | -4,772 | 1,768 | 2 | 2 | 3,817 | 12 | No | ns | 0,3264 |

| Table Analyzed | VO inputs |
|---|---|

**ANOVA summary**

| | |
|---|---|
| F | 51,49 |
| P value | < 0,0001 |
| P value summary | **** |
| Are differences among means statistically signif | Yes |
| R square | 0,9402 |

**Tukey's multiple comparisons test**

| | Mean 1 | Mean 2 | Mean Diff, | SE of diff, | n1 | n2 | q | Significant | Summary | Adjusted P Value |
|---|---|---|---|---|---|---|---|---|---|---|
| Sensory Cortex vs. Motor cortex | 3,503 | 2,718 | 0,7854 | 2,804 | 4 | 4 | 0,3962 | No | ns | > 0,9999 |
| Sensory Cortex vs. Association cortex | 3,503 | 7,089 | -3,585 | 2,804 | 4 | 4 | 1,808 | No | ns | 0,9768 |
| Sensory Cortex vs. Prefrontal cortex | 3,503 | 9,472 | -5,968 | 2,804 | 4 | 4 | 3,011 | No | ns | 0,607 |
| Sensory Cortex vs. Olfactory regions | 3,503 | 0,34 | 3,164 | 2,804 | 4 | 4 | 1,596 | No | ns | 0,9911 |
| Sensory Cortex vs. Hippocampus | 3,503 | 0,5784 | 2,925 | 2,804 | 4 | 4 | 1,475 | No | ns | 0,9953 |
| Sensory Cortex vs. Thalamus | 3,503 | 52,6 | -49,1 | 2,804 | 4 | 4 | 24,76 | Yes | **** | < 0,0001 |
| Sensory Cortex vs. Amygdala | 3,503 | 6,647 | -3,144 | 2,804 | 4 | 4 | 1,586 | No | ns | 0,9915 |
| Sensory Cortex vs. Hypothalamus | 3,503 | 0,9411 | 2,562 | 2,804 | 4 | 4 | 1,293 | No | ns | 0,9985 |
| Sensory Cortex vs. Brainstem | 3,503 | 5,525 | -2,021 | 2,804 | 4 | 4 | 1,02 | No | ns | 0,9998 |
| Sensory Cortex vs. Striatum | 3,503 | 4,569 | -1,066 | 2,804 | 4 | 4 | 0,5375 | No | ns | > 0,9999 |
| Sensory Cortex vs. Claustrum | 3,503 | 6,018 | -2,514 | 2,804 | 4 | 4 | 1,268 | No | ns | 0,9987 |
| Motor cortex vs. Association cortex | 2,718 | 7,089 | -4,371 | 2,804 | 4 | 4 | 2,205 | No | ns | 0,9125 |
| Motor cortex vs. Prefrontal cortex | 2,718 | 9,472 | -6,754 | 2,804 | 4 | 4 | 3,407 | No | ns | 0,4259 |
| Motor cortex vs. Olfactory regions | 2,718 | 0,34 | 2,378 | 2,804 | 4 | 4 | 1,2 | No | ns | 0,9992 |
| Motor cortex vs. Hippocampus | 2,718 | 0,5784 | 2,14 | 2,804 | 4 | 4 | 1,079 | No | ns | 0,9997 |
| Motor cortex vs. Thalamus | 2,718 | 52,6 | -49,88 | 2,804 | 4 | 4 | 25,16 | Yes | **** | < 0,0001 |
| Motor cortex vs. Amygdala | 2,718 | 6,647 | -3,929 | 2,804 | 4 | 4 | 1,982 | No | ns | 0,9557 |
| Motor cortex vs. Hypothalamus | 2,718 | 0,9411 | 1,777 | 2,804 | 4 | 4 | 0,8963 | No | ns | > 0,9999 |
| Motor cortex vs. Brainstem | 2,718 | 5,525 | -2,807 | 2,804 | 4 | 4 | 1,416 | No | ns | 0,9967 |
| Motor cortex vs. Striatum | 2,718 | 4,569 | -1,851 | 2,804 | 4 | 4 | 0,9337 | No | ns | > 0,9999 |
| Motor cortex vs. Claustrum | 2,718 | 6,018 | -3,3 | 2,804 | 4 | 4 | 1,664 | No | ns | 0,9876 |
| Association cortex vs. Prefrontal cortex | 7,089 | 9,472 | -2,383 | 2,804 | 4 | 4 | 1,202 | No | ns | 0,9992 |
| Association cortex vs. Olfactory regions | 7,089 | 0,34 | 6,749 | 2,804 | 4 | 4 | 3,404 | No | ns | 0,427 |
| Association cortex vs. Hippocampus | 7,089 | 0,5784 | 6,51 | 2,804 | 4 | 4 | 3,284 | No | ns | 0,4804 |
| Association cortex vs. Thalamus | 7,089 | 52,6 | -45,51 | 2,804 | 4 | 4 | 22,96 | Yes | **** | < 0,0001 |
| Association cortex vs. Amygdala | 7,089 | 6,647 | 0,4412 | 2,804 | 4 | 4 | 0,2226 | No | ns | > 0,9999 |
| Association cortex vs. Hypothalamus | 7,089 | 0,9411 | 6,147 | 2,804 | 4 | 4 | 3,101 | No | ns | 0,5648 |
| Association cortex vs. Brainstem | 7,089 | 5,525 | 1,564 | 2,804 | 4 | 4 | 0,7887 | No | ns | > 0,9999 |
| Association cortex vs. Striatum | 7,089 | 4,569 | 2,52 | 2,804 | 4 | 4 | 1,271 | No | ns | 0,9987 |
| Association cortex vs. Claustrum | 7,089 | 6,018 | 1,071 | 2,804 | 4 | 4 | 0,5401 | No | ns | > 0,9999 |
| Prefrontal cortex vs. Olfactory regions | 9,472 | 0,34 | 9,132 | 2,804 | 4 | 4 | 4,606 | No | ns | 0,0862 |
| Prefrontal cortex vs. Hippocampus | 9,472 | 0,5784 | 8,894 | 2,804 | 4 | 4 | 4,486 | No | ns | 0,1042 |
| Prefrontal cortex vs. Thalamus | 9,472 | 52,6 | -43,13 | 2,804 | 4 | 4 | 21,75 | Yes | **** | < 0,0001 |
| Prefrontal cortex vs. Amygdala | 9,472 | 6,647 | 2,825 | 2,804 | 4 | 4 | 1,425 | No | ns | 0,9965 |
| Prefrontal cortex vs. Hypothalamus | 9,472 | 0,9411 | 8,531 | 2,804 | 4 | 4 | 4,303 | No | ns | 0,1375 |
| Prefrontal cortex vs. Brainstem | 9,472 | 5,525 | 3,947 | 2,804 | 4 | 4 | 1,991 | No | ns | 0,9544 |
| Prefrontal cortex vs. Striatum | 9,472 | 4,569 | 4,903 | 2,804 | 4 | 4 | 2,473 | No | ns | 0,8336 |
| Prefrontal cortex vs. Claustrum | 9,472 | 6,018 | 3,454 | 2,804 | 4 | 4 | 1,742 | No | ns | 0,9824 |
| Olfactory regions vs. Hippocampus | 0,34 | 0,5784 | -0,2384 | 2,804 | 4 | 4 | 0,1203 | No | ns | > 0,9999 |
| Olfactory regions vs. Thalamus | 0,34 | 52,6 | -52,26 | 2,804 | 4 | 4 | 26,36 | Yes | **** | < 0,0001 |
| Olfactory regions vs. Amygdala | 0,34 | 6,647 | -6,307 | 2,804 | 4 | 4 | 3,182 | No | ns | 0,5273 |
| Olfactory regions vs. Hypothalamus | 0,34 | 0,9411 | -0,6011 | 2,804 | 4 | 4 | 0,3032 | No | ns | > 0,9999 |
| Olfactory regions vs. Brainstem | 0,34 | 5,525 | -5,185 | 2,804 | 4 | 4 | 2,615 | No | ns | 0,7806 |
| Olfactory regions vs. Striatum | 0,34 | 4,569 | -4,229 | 2,804 | 4 | 4 | 2,133 | No | ns | 0,9285 |
| Olfactory regions vs. Claustrum | 0,34 | 6,018 | -5,678 | 2,804 | 4 | 4 | 2,864 | No | ns | 0,6744 |
| Hippocampus vs. Thalamus | 0,5784 | 52,6 | -52,02 | 2,804 | 4 | 4 | 26,24 | Yes | **** | < 0,0001 |
| Hippocampus vs. Amygdala | 0,5784 | 6,647 | -6,069 | 2,804 | 4 | 4 | 3,061 | No | ns | 0,5833 |
| Hippocampus vs. Hypothalamus | 0,5784 | 0,9411 | -0,3627 | 2,804 | 4 | 4 | 0,183 | No | ns | > 0,9999 |
| Hippocampus vs. Brainstem | 0,5784 | 5,525 | -4,947 | 2,804 | 4 | 4 | 2,495 | No | ns | 0,8258 |
| Hippocampus vs. Striatum | 0,5784 | 4,569 | -3,991 | 2,804 | 4 | 4 | 2,013 | No | ns | 0,9509 |
| Hippocampus vs. Claustrum | 0,5784 | 6,018 | -5,439 | 2,804 | 4 | 4 | 2,744 | No | ns | 0,7276 |
| Thalamus vs. Amygdala | 52,6 | 6,647 | 45,95 | 2,804 | 4 | 4 | 23,18 | Yes | **** | < 0,0001 |
| Thalamus vs. Hypothalamus | 52,6 | 0,9411 | 51,66 | 2,804 | 4 | 4 | 26,06 | Yes | **** | < 0,0001 |
| Thalamus vs. Brainstem | 52,6 | 5,525 | 47,07 | 2,804 | 4 | 4 | 23,75 | Yes | **** | < 0,0001 |
| Thalamus vs. Striatum | 52,6 | 4,569 | 48,03 | 2,804 | 4 | 4 | 24,23 | Yes | **** | < 0,0001 |
| Thalamus vs. Claustrum | 52,6 | 6,018 | 46,58 | 2,804 | 4 | 4 | 23,5 | Yes | **** | < 0,0001 |
| Amygdala vs. Hypothalamus | 6,647 | 0,9411 | 5,706 | 2,804 | 4 | 4 | 2,878 | No | ns | 0,6679 |
| Amygdala vs. Brainstem | 6,647 | 5,525 | 1,122 | 2,804 | 4 | 4 | 0,5662 | No | ns | > 0,9999 |
| Amygdala vs. Striatum | 6,647 | 4,569 | 2,078 | 2,804 | 4 | 4 | 1,048 | No | ns | 0,9998 |
| Amygdala vs. Claustrum | 6,647 | 6,018 | 0,6296 | 2,804 | 4 | 4 | 0,3176 | No | ns | > 0,9999 |
| Hypothalamus vs. Brainstem | 0,9411 | 5,525 | -4,584 | 2,804 | 4 | 4 | 2,312 | No | ns | 0,8844 |
| Hypothalamus vs. Striatum | 0,9411 | 4,569 | -3,628 | 2,804 | 4 | 4 | 1,83 | No | ns | 0,9747 |
| Hypothalamus vs. Claustrum | 0,9411 | 6,018 | -5,077 | 2,804 | 4 | 4 | 2,561 | No | ns | 0,8018 |
| Brainstem vs. Striatum | 5,525 | 4,569 | 0,9559 | 2,804 | 4 | 4 | 0,4821 | No | ns | > 0,9999 |
| Brainstem vs. Claustrum | 5,525 | 6,018 | -0,4929 | 2,804 | 4 | 4 | 0,2486 | No | ns | > 0,9999 |
| Striatum vs. Claustrum | 4,569 | 6,018 | -1,449 | 2,804 | 4 | 4 | 0,7308 | No | ns | > 0,9999 |

**Table Analyzed** | **LO inputs**

**ANOVA summary**

| | |
|---|---|
| F | 18,89 |
| P value | < 0,0001 |
| P value summary | **** |
| Are differences among means statistically s | Yes |
| R square | 0,8965 |

**Tukey's multiple comparisons test**

| | Mean 1 | Mean 2 | Mean Diff, | SE of diff, | n1 | n2 | q | DF | Significant | Summary | Adjusted P Value |
|---|---|---|---|---|---|---|---|---|---|---|---|
| Sensory Cortex vs. Motor cortex | 4,271 | 7,314 | -3,043 | 4,096 | 3 | 3 | 1,05 | 24 | No | ns | 0,9997 |
| Sensory Cortex vs. Association cortex | 4,271 | 7,668 | -3,397 | 4,096 | 3 | 3 | 1,173 | 24 | No | ns | 0,9993 |
| Sensory Cortex vs. Prefrontal cortex | 4,271 | 6,124 | -1,853 | 4,096 | 3 | 3 | 0,6398 | 24 | No | ns | > 0,9999 |
| Sensory Cortex vs. Olfactory regions | 4,271 | 3,755 | 0,5158 | 4,096 | 3 | 3 | 0,1781 | 24 | No | ns | > 0,9999 |
| Sensory Cortex vs. Hippocampus | 4,271 | 0,07276 | 4,198 | 4,096 | 3 | 3 | 1,449 | 24 | No | ns | 0,9953 |
| Sensory Cortex vs. Thalamus | 4,271 | 47,44 | -43,17 | 4,096 | 3 | 3 | 14,9 | 24 | Yes | **** | < 0,0001 |
| Sensory Cortex vs. Amygdala | 4,271 | 4,454 | -0,1828 | 4,096 | 3 | 3 | 0,0631 | 24 | No | ns | > 0,9999 |
| Sensory Cortex vs. Hypothalamus | 4,271 | 0,4891 | 3,782 | 4,096 | 3 | 3 | 1,306 | 24 | No | ns | 0,9981 |
| Sensory Cortex vs. Brainstem | 4,271 | 4,054 | 0,2174 | 4,096 | 3 | 3 | 0,0751 | 24 | No | ns | > 0,9999 |
| Sensory Cortex vs. Striatum | 4,271 | 5,664 | -1,393 | 4,096 | 3 | 3 | 0,481 | 24 | No | ns | > 0,9999 |
| Sensory Cortex vs. Claustrum | 4,271 | 8,697 | -4,426 | 4,096 | 3 | 3 | 1,528 | 24 | No | ns | 0,9928 |
| Motor cortex vs. Association cortex | 7,314 | 7,668 | -0,354 | 4,096 | 3 | 3 | 0,1222 | 24 | No | ns | > 0,9999 |
| Motor cortex vs. Prefrontal cortex | 7,314 | 6,124 | 1,189 | 4,096 | 3 | 3 | 0,4106 | 24 | No | ns | > 0,9999 |
| Motor cortex vs. Olfactory regions | 7,314 | 3,755 | 3,558 | 4,096 | 3 | 3 | 1,228 | 24 | No | ns | 0,9989 |
| Motor cortex vs. Hippocampus | 7,314 | 0,07276 | 7,241 | 4,096 | 3 | 3 | 2,5 | 24 | No | ns | 0,8194 |
| Motor cortex vs. Thalamus | 7,314 | 47,44 | -40,12 | 4,096 | 3 | 3 | 13,85 | 24 | Yes | **** | < 0,0001 |
| Motor cortex vs. Amygdala | 7,314 | 4,454 | 2,86 | 4,096 | 3 | 3 | 0,9873 | 24 | No | ns | 0,9999 |
| Motor cortex vs. Hypothalamus | 7,314 | 0,4891 | 6,825 | 4,096 | 3 | 3 | 2,356 | 24 | No | ns | 0,8664 |
| Motor cortex vs. Brainstem | 7,314 | 4,054 | 3,26 | 4,096 | 3 | 3 | 1,125 | 24 | No | ns | 0,9995 |
| Motor cortex vs. Striatum | 7,314 | 5,664 | 1,649 | 4,096 | 3 | 3 | 0,5694 | 24 | No | ns | > 0,9999 |
| Motor cortex vs. Claustrum | 7,314 | 8,697 | -1,384 | 4,096 | 3 | 3 | 0,4777 | 24 | No | ns | > 0,9999 |
| Association cortex vs. Prefrontal cortex | 7,668 | 6,124 | 1,543 | 4,096 | 3 | 3 | 0,5328 | 24 | No | ns | > 0,9999 |
| Association cortex vs. Olfactory regions | 7,668 | 3,755 | 3,912 | 4,096 | 3 | 3 | 1,351 | 24 | No | ns | 0,9974 |
| Association cortex vs. Hippocampus | 7,668 | 0,07276 | 7,595 | 4,096 | 3 | 3 | 2,622 | 24 | No | ns | 0,7741 |
| Association cortex vs. Thalamus | 7,668 | 47,44 | -39,77 | 4,096 | 3 | 3 | 13,73 | 24 | Yes | **** | < 0,0001 |
| Association cortex vs. Amygdala | 7,668 | 4,454 | 3,214 | 4,096 | 3 | 3 | 1,11 | 24 | No | ns | 0,9996 |
| Association cortex vs. Hypothalamus | 7,668 | 0,4891 | 7,179 | 4,096 | 3 | 3 | 2,478 | 24 | No | ns | 0,8269 |
| Association cortex vs. Brainstem | 7,668 | 4,054 | 3,614 | 4,096 | 3 | 3 | 1,248 | 24 | No | ns | 0,9987 |
| Association cortex vs. Striatum | 7,668 | 5,664 | 2,003 | 4,096 | 3 | 3 | 0,6916 | 24 | No | ns | > 0,9999 |
| Association cortex vs. Claustrum | 7,668 | 8,697 | -1,03 | 4,096 | 3 | 3 | 0,3555 | 24 | No | ns | > 0,9999 |
| Prefrontal cortex vs. Olfactory regions | 6,124 | 3,755 | 2,369 | 4,096 | 3 | 3 | 0,8179 | 24 | No | ns | > 0,9999 |
| Prefrontal cortex vs. Hippocampus | 6,124 | 0,07276 | 6,052 | 4,096 | 3 | 3 | 2,089 | 24 | No | ns | 0,933 |
| Prefrontal cortex vs. Thalamus | 6,124 | 47,44 | -41,31 | 4,096 | 3 | 3 | 14,26 | 24 | Yes | **** | < 0,0001 |
| Prefrontal cortex vs. Amygdala | 6,124 | 4,454 | 1,67 | 4,096 | 3 | 3 | 0,5767 | 24 | No | ns | > 0,9999 |
| Prefrontal cortex vs. Hypothalamus | 6,124 | 0,4891 | 5,635 | 4,096 | 3 | 3 | 1,946 | 24 | No | ns | 0,9576 |
| Prefrontal cortex vs. Brainstem | 6,124 | 4,054 | 2,071 | 4,096 | 3 | 3 | 0,7149 | 24 | No | ns | > 0,9999 |
| Prefrontal cortex vs. Striatum | 6,124 | 5,664 | 0,4599 | 4,096 | 3 | 3 | 0,1588 | 24 | No | ns | > 0,9999 |
| Prefrontal cortex vs. Claustrum | 6,124 | 8,697 | -2,573 | 4,096 | 3 | 3 | 0,8883 | 24 | No | ns | > 0,9999 |
| Olfactory regions vs. Hippocampus | 3,755 | 0,07276 | 3,683 | 4,096 | 3 | 3 | 1,271 | 24 | No | ns | 0,9985 |
| Olfactory regions vs. Thalamus | 3,755 | 47,44 | -43,68 | 4,096 | 3 | 3 | 15,08 | 24 | Yes | **** | < 0,0001 |
| Olfactory regions vs. Amygdala | 3,755 | 4,454 | -0,6986 | 4,096 | 3 | 3 | 0,2412 | 24 | No | ns | > 0,9999 |
| Olfactory regions vs. Hypothalamus | 3,755 | 0,4891 | 3,266 | 4,096 | 3 | 3 | 1,128 | 24 | No | ns | 0,9995 |
| Olfactory regions vs. Brainstem | 3,755 | 4,054 | -0,2984 | 4,096 | 3 | 3 | 0,103 | 24 | No | ns | > 0,9999 |
| Olfactory regions vs. Striatum | 3,755 | 5,664 | -1,909 | 4,096 | 3 | 3 | 0,6591 | 24 | No | ns | > 0,9999 |
| Olfactory regions vs. Claustrum | 3,755 | 8,697 | -4,942 | 4,096 | 3 | 3 | 1,706 | 24 | No | ns | 0,9832 |
| Hippocampus vs. Thalamus | 0,07276 | 47,44 | -47,36 | 4,096 | 3 | 3 | 16,35 | 24 | Yes | **** | < 0,0001 |
| Hippocampus vs. Amygdala | 0,07276 | 4,454 | -4,381 | 4,096 | 3 | 3 | 1,513 | 24 | No | ns | 0,9934 |
| Hippocampus vs. Hypothalamus | 0,07276 | 0,4891 | -0,4164 | 4,096 | 3 | 3 | 0,1438 | 24 | No | ns | > 0,9999 |
| Hippocampus vs. Brainstem | 0,07276 | 4,054 | -3,981 | 4,096 | 3 | 3 | 1,374 | 24 | No | ns | 0,997 |
| Hippocampus vs. Striatum | 0,07276 | 5,664 | -5,592 | 4,096 | 3 | 3 | 1,931 | 24 | No | ns | 0,9597 |
| Hippocampus vs. Claustrum | 0,07276 | 8,697 | -8,625 | 4,096 | 3 | 3 | 2,978 | 24 | No | ns | 0,624 |
| Thalamus vs. Amygdala | 47,44 | 4,454 | 42,98 | 4,096 | 3 | 3 | 14,84 | 24 | Yes | **** | < 0,0001 |
| Thalamus vs. Hypothalamus | 47,44 | 0,4891 | 46,95 | 4,096 | 3 | 3 | 16,21 | 24 | Yes | **** | < 0,0001 |
| Thalamus vs. Brainstem | 47,44 | 4,054 | 43,38 | 4,096 | 3 | 3 | 14,98 | 24 | Yes | **** | < 0,0001 |
| Thalamus vs. Striatum | 47,44 | 5,664 | 41,77 | 4,096 | 3 | 3 | 14,42 | 24 | Yes | **** | < 0,0001 |
| Thalamus vs. Claustrum | 47,44 | 8,697 | 38,74 | 4,096 | 3 | 3 | 13,37 | 24 | Yes | **** | < 0,0001 |
| Amygdala vs. Hypothalamus | 4,454 | 0,4891 | 3,965 | 4,096 | 3 | 3 | 1,369 | 24 | No | ns | 0,9971 |
| Amygdala vs. Brainstem | 4,454 | 4,054 | 0,4002 | 4,096 | 3 | 3 | 0,1382 | 24 | No | ns | > 0,9999 |
| Amygdala vs. Striatum | 4,454 | 5,664 | -1,211 | 4,096 | 3 | 3 | 0,4179 | 24 | No | ns | > 0,9999 |
| Amygdala vs. Claustrum | 4,454 | 8,697 | -4,244 | 4,096 | 3 | 3 | 1,465 | 24 | No | ns | 0,9949 |
| Hypothalamus vs. Brainstem | 0,4891 | 4,054 | -3,565 | 4,096 | 3 | 3 | 1,231 | 24 | No | ns | 0,9989 |
| Hypothalamus vs. Striatum | 0,4891 | 5,664 | -5,175 | 4,096 | 3 | 3 | 1,787 | 24 | No | ns | 0,9764 |
| Hypothalamus vs. Claustrum | 0,4891 | 8,697 | -8,208 | 4,096 | 3 | 3 | 2,834 | 24 | No | ns | 0,687 |
| Brainstem vs. Striatum | 4,054 | 5,664 | -1,611 | 4,096 | 3 | 3 | 0,5561 | 24 | No | ns | > 0,9999 |
| Brainstem vs. Claustrum | 4,054 | 8,697 | -4,644 | 4,096 | 3 | 3 | 1,603 | 24 | No | ns | 0,9895 |
| Striatum vs. Claustrum | 5,664 | 8,697 | -3,033 | 4,096 | 3 | 3 | 1,047 | 24 | No | ns | 0,9997 |

| Table Analyzed | AI inputs |
|---|---|
| **ANOVA summary** | |
| F | 3,911 |
| P value | 0,0025 |
| P value summary | ** |
| Are differences among means statistically sig | Yes |
| R square | 0,6419 |

**Tukey's multiple comparisons test**

| | Mean 1 | Mean 2 | Mean Diff, | SE of diff, | n1 | n2 | q | DF | Significant | Summary | Adjusted P Value |
|---|---|---|---|---|---|---|---|---|---|---|---|
| Sensory Cortex vs. Motor cortex | 14,72 | 1,22 | 13,5 | 4,125 | 3 | 3 | 4,63 | 24 | No | ns | 0,0998 |
| Sensory Cortex vs. Association cortex | 14,72 | 12,88 | 1,847 | 4,125 | 3 | 3 | 0,6334 | 24 | No | ns | > 0,9999 |
| Sensory Cortex vs. Prefrontal cortex | 14,72 | 5,658 | 9,067 | 4,125 | 3 | 3 | 3,108 | 24 | No | ns | 0,5658 |
| Sensory Cortex vs. Olfactory regions | 14,72 | 12,24 | 2,487 | 4,125 | 3 | 3 | 0,8526 | 24 | No | ns | > 0,9999 |
| Sensory Cortex vs. Hippocampus | 14,72 | 0,06216 | 14,66 | 4,125 | 3 | 3 | 5,027 | 24 | No | ns | 0,0558 |
| Sensory Cortex vs. Thalamus | 14,72 | 17,77 | -3,047 | 4,125 | 3 | 3 | 1,045 | 24 | No | ns | 0,9997 |
| Sensory Cortex vs. Amygdala | 14,72 | 9,187 | 5,537 | 4,125 | 3 | 3 | 1,898 | 24 | No | ns | 0,964 |
| Sensory Cortex vs. Hypothalamus | 14,72 | 0,2671 | 14,46 | 4,125 | 3 | 3 | 4,956 | 24 | No | ns | 0,062 |
| Sensory Cortex vs. Brainstem | 14,72 | 8,76 | 5,964 | 4,125 | 3 | 3 | 2,045 | 24 | No | ns | 0,9415 |
| Sensory Cortex vs. Striatum | 14,72 | 6,63 | 8,095 | 4,125 | 3 | 3 | 2,775 | 24 | No | ns | 0,7121 |
| Sensory Cortex vs. Claustrum | 14,72 | 10,61 | 4,118 | 4,125 | 3 | 3 | 1,412 | 24 | No | ns | 0,9963 |
| Motor cortex vs. Association cortex | 1,22 | 12,88 | -11,66 | 4,125 | 3 | 3 | 3,996 | 24 | No | ns | 0,2303 |
| Motor cortex vs. Prefrontal cortex | 1,22 | 5,658 | -4,438 | 4,125 | 3 | 3 | 1,521 | 24 | No | ns | 0,9931 |
| Motor cortex vs. Olfactory regions | 1,22 | 12,24 | -11,02 | 4,125 | 3 | 3 | 3,777 | 24 | No | ns | 0,2975 |
| Motor cortex vs. Hippocampus | 1,22 | 0,06216 | 1,158 | 4,125 | 3 | 3 | 0,3969 | 24 | No | ns | > 0,9999 |
| Motor cortex vs. Thalamus | 1,22 | 17,77 | -16,55 | 4,125 | 3 | 3 | 5,674 | 24 | Yes | * | 0,0203 |
| Motor cortex vs. Amygdala | 1,22 | 9,187 | -7,967 | 4,125 | 3 | 3 | 2,731 | 24 | No | ns | 0,7303 |
| Motor cortex vs. Hypothalamus | 1,22 | 0,2671 | 0,9527 | 4,125 | 3 | 3 | 0,3266 | 24 | No | ns | > 0,9999 |
| Motor cortex vs. Brainstem | 1,22 | 8,76 | -7,54 | 4,125 | 3 | 3 | 2,585 | 24 | No | ns | 0,7884 |
| Motor cortex vs. Striatum | 1,22 | 6,63 | -5,41 | 4,125 | 3 | 3 | 1,855 | 24 | No | ns | 0,9694 |
| Motor cortex vs. Claustrum | 1,22 | 10,61 | -9,386 | 4,125 | 3 | 3 | 3,218 | 24 | No | ns | 0,5173 |
| Association cortex vs. Prefrontal cortex | 12,88 | 5,658 | 7,219 | 4,125 | 3 | 3 | 2,475 | 24 | No | ns | 0,8281 |
| Association cortex vs. Olfactory regions | 12,88 | 12,24 | 0,6394 | 4,125 | 3 | 3 | 0,2192 | 24 | No | ns | > 0,9999 |
| Association cortex vs. Hippocampus | 12,88 | 0,06216 | 12,81 | 4,125 | 3 | 3 | 4,393 | 24 | No | ns | 0,1384 |
| Association cortex vs. Thalamus | 12,88 | 17,77 | -4,895 | 4,125 | 3 | 3 | 1,678 | 24 | No | ns | 0,9851 |
| Association cortex vs. Amygdala | 12,88 | 9,187 | 3,69 | 4,125 | 3 | 3 | 1,265 | 24 | No | ns | 0,9985 |
| Association cortex vs. Hypothalamus | 12,88 | 0,2671 | 12,61 | 4,125 | 3 | 3 | 4,323 | 24 | No | ns | 0,1521 |
| Association cortex vs. Brainstem | 12,88 | 8,76 | 4,117 | 4,125 | 3 | 3 | 1,411 | 24 | No | ns | 0,9963 |
| Association cortex vs. Striatum | 12,88 | 6,63 | 6,247 | 4,125 | 3 | 3 | 2,142 | 24 | No | ns | 0,9221 |
| Association cortex vs. Claustrum | 12,88 | 10,61 | 2,27 | 4,125 | 3 | 3 | 0,7784 | 24 | No | ns | > 0,9999 |
| Prefrontal cortex vs. Olfactory regions | 5,658 | 12,24 | -6,58 | 4,125 | 3 | 3 | 2,256 | 24 | No | ns | 0,8947 |
| Prefrontal cortex vs. Hippocampus | 5,658 | 0,06216 | 5,596 | 4,125 | 3 | 3 | 1,918 | 24 | No | ns | 0,9614 |
| Prefrontal cortex vs. Thalamus | 5,658 | 17,77 | -12,11 | 4,125 | 3 | 3 | 4,153 | 24 | No | ns | 0,1896 |
| Prefrontal cortex vs. Amygdala | 5,658 | 9,187 | -3,529 | 4,125 | 3 | 3 | 1,21 | 24 | No | ns | 0,999 |
| Prefrontal cortex vs. Hypothalamus | 5,658 | 0,2671 | 5,391 | 4,125 | 3 | 3 | 1,848 | 24 | No | ns | 0,9701 |
| Prefrontal cortex vs. Brainstem | 5,658 | 8,76 | -3,102 | 4,125 | 3 | 3 | 1,064 | 24 | No | ns | 0,9997 |
| Prefrontal cortex vs. Striatum | 5,658 | 6,63 | -0,9719 | 4,125 | 3 | 3 | 0,3332 | 24 | No | ns | > 0,9999 |
| Prefrontal cortex vs. Claustrum | 5,658 | 10,61 | -4,949 | 4,125 | 3 | 3 | 1,696 | 24 | No | ns | 0,9839 |
| Olfactory regions vs. Hippocampus | 12,24 | 0,06216 | 12,18 | 4,125 | 3 | 3 | 4,174 | 24 | No | ns | 0,1846 |
| Olfactory regions vs. Thalamus | 12,24 | 17,77 | -5,534 | 4,125 | 3 | 3 | 1,897 | 24 | No | ns | 0,9642 |
| Olfactory regions vs. Amygdala | 12,24 | 9,187 | 3,05 | 4,125 | 3 | 3 | 1,046 | 24 | No | ns | 0,9997 |
| Olfactory regions vs. Hypothalamus | 12,24 | 0,2671 | 11,97 | 4,125 | 3 | 3 | 4,104 | 24 | No | ns | 0,2018 |
| Olfactory regions vs. Brainstem | 12,24 | 8,76 | 3,477 | 4,125 | 3 | 3 | 1,192 | 24 | No | ns | 0,9991 |
| Olfactory regions vs. Striatum | 12,24 | 6,63 | 5,608 | 4,125 | 3 | 3 | 1,922 | 24 | No | ns | 0,9608 |
| Olfactory regions vs. Claustrum | 12,24 | 10,61 | 1,631 | 4,125 | 3 | 3 | 0,5592 | 24 | No | ns | > 0,9999 |
| Hippocampus vs. Thalamus | 0,06216 | 17,77 | -17,71 | 4,125 | 3 | 3 | 6,071 | 24 | Yes | * | 0,0106 |
| Hippocampus vs. Amygdala | 0,06216 | 9,187 | -9,125 | 4,125 | 3 | 3 | 3,128 | 24 | No | ns | 0,5569 |
| Hippocampus vs. Hypothalamus | 0,06216 | 0,2671 | -0,205 | 4,125 | 3 | 3 | 0,0703 | 24 | No | ns | > 0,9999 |
| Hippocampus vs. Brainstem | 0,06216 | 8,76 | -8,698 | 4,125 | 3 | 3 | 2,982 | 24 | No | ns | 0,6221 |
| Hippocampus vs. Striatum | 0,06216 | 6,63 | -6,567 | 4,125 | 3 | 3 | 2,251 | 24 | No | ns | 0,8958 |
| Hippocampus vs. Claustrum | 0,06216 | 10,61 | -10,54 | 4,125 | 3 | 3 | 3,615 | 24 | No | ns | 0,3548 |
| Thalamus vs. Amygdala | 17,77 | 9,187 | 8,584 | 4,125 | 3 | 3 | 2,943 | 24 | No | ns | 0,6393 |
| Thalamus vs. Hypothalamus | 17,77 | 0,2671 | 17,5 | 4,125 | 3 | 3 | 6,001 | 24 | Yes | * | 0,0119 |
| Thalamus vs. Brainstem | 17,77 | 8,76 | 9,012 | 4,125 | 3 | 3 | 3,089 | 24 | No | ns | 0,5742 |
| Thalamus vs. Striatum | 17,77 | 6,63 | 11,14 | 4,125 | 3 | 3 | 3,82 | 24 | No | ns | 0,2834 |
| Thalamus vs. Claustrum | 17,77 | 10,61 | 7,165 | 4,125 | 3 | 3 | 2,456 | 24 | No | ns | 0,8344 |
| Amygdala vs. Hypothalamus | 9,187 | 0,2671 | 8,92 | 4,125 | 3 | 3 | 3,058 | 24 | No | ns | 0,5882 |
| Amygdala vs. Brainstem | 9,187 | 8,76 | 0,4272 | 4,125 | 3 | 3 | 0,1465 | 24 | No | ns | > 0,9999 |
| Amygdala vs. Striatum | 9,187 | 6,63 | 2,558 | 4,125 | 3 | 3 | 0,8768 | 24 | No | ns | > 0,9999 |
| Amygdala vs. Claustrum | 9,187 | 10,61 | -1,419 | 4,125 | 3 | 3 | 0,4865 | 24 | No | ns | > 0,9999 |
| Hypothalamus vs. Brainstem | 0,2671 | 8,76 | -8,493 | 4,125 | 3 | 3 | 2,912 | 24 | No | ns | 0,6532 |
| Hypothalamus vs. Striatum | 0,2671 | 6,63 | -6,363 | 4,125 | 3 | 3 | 2,181 | 24 | No | ns | 0,9132 |
| Hypothalamus vs. Claustrum | 0,2671 | 10,61 | -10,34 | 4,125 | 3 | 3 | 3,545 | 24 | No | ns | 0,3814 |
| Brainstem vs. Striatum | 8,76 | 6,63 | 2,13 | 4,125 | 3 | 3 | 0,7303 | 24 | No | ns | > 0,9999 |
| Brainstem vs. Claustrum | 8,76 | 10,61 | -1,846 | 4,125 | 3 | 3 | 0,633 | 24 | No | ns | > 0,9999 |
| Striatum vs. Claustrum | 6,63 | 10,61 | -3,977 | 4,125 | 3 | 3 | 1,363 | 24 | No | ns | 0,9972 |

**Table Analyzed** **DLO inputs**

**ANOVA summary**

| | |
|---|---|
| F | 7,882 |
| P value | < 0,0001 |
| P value summary | **** |
| Are differences among means statistically s | Yes |
| R square | 0,7832 |

**Tukey's multiple comparisons test**

| | Mean 1 | Mean 2 | Mean Diff, | SE of diff, | n1 | n2 | q | DF | Significant | Summary | Adjusted P Value |
|---|---|---|---|---|---|---|---|---|---|---|---|
| Sensory Cortex vs. Motor cortex | 4,761 | 7,96 | -3,2 | 4,633 | 3 | 3 | 0,9767 | 24 | No | ns | 0,9999 |
| Sensory Cortex vs. Association cortex | 4,761 | 8,911 | -4,151 | 4,633 | 3 | 3 | 1,267 | 24 | No | ns | 0,9985 |
| Sensory Cortex vs. Prefrontal cortex | 4,761 | 7,09 | -2,329 | 4,633 | 3 | 3 | 0,711 | 24 | No | ns | > 0,9999 |
| Sensory Cortex vs. Olfactory regions | 4,761 | 6,058 | -1,297 | 4,633 | 3 | 3 | 0,396 | 24 | No | ns | > 0,9999 |
| Sensory Cortex vs. Hippocampus | 4,761 | 0 | 4,761 | 4,633 | 3 | 3 | 1,453 | 24 | No | ns | 0,9952 |
| Sensory Cortex vs. Thalamus | 4,761 | 34,73 | -29,97 | 4,633 | 3 | 3 | 9,149 | 24 | Yes | **** | < 0,0001 |
| Sensory Cortex vs. Amygdala | 4,761 | 9,563 | -4,802 | 4,633 | 3 | 3 | 1,466 | 24 | No | ns | 0,9949 |
| Sensory Cortex vs. Hypothalamus | 4,761 | 0,0214 | 4,739 | 4,633 | 3 | 3 | 1,447 | 24 | No | ns | 0,9954 |
| Sensory Cortex vs. Brainstem | 4,761 | 0,9867 | 3,774 | 4,633 | 3 | 3 | 1,152 | 24 | No | ns | 0,9994 |
| Sensory Cortex vs. Striatum | 4,761 | 7,283 | -2,523 | 4,633 | 3 | 3 | 0,7701 | 24 | No | ns | > 0,9999 |
| Sensory Cortex vs. Claustrum | 4,761 | 12,64 | -7,874 | 4,633 | 3 | 3 | 2,404 | 24 | No | ns | 0,8516 |
| Motor cortex vs. Association cortex | 7,96 | 8,911 | -0,9512 | 4,633 | 3 | 3 | 0,2904 | 24 | No | ns | > 0,9999 |
| Motor cortex vs. Prefrontal cortex | 7,96 | 7,09 | 0,8705 | 4,633 | 3 | 3 | 0,2657 | 24 | No | ns | > 0,9999 |
| Motor cortex vs. Olfactory regions | 7,96 | 6,058 | 1,902 | 4,633 | 3 | 3 | 0,5807 | 24 | No | ns | > 0,9999 |
| Motor cortex vs. Hippocampus | 7,96 | 0 | 7,96 | 4,633 | 3 | 3 | 2,43 | 24 | No | ns | 0,8432 |
| Motor cortex vs. Thalamus | 7,96 | 34,73 | -26,77 | 4,633 | 3 | 3 | 8,172 | 24 | Yes | *** | 0,0003 |
| Motor cortex vs. Amygdala | 7,96 | 9,563 | -1,603 | 4,633 | 3 | 3 | 0,4892 | 24 | No | ns | > 0,9999 |
| Motor cortex vs. Hypothalamus | 7,96 | 0,0214 | 7,939 | 4,633 | 3 | 3 | 2,423 | 24 | No | ns | 0,8453 |
| Motor cortex vs. Brainstem | 7,96 | 0,9867 | 6,974 | 4,633 | 3 | 3 | 2,129 | 24 | No | ns | 0,9249 |
| Motor cortex vs. Striatum | 7,96 | 7,283 | 0,6769 | 4,633 | 3 | 3 | 0,2066 | 24 | No | ns | > 0,9999 |
| Motor cortex vs. Claustrum | 7,96 | 12,64 | -4,675 | 4,633 | 3 | 3 | 1,427 | 24 | No | ns | 0,9959 |
| Association cortex vs. Prefrontal cortex | 8,911 | 7,09 | 1,822 | 4,633 | 3 | 3 | 0,5561 | 24 | No | ns | > 0,9999 |
| Association cortex vs. Olfactory regions | 8,911 | 6,058 | 2,853 | 4,633 | 3 | 3 | 0,871 | 24 | No | ns | > 0,9999 |
| Association cortex vs. Hippocampus | 8,911 | 0 | 8,911 | 4,633 | 3 | 3 | 2,72 | 24 | No | ns | 0,7349 |
| Association cortex vs. Thalamus | 8,911 | 34,73 | -25,82 | 4,633 | 3 | 3 | 7,882 | 24 | Yes | *** | 0,0005 |
| Association cortex vs. Amygdala | 8,911 | 9,563 | -0,6513 | 4,633 | 3 | 3 | 0,1988 | 24 | No | ns | > 0,9999 |
| Association cortex vs. Hypothalamus | 8,911 | 0,0214 | 8,89 | 4,633 | 3 | 3 | 2,714 | 24 | No | ns | 0,7375 |
| Association cortex vs. Brainstem | 8,911 | 0,9867 | 7,925 | 4,633 | 3 | 3 | 2,419 | 24 | No | ns | 0,8467 |
| Association cortex vs. Striatum | 8,911 | 7,283 | 1,628 | 4,633 | 3 | 3 | 0,497 | 24 | No | ns | > 0,9999 |
| Association cortex vs. Claustrum | 8,911 | 12,64 | -3,724 | 4,633 | 3 | 3 | 1,137 | 24 | No | ns | 0,9994 |
| Prefrontal cortex vs. Olfactory regions | 7,09 | 6,058 | 1,032 | 4,633 | 3 | 3 | 0,315 | 24 | No | ns | > 0,9999 |
| Prefrontal cortex vs. Hippocampus | 7,09 | 0 | 7,09 | 4,633 | 3 | 3 | 2,164 | 24 | No | ns | 0,9171 |
| Prefrontal cortex vs. Thalamus | 7,09 | 34,73 | -27,64 | 4,633 | 3 | 3 | 8,438 | 24 | Yes | *** | 0,0002 |
| Prefrontal cortex vs. Amygdala | 7,09 | 9,563 | -2,473 | 4,633 | 3 | 3 | 0,7549 | 24 | No | ns | > 0,9999 |
| Prefrontal cortex vs. Hypothalamus | 7,09 | 0,0214 | 7,068 | 4,633 | 3 | 3 | 2,158 | 24 | No | ns | 0,9186 |
| Prefrontal cortex vs. Brainstem | 7,09 | 0,9867 | 6,103 | 4,633 | 3 | 3 | 1,863 | 24 | No | ns | 0,9684 |
| Prefrontal cortex vs. Striatum | 7,09 | 7,283 | -0,1936 | 4,633 | 3 | 3 | 0,0591 | 24 | No | ns | > 0,9999 |
| Prefrontal cortex vs. Claustrum | 7,09 | 12,64 | -5,545 | 4,633 | 3 | 3 | 1,693 | 24 | No | ns | 0,9841 |
| Olfactory regions vs. Hippocampus | 6,058 | 0 | 6,058 | 4,633 | 3 | 3 | 1,849 | 24 | No | ns | 0,97 |
| Olfactory regions vs. Thalamus | 6,058 | 34,73 | -28,67 | 4,633 | 3 | 3 | 8,753 | 24 | Yes | *** | 0,0001 |
| Olfactory regions vs. Amygdala | 6,058 | 9,563 | -3,505 | 4,633 | 3 | 3 | 1,07 | 24 | No | ns | 0,9997 |
| Olfactory regions vs. Hypothalamus | 6,058 | 0,0214 | 6,037 | 4,633 | 3 | 3 | 1,843 | 24 | No | ns | 0,9707 |
| Olfactory regions vs. Brainstem | 6,058 | 0,9867 | 5,071 | 4,633 | 3 | 3 | 1,548 | 24 | No | ns | 0,992 |
| Olfactory regions vs. Striatum | 6,058 | 7,283 | -1,225 | 4,633 | 3 | 3 | 0,3741 | 24 | No | ns | > 0,9999 |
| Olfactory regions vs. Claustrum | 6,058 | 12,64 | -6,577 | 4,633 | 3 | 3 | 2,008 | 24 | No | ns | 0,9479 |
| Hippocampus vs. Thalamus | 0 | 34,73 | -34,73 | 4,633 | 3 | 3 | 10,6 | 24 | Yes | **** | < 0,0001 |
| Hippocampus vs. Amygdala | 0 | 9,563 | -9,563 | 4,633 | 3 | 3 | 2,919 | 24 | No | ns | 0,6498 |
| Hippocampus vs. Hypothalamus | 0 | 0,0214 | -0,0214 | 4,633 | 3 | 3 | 0,0065 | 24 | No | ns | > 0,9999 |
| Hippocampus vs. Brainstem | 0 | 0,9867 | -0,9867 | 4,633 | 3 | 3 | 0,3012 | 24 | No | ns | > 0,9999 |
| Hippocampus vs. Striatum | 0 | 7,283 | -7,283 | 4,633 | 3 | 3 | 2,223 | 24 | No | ns | 0,903 |
| Hippocampus vs. Claustrum | 0 | 12,64 | -12,64 | 4,633 | 3 | 3 | 3,857 | 24 | No | ns | 0,2716 |
| Thalamus vs. Amygdala | 34,73 | 9,563 | 25,17 | 4,633 | 3 | 3 | 7,683 | 24 | Yes | *** | 0,0007 |
| Thalamus vs. Hypothalamus | 34,73 | 0,0214 | 34,71 | 4,633 | 3 | 3 | 10,6 | 24 | Yes | **** | < 0,0001 |
| Thalamus vs. Brainstem | 34,73 | 0,9867 | 33,74 | 4,633 | 3 | 3 | 10,3 | 24 | Yes | **** | < 0,0001 |
| Thalamus vs. Striatum | 34,73 | 7,283 | 27,45 | 4,633 | 3 | 3 | 8,379 | 24 | Yes | *** | 0,0002 |
| Thalamus vs. Claustrum | 34,73 | 12,64 | 22,1 | 4,633 | 3 | 3 | 6,745 | 24 | Yes | ** | 0,0034 |
| Amygdala vs. Hypothalamus | 9,563 | 0,0214 | 9,541 | 4,633 | 3 | 3 | 2,913 | 24 | No | ns | 0,6527 |
| Amygdala vs. Brainstem | 9,563 | 0,9867 | 8,576 | 4,633 | 3 | 3 | 2,618 | 24 | No | ns | 0,7757 |
| Amygdala vs. Striatum | 9,563 | 7,283 | 2,279 | 4,633 | 3 | 3 | 0,6958 | 24 | No | ns | > 0,9999 |
| Amygdala vs. Claustrum | 9,563 | 12,64 | -3,072 | 4,633 | 3 | 3 | 0,9378 | 24 | No | ns | > 0,9999 |
| Hypothalamus vs. Brainstem | 0,0214 | 0,9867 | -0,9653 | 4,633 | 3 | 3 | 0,2947 | 24 | No | ns | > 0,9999 |
| Hypothalamus vs. Striatum | 0,0214 | 7,283 | -7,262 | 4,633 | 3 | 3 | 2,217 | 24 | No | ns | 0,9047 |
| Hypothalamus vs. Claustrum | 0,0214 | 12,64 | -12,61 | 4,633 | 3 | 3 | 3,85 | 24 | No | ns | 0,2736 |
| Brainstem vs. Striatum | 0,9867 | 7,283 | -6,297 | 4,633 | 3 | 3 | 1,922 | 24 | No | ns | 0,9609 |
| Brainstem vs. Claustrum | 0,9867 | 12,64 | -11,65 | 4,633 | 3 | 3 | 3,556 | 24 | No | ns | 0,3771 |
| Striatum vs. Claustrum | 7,283 | 12,64 | -5,352 | 4,633 | 3 | 3 | 1,634 | 24 | No | ns | 0,9879 |

| Table Analyzed | MO outputs |
|---|---|

**ANOVA summary**

| | |
|---|---|
| F | 6,955 |
| P value | < 0,0001 |
| P value summary | **** |
| Are differences among means statistically significant? (P < ( | Yes |
| R square | 0,7612 |

**Tukey's multiple comparisons test**

| | Mean 1 | Mean 2 | Mean Diff, | SE of diff, | n1 | n2 | q | DF | Significant | Summary | Adjusted P |
|---|---|---|---|---|---|---|---|---|---|---|---|
| Sensory Cortex vs. Motor cortex | 8,934 | 0,9539 | 7,98 | 4,784 | 3 | 3 | 2,359 | 24 | No | ns | 0,8655 |
| Sensory Cortex vs. Association cortex | 8,934 | 11,44 | -2,509 | 4,784 | 3 | 3 | 0,7415 | 24 | No | ns | > 0,9999 |
| Sensory Cortex vs. Prefrontal cortex | 8,934 | 30,24 | -21,31 | 4,784 | 3 | 3 | 6,299 | 24 | Yes | ** | 0,0072 |
| Sensory Cortex vs. Olfactory regions | 8,934 | 2,868 | 6,066 | 4,784 | 3 | 3 | 1,793 | 24 | No | ns | 0,9758 |
| Sensory Cortex vs. Hippocampus | 8,934 | 0,03402 | 8,9 | 4,784 | 3 | 3 | 2,631 | 24 | No | ns | 0,7707 |
| Sensory Cortex vs. Thalamus | 8,934 | 20,59 | -11,66 | 4,784 | 3 | 3 | 3,447 | 24 | No | ns | 0,4203 |
| Sensory Cortex vs. Amygdala | 8,934 | 3,472 | 5,462 | 4,784 | 3 | 3 | 1,615 | 24 | No | ns | 0,9889 |
| Sensory Cortex vs. Hypothalamus | 8,934 | 3,058 | 5,876 | 4,784 | 3 | 3 | 1,737 | 24 | No | ns | 0,9808 |
| Sensory Cortex vs. Brainstem | 8,934 | 3,091 | 5,844 | 4,784 | 3 | 3 | 1,727 | 24 | No | ns | 0,9816 |
| Sensory Cortex vs. Striatum | 8,934 | 8,653 | 0,2811 | 4,784 | 3 | 3 | 0,08309 | 24 | No | ns | > 0,9999 |
| Sensory Cortex vs. Claustrum | 8,934 | 6,655 | 2,279 | 4,784 | 3 | 3 | 0,6737 | 24 | No | ns | > 0,9999 |
| Motor cortex vs. Association cortex | 0,9539 | 11,44 | -10,49 | 4,784 | 3 | 3 | 3,101 | 24 | No | ns | 0,5692 |
| Motor cortex vs. Prefrontal cortex | 0,9539 | 30,24 | -29,29 | 4,784 | 3 | 3 | 8,658 | 24 | Yes | *** | 0,0001 |
| Motor cortex vs. Olfactory regions | 0,9539 | 2,868 | -1,914 | 4,784 | 3 | 3 | 0,5658 | 24 | No | ns | > 0,9999 |
| Motor cortex vs. Hippocampus | 0,9539 | 0,03402 | 0,9199 | 4,784 | 3 | 3 | 0,2719 | 24 | No | ns | > 0,9999 |
| Motor cortex vs. Thalamus | 0,9539 | 20,59 | -19,64 | 4,784 | 3 | 3 | 5,806 | 24 | Yes | * | 0,0164 |
| Motor cortex vs. Amygdala | 0,9539 | 3,472 | -2,519 | 4,784 | 3 | 3 | 0,7445 | 24 | No | ns | > 0,9999 |
| Motor cortex vs. Hypothalamus | 0,9539 | 3,058 | -2,104 | 4,784 | 3 | 3 | 0,6219 | 24 | No | ns | > 0,9999 |
| Motor cortex vs. Brainstem | 0,9539 | 3,091 | -2,137 | 4,784 | 3 | 3 | 0,6316 | 24 | No | ns | > 0,9999 |
| Motor cortex vs. Striatum | 0,9539 | 8,653 | -7,699 | 4,784 | 3 | 3 | 2,276 | 24 | No | ns | 0,8893 |
| Motor cortex vs. Claustrum | 0,9539 | 6,655 | -5,701 | 4,784 | 3 | 3 | 1,685 | 24 | No | ns | 0,9847 |
| Association cortex vs. Prefrontal cortex | 11,44 | 30,24 | -18,8 | 4,784 | 3 | 3 | 5,558 | 24 | Yes | * | 0,0244 |
| Association cortex vs. Olfactory regions | 11,44 | 2,868 | 8,575 | 4,784 | 3 | 3 | 2,535 | 24 | No | ns | 0,8069 |
| Association cortex vs. Hippocampus | 11,44 | 0,03402 | 11,41 | 4,784 | 3 | 3 | 3,372 | 24 | No | ns | 0,4509 |
| Association cortex vs. Thalamus | 11,44 | 20,59 | -9,152 | 4,784 | 3 | 3 | 2,705 | 24 | No | ns | 0,741 |
| Association cortex vs. Amygdala | 11,44 | 3,472 | 7,97 | 4,784 | 3 | 3 | 2,356 | 24 | No | ns | 0,8664 |
| Association cortex vs. Hypothalamus | 11,44 | 3,058 | 8,385 | 4,784 | 3 | 3 | 2,479 | 24 | No | ns | 0,8268 |
| Association cortex vs. Brainstem | 11,44 | 3,091 | 8,352 | 4,784 | 3 | 3 | 2,469 | 24 | No | ns | 0,8301 |
| Association cortex vs. Striatum | 11,44 | 8,653 | 2,79 | 4,784 | 3 | 3 | 0,8246 | 24 | No | ns | > 0,9999 |
| Association cortex vs. Claustrum | 11,44 | 6,655 | 4,787 | 4,784 | 3 | 3 | 1,415 | 24 | No | ns | 0,9962 |
| Prefrontal cortex vs. Olfactory regions | 30,24 | 2,868 | 27,38 | 4,784 | 3 | 3 | 8,092 | 24 | Yes | *** | 0,0003 |
| Prefrontal cortex vs. Hippocampus | 30,24 | 0,03402 | 30,21 | 4,784 | 3 | 3 | 8,93 | 24 | Yes | **** | < 0,0001 |
| Prefrontal cortex vs. Thalamus | 30,24 | 20,59 | 9,649 | 4,784 | 3 | 3 | 2,852 | 24 | No | ns | 0,679 |
| Prefrontal cortex vs. Amygdala | 30,24 | 3,472 | 26,77 | 4,784 | 3 | 3 | 7,914 | 24 | Yes | *** | 0,0005 |
| Prefrontal cortex vs. Hypothalamus | 30,24 | 3,058 | 27,19 | 4,784 | 3 | 3 | 8,036 | 24 | Yes | *** | 0,0004 |
| Prefrontal cortex vs. Brainstem | 30,24 | 3,091 | 27,15 | 4,784 | 3 | 3 | 8,027 | 24 | Yes | *** | 0,0004 |
| Prefrontal cortex vs. Striatum | 30,24 | 8,653 | 21,59 | 4,784 | 3 | 3 | 6,382 | 24 | Yes | ** | 0,0063 |
| Prefrontal cortex vs. Claustrum | 30,24 | 6,655 | 23,59 | 4,784 | 3 | 3 | 6,973 | 24 | Yes | ** | 0,0023 |
| Olfactory regions vs. Hippocampus | 2,868 | 0,03402 | 2,834 | 4,784 | 3 | 3 | 0,8377 | 24 | No | ns | > 0,9999 |
| Olfactory regions vs. Thalamus | 2,868 | 20,59 | -17,73 | 4,784 | 3 | 3 | 5,24 | 24 | Yes | * | 0,0403 |
| Olfactory regions vs. Amygdala | 2,868 | 3,472 | -0,6045 | 4,784 | 3 | 3 | 0,1787 | 24 | No | ns | > 0,9999 |
| Olfactory regions vs. Hypothalamus | 2,868 | 3,058 | -0,1899 | 4,784 | 3 | 3 | 0,05614 | 24 | No | ns | > 0,9999 |
| Olfactory regions vs. Brainstem | 2,868 | 3,091 | -0,2226 | 4,784 | 3 | 3 | 0,0658 | 24 | No | ns | > 0,9999 |
| Olfactory regions vs. Striatum | 2,868 | 8,653 | -5,785 | 4,784 | 3 | 3 | 1,71 | 24 | No | ns | 0,9829 |
| Olfactory regions vs. Claustrum | 2,868 | 6,655 | -3,787 | 4,784 | 3 | 3 | 1,12 | 24 | No | ns | 0,9995 |
| Hippocampus vs. Thalamus | 0,03402 | 20,59 | -20,56 | 4,784 | 3 | 3 | 6,078 | 24 | Yes | * | 0,0105 |
| Hippocampus vs. Amygdala | 0,03402 | 3,472 | -3,438 | 4,784 | 3 | 3 | 1,016 | 24 | No | ns | 0,9998 |
| Hippocampus vs. Hypothalamus | 0,03402 | 3,058 | -3,024 | 4,784 | 3 | 3 | 0,8939 | 24 | No | ns | > 0,9999 |
| Hippocampus vs. Brainstem | 0,03402 | 3,091 | -3,057 | 4,784 | 3 | 3 | 0,9035 | 24 | No | ns | > 0,9999 |
| Hippocampus vs. Striatum | 0,03402 | 8,653 | -8,619 | 4,784 | 3 | 3 | 2,548 | 24 | No | ns | 0,8022 |
| Hippocampus vs. Claustrum | 0,03402 | 6,655 | -6,621 | 4,784 | 3 | 3 | 1,957 | 24 | No | ns | 0,9559 |
| Thalamus vs. Amygdala | 20,59 | 3,472 | 17,12 | 4,784 | 3 | 3 | 5,061 | 24 | No | ns | 0,053 |
| Thalamus vs. Hypothalamus | 20,59 | 3,058 | 17,54 | 4,784 | 3 | 3 | 5,184 | 24 | Yes | * | 0,0439 |
| Thalamus vs. Brainstem | 20,59 | 3,091 | 17,5 | 4,784 | 3 | 3 | 5,174 | 24 | Yes | * | 0,0446 |
| Thalamus vs. Striatum | 20,59 | 8,653 | 11,94 | 4,784 | 3 | 3 | 3,53 | 24 | No | ns | 0,3871 |
| Thalamus vs. Claustrum | 20,59 | 6,655 | 13,94 | 4,784 | 3 | 3 | 4,12 | 24 | No | ns | 0,1976 |
| Amygdala vs. Hypothalamus | 3,472 | 3,058 | 0,4146 | 4,784 | 3 | 3 | 0,1226 | 24 | No | ns | > 0,9999 |
| Amygdala vs. Brainstem | 3,472 | 3,091 | 0,3819 | 4,784 | 3 | 3 | 0,1129 | 24 | No | ns | > 0,9999 |
| Amygdala vs. Striatum | 3,472 | 8,653 | -5,181 | 4,784 | 3 | 3 | 1,531 | 24 | No | ns | 0,9927 |
| Amygdala vs. Claustrum | 3,472 | 6,655 | -3,183 | 4,784 | 3 | 3 | 0,9409 | 24 | No | ns | > 0,9999 |
| Hypothalamus vs. Brainstem | 3,058 | 3,091 | -0,0327 | 4,784 | 3 | 3 | 0,009665 | 24 | No | ns | > 0,9999 |
| Hypothalamus vs. Striatum | 3,058 | 8,653 | -5,595 | 4,784 | 3 | 3 | 1,654 | 24 | No | ns | 0,9867 |
| Hypothalamus vs. Claustrum | 3,058 | 6,655 | -3,597 | 4,784 | 3 | 3 | 1,063 | 24 | No | ns | 0,9997 |
| Brainstem vs. Striatum | 3,091 | 8,653 | -5,563 | 4,784 | 3 | 3 | 1,644 | 24 | No | ns | 0,9873 |
| Brainstem vs. Claustrum | 3,091 | 6,655 | -3,565 | 4,784 | 3 | 3 | 1,054 | 24 | No | ns | 0,9997 |
| Striatum vs. Claustrum | 8,653 | 6,655 | 1,998 | 4,784 | 3 | 3 | 0,5906 | 24 | No | ns | > 0,9999 |

| Table Analyzed | VO outputs |
|---|---|

**ANOVA summary**

| | |
|---|---|
| F | 4,523 |
| P value | 0,0003 |
| P value summary | *** |
| Are differences among means statistically significant? (P < ( | Yes |
| R square | 0,5802 |

**Tukey's multiple comparisons test**

| | Mean 1 | Mean 2 | Mean Diff, | SE of diff, | n1 | n2 | q | DF | Significant | Summary | Adjusted P |
|---|---|---|---|---|---|---|---|---|---|---|---|
| Sensory Cortex vs. Motor cortex | 26,19 | 1,423 | 24,77 | 6,117 | 4 | 4 | 5,726 | 36 | Yes | * | 0,0119 |
| Sensory Cortex vs. Association cortex | 26,19 | 6,064 | 20,13 | 6,117 | 4 | 4 | 4,653 | 36 | No | ns | 0,08 |
| Sensory Cortex vs. Prefrontal cortex | 26,19 | 10,3 | 15,89 | 6,117 | 4 | 4 | 3,673 | 36 | No | ns | 0,3182 |
| Sensory Cortex vs. Olfactory regions | 26,19 | 1,397 | 24,79 | 6,117 | 4 | 4 | 5,732 | 36 | Yes | * | 0,0118 |
| Sensory Cortex vs. Hippocampus | 26,19 | 0,01064 | 26,18 | 6,117 | 4 | 4 | 6,053 | 36 | Yes | ** | 0,0064 |
| Sensory Cortex vs. Thalamus | 26,19 | 26,42 | -0,2281 | 6,117 | 4 | 4 | 0,05274 | 36 | No | ns | > 0,9999 |
| Sensory Cortex vs. Amygdala | 26,19 | 1,126 | 25,06 | 6,117 | 4 | 4 | 5,795 | 36 | Yes | * | 0,0105 |
| Sensory Cortex vs. Hypothalamus | 26,19 | 6,8 | 19,39 | 6,117 | 4 | 4 | 4,483 | 36 | No | ns | 0,1047 |
| Sensory Cortex vs. Brainstem | 26,19 | 12,27 | 13,92 | 6,117 | 4 | 4 | 3,219 | 36 | No | ns | 0,51 |
| Sensory Cortex vs. Striatum | 26,19 | 5,311 | 20,88 | 6,117 | 4 | 4 | 4,827 | 36 | No | ns | 0,0601 |
| Sensory Cortex vs. Claustrum | 26,19 | 2,69 | 23,5 | 6,117 | 4 | 4 | 5,433 | 36 | Yes | * | 0,0207 |
| Motor cortex vs. Association cortex | 1,423 | 6,064 | -4,641 | 6,117 | 4 | 4 | 1,073 | 36 | No | ns | 0,9997 |
| Motor cortex vs. Prefrontal cortex | 1,423 | 10,3 | -8,881 | 6,117 | 4 | 4 | 2,053 | 36 | No | ns | 0,944 |
| Motor cortex vs. Olfactory regions | 1,423 | 1,397 | 0,02622 | 6,117 | 4 | 4 | 0,006063 | 36 | No | ns | > 0,9999 |
| Motor cortex vs. Hippocampus | 1,423 | 0,01064 | 1,412 | 6,117 | 4 | 4 | 0,3266 | 36 | No | ns | > 0,9999 |
| Motor cortex vs. Thalamus | 1,423 | 26,42 | -24,99 | 6,117 | 4 | 4 | 5,779 | 36 | Yes | * | 0,0108 |
| Motor cortex vs. Amygdala | 1,423 | 1,126 | 0,2975 | 6,117 | 4 | 4 | 0,06878 | 36 | No | ns | > 0,9999 |
| Motor cortex vs. Hypothalamus | 1,423 | 6,8 | -5,377 | 6,117 | 4 | 4 | 1,243 | 36 | No | ns | 0,999 |
| Motor cortex vs. Brainstem | 1,423 | 12,27 | -10,84 | 6,117 | 4 | 4 | 2,507 | 36 | No | ns | 0,8215 |
| Motor cortex vs. Striatum | 1,423 | 5,311 | -3,888 | 6,117 | 4 | 4 | 0,899 | 36 | No | ns | > 0,9999 |
| Motor cortex vs. Claustrum | 1,423 | 2,69 | -1,267 | 6,117 | 4 | 4 | 0,2929 | 36 | No | ns | > 0,9999 |
| Association cortex vs. Prefrontal cortex | 6,064 | 10,3 | -4,24 | 6,117 | 4 | 4 | 0,9804 | 36 | No | ns | 0,9999 |
| Association cortex vs. Olfactory regions | 6,064 | 1,397 | 4,667 | 6,117 | 4 | 4 | 1,079 | 36 | No | ns | 0,9997 |
| Association cortex vs. Hippocampus | 6,064 | 0,01064 | 6,053 | 6,117 | 4 | 4 | 1,4 | 36 | No | ns | 0,997 |
| Association cortex vs. Thalamus | 6,064 | 26,42 | -20,35 | 6,117 | 4 | 4 | 4,706 | 36 | No | ns | 0,0734 |
| Association cortex vs. Amygdala | 6,064 | 1,126 | 4,938 | 6,117 | 4 | 4 | 1,142 | 36 | No | ns | 0,9995 |
| Association cortex vs. Hypothalamus | 6,064 | 6,8 | -0,7359 | 6,117 | 4 | 4 | 0,1702 | 36 | No | ns | > 0,9999 |
| Association cortex vs. Brainstem | 6,064 | 12,27 | -6,203 | 6,117 | 4 | 4 | 1,434 | 36 | No | ns | 0,9963 |
| Association cortex vs. Striatum | 6,064 | 5,311 | 0,7527 | 6,117 | 4 | 4 | 0,174 | 36 | No | ns | > 0,9999 |
| Association cortex vs. Claustrum | 6,064 | 2,69 | 3,374 | 6,117 | 4 | 4 | 0,7801 | 36 | No | ns | > 0,9999 |
| Prefrontal cortex vs. Olfactory regions | 10,3 | 1,397 | 8,908 | 6,117 | 4 | 4 | 2,059 | 36 | No | ns | 0,9429 |
| Prefrontal cortex vs. Hippocampus | 10,3 | 0,01064 | 10,29 | 6,117 | 4 | 4 | 2,38 | 36 | No | ns | 0,8642 |
| Prefrontal cortex vs. Thalamus | 10,3 | 26,42 | -16,11 | 6,117 | 4 | 4 | 3,725 | 36 | No | ns | 0,2989 |
| Prefrontal cortex vs. Amygdala | 10,3 | 1,126 | 9,179 | 6,117 | 4 | 4 | 2,122 | 36 | No | ns | 0,9308 |
| Prefrontal cortex vs. Hypothalamus | 10,3 | 6,8 | 3,504 | 6,117 | 4 | 4 | 0,8102 | 36 | No | ns | > 0,9999 |
| Prefrontal cortex vs. Brainstem | 10,3 | 12,27 | -1,963 | 6,117 | 4 | 4 | 0,4537 | 36 | No | ns | > 0,9999 |
| Prefrontal cortex vs. Striatum | 10,3 | 5,311 | 4,993 | 6,117 | 4 | 4 | 1,154 | 36 | No | ns | 0,9995 |
| Prefrontal cortex vs. Claustrum | 10,3 | 2,69 | 7,615 | 6,117 | 4 | 4 | 1,761 | 36 | No | ns | 0,981 |
| Olfactory regions vs. Hippocampus | 1,397 | 0,01064 | 1,386 | 6,117 | 4 | 4 | 0,3205 | 36 | No | ns | > 0,9999 |
| Olfactory regions vs. Thalamus | 1,397 | 26,42 | -25,02 | 6,117 | 4 | 4 | 5,785 | 36 | Yes | * | 0,0107 |
| Olfactory regions vs. Amygdala | 1,397 | 1,126 | 0,2713 | 6,117 | 4 | 4 | 0,06272 | 36 | No | ns | > 0,9999 |
| Olfactory regions vs. Hypothalamus | 1,397 | 6,8 | -5,403 | 6,117 | 4 | 4 | 1,249 | 36 | No | ns | 0,9989 |
| Olfactory regions vs. Brainstem | 1,397 | 12,27 | -10,87 | 6,117 | 4 | 4 | 2,513 | 36 | No | ns | 0,8193 |
| Olfactory regions vs. Striatum | 1,397 | 5,311 | -3,915 | 6,117 | 4 | 4 | 0,905 | 36 | No | ns | > 0,9999 |
| Olfactory regions vs. Claustrum | 1,397 | 2,69 | -1,293 | 6,117 | 4 | 4 | 0,2989 | 36 | No | ns | > 0,9999 |
| Hippocampus vs. Thalamus | 0,01064 | 26,42 | -26,41 | 6,117 | 4 | 4 | 6,105 | 36 | Yes | ** | 0,0057 |
| Hippocampus vs. Amygdala | 0,01064 | 1,126 | -1,115 | 6,117 | 4 | 4 | 0,2578 | 36 | No | ns | > 0,9999 |
| Hippocampus vs. Hypothalamus | 0,01064 | 6,8 | -6,789 | 6,117 | 4 | 4 | 1,57 | 36 | No | ns | 0,9922 |
| Hippocampus vs. Brainstem | 0,01064 | 12,27 | -12,26 | 6,117 | 4 | 4 | 2,834 | 36 | No | ns | 0,688 |
| Hippocampus vs. Striatum | 0,01064 | 5,311 | -5,301 | 6,117 | 4 | 4 | 1,226 | 36 | No | ns | 0,9991 |
| Hippocampus vs. Claustrum | 0,01064 | 2,69 | -2,679 | 6,117 | 4 | 4 | 0,6194 | 36 | No | ns | > 0,9999 |
| Thalamus vs. Amygdala | 26,42 | 1,126 | 25,29 | 6,117 | 4 | 4 | 5,848 | 36 | Yes | ** | 0,0095 |
| Thalamus vs. Hypothalamus | 26,42 | 6,8 | 19,62 | 6,117 | 4 | 4 | 4,536 | 36 | No | ns | 0,0964 |
| Thalamus vs. Brainstem | 26,42 | 12,27 | 14,15 | 6,117 | 4 | 4 | 3,272 | 36 | No | ns | 0,4859 |
| Thalamus vs. Striatum | 26,42 | 5,311 | 21,11 | 6,117 | 4 | 4 | 4,88 | 36 | No | ns | 0,055 |
| Thalamus vs. Claustrum | 26,42 | 2,69 | 23,73 | 6,117 | 4 | 4 | 5,486 | 36 | Yes | * | 0,0188 |
| Amygdala vs. Hypothalamus | 1,126 | 6,8 | -5,674 | 6,117 | 4 | 4 | 1,312 | 36 | No | ns | 0,9983 |
| Amygdala vs. Brainstem | 1,126 | 12,27 | -11,14 | 6,117 | 4 | 4 | 2,576 | 36 | No | ns | 0,796 |
| Amygdala vs. Striatum | 1,126 | 5,311 | -4,186 | 6,117 | 4 | 4 | 0,9678 | 36 | No | ns | > 0,9999 |
| Amygdala vs. Claustrum | 1,126 | 2,69 | -1,564 | 6,117 | 4 | 4 | 0,3617 | 36 | No | ns | > 0,9999 |
| Hypothalamus vs. Brainstem | 6,8 | 12,27 | -5,467 | 6,117 | 4 | 4 | 1,264 | 36 | No | ns | 0,9988 |
| Hypothalamus vs. Striatum | 6,8 | 5,311 | 1,489 | 6,117 | 4 | 4 | 0,3442 | 36 | No | ns | > 0,9999 |
| Hypothalamus vs. Claustrum | 6,8 | 2,69 | 4,11 | 6,117 | 4 | 4 | 0,9503 | 36 | No | ns | > 0,9999 |
| Brainstem vs. Striatum | 12,27 | 5,311 | 6,956 | 6,117 | 4 | 4 | 1,608 | 36 | No | ns | 0,9905 |
| Brainstem vs. Claustrum | 12,27 | 2,69 | 9,577 | 6,117 | 4 | 4 | 2,214 | 36 | No | ns | 0,9102 |
| Striatum vs. Claustrum | 5,311 | 2,69 | 2,622 | 6,117 | 4 | 4 | 0,6061 | 36 | No | ns | > 0,9999 |

**Table Analyzed** **LO outputs**

| **ANOVA summary** | |
|---|---|
| F | 8,524 |
| P value | < 0,0001 |
| P value summary | **** |
| Are differences among means statistically significant? (P < ( | Yes |
| R square | 0,4941 |

**Tukey's multiple comparisons test**

| | Mean 1 | Mean 2 | Mean Diff, | SE of diff, | n1 | n2 | q | DF | Significant | Summary | Adjusted P |
|---|---|---|---|---|---|---|---|---|---|---|---|
| Sensory Cortex vs. Motor cortex | 3,653 | 4,2 | -0,547 | 4,622 | 9 | 9 | 0,1674 | 96 | No | ns | > 0,9999 |
| Sensory Cortex vs. Association cortex | 3,653 | 2,284 | 1,369 | 4,622 | 9 | 9 | 0,419 | 96 | No | ns | > 0,9999 |
| Sensory Cortex vs. Prefrontal cortex | 3,653 | 16 | -12,35 | 4,622 | 9 | 9 | 3,778 | 96 | No | ns | 0,2565 |
| Sensory Cortex vs. Olfactory regions | 3,653 | 2,74 | 0,9131 | 4,622 | 9 | 9 | 0,2794 | 96 | No | ns | > 0,9999 |
| Sensory Cortex vs. Hippocampus | 3,653 | 0,05969 | 3,593 | 4,622 | 9 | 9 | 1,1 | 96 | No | ns | 0,9997 |
| Sensory Cortex vs. Thalamus | 3,653 | 33,08 | -29,43 | 4,622 | 9 | 9 | 9,004 | 96 | Yes | **** | < 0,0001 |
| Sensory Cortex vs. Amygdala | 3,653 | 2,236 | 1,418 | 4,622 | 9 | 9 | 0,4338 | 96 | No | ns | > 0,9999 |
| Sensory Cortex vs. Hypothalamus | 3,653 | 6,252 | -2,599 | 4,622 | 9 | 9 | 0,7952 | 96 | No | ns | > 0,9999 |
| Sensory Cortex vs. Brainstem | 3,653 | 17,84 | -14,19 | 4,622 | 9 | 9 | 4,341 | 96 | No | ns | 0,1052 |
| Sensory Cortex vs. Striatum | 3,653 | 8,122 | -4,469 | 4,622 | 9 | 9 | 1,367 | 96 | No | ns | 0,9981 |
| Sensory Cortex vs. Claustrum | 3,653 | 3,533 | 0,1201 | 4,622 | 9 | 9 | 0,03674 | 96 | No | ns | > 0,9999 |
| Motor cortex vs. Association cortex | 4,2 | 2,284 | 1,916 | 4,622 | 9 | 9 | 0,5864 | 96 | No | ns | > 0,9999 |
| Motor cortex vs. Prefrontal cortex | 4,2 | 16 | -11,8 | 4,622 | 9 | 9 | 3,611 | 96 | No | ns | 0,321 |
| Motor cortex vs. Olfactory regions | 4,2 | 2,74 | 1,46 | 4,622 | 9 | 9 | 0,4467 | 96 | No | ns | > 0,9999 |
| Motor cortex vs. Hippocampus | 4,2 | 0,05969 | 4,14 | 4,622 | 9 | 9 | 1,267 | 96 | No | ns | 0,999 |
| Motor cortex vs. Thalamus | 4,2 | 33,08 | -28,88 | 4,622 | 9 | 9 | 8,836 | 96 | Yes | **** | < 0,0001 |
| Motor cortex vs. Amygdala | 4,2 | 2,236 | 1,965 | 4,622 | 9 | 9 | 0,6011 | 96 | No | ns | > 0,9999 |
| Motor cortex vs. Hypothalamus | 4,2 | 6,252 | -2,052 | 4,622 | 9 | 9 | 0,6278 | 96 | No | ns | > 0,9999 |
| Motor cortex vs. Brainstem | 4,2 | 17,84 | -13,64 | 4,622 | 9 | 9 | 4,173 | 96 | No | ns | 0,1401 |
| Motor cortex vs. Striatum | 4,2 | 8,122 | -3,922 | 4,622 | 9 | 9 | 1,2 | 96 | No | ns | 0,9994 |
| Motor cortex vs. Claustrum | 4,2 | 3,533 | 0,6671 | 4,622 | 9 | 9 | 0,2041 | 96 | No | ns | > 0,9999 |
| Association cortex vs. Prefrontal cortex | 2,284 | 16 | -13,72 | 4,622 | 9 | 9 | 4,197 | 96 | No | ns | 0,1346 |
| Association cortex vs. Olfactory regions | 2,284 | 2,74 | -0,4563 | 4,622 | 9 | 9 | 0,1396 | 96 | No | ns | > 0,9999 |
| Association cortex vs. Hippocampus | 2,284 | 0,05969 | 2,224 | 4,622 | 9 | 9 | 0,6805 | 96 | No | ns | > 0,9999 |
| Association cortex vs. Thalamus | 2,284 | 33,08 | -30,8 | 4,622 | 9 | 9 | 9,423 | 96 | Yes | **** | < 0,0001 |
| Association cortex vs. Amygdala | 2,284 | 2,236 | 0,04824 | 4,622 | 9 | 9 | 0,01476 | 96 | No | ns | > 0,9999 |
| Association cortex vs. Hypothalamus | 2,284 | 6,252 | -3,968 | 4,622 | 9 | 9 | 1,214 | 96 | No | ns | 0,9993 |
| Association cortex vs. Brainstem | 2,284 | 17,84 | -15,56 | 4,622 | 9 | 9 | 4,76 | 96 | Yes | * | 0,0479 |
| Association cortex vs. Striatum | 2,284 | 8,122 | -5,838 | 4,622 | 9 | 9 | 1,786 | 96 | No | ns | 0,9817 |
| Association cortex vs. Claustrum | 2,284 | 3,533 | -1,249 | 4,622 | 9 | 9 | 0,3823 | 96 | No | ns | > 0,9999 |
| Prefrontal cortex vs. Olfactory regions | 16 | 2,74 | 13,26 | 4,622 | 9 | 9 | 4,058 | 96 | No | ns | 0,169 |
| Prefrontal cortex vs. Hippocampus | 16 | 0,05969 | 15,94 | 4,622 | 9 | 9 | 4,878 | 96 | Yes | * | 0,0377 |
| Prefrontal cortex vs. Thalamus | 16 | 33,08 | -17,08 | 4,622 | 9 | 9 | 5,225 | 96 | Yes | * | 0,0179 |
| Prefrontal cortex vs. Amygdala | 16 | 2,236 | 13,77 | 4,622 | 9 | 9 | 4,212 | 96 | No | ns | 0,1313 |
| Prefrontal cortex vs. Hypothalamus | 16 | 6,252 | 9,75 | 4,622 | 9 | 9 | 2,983 | 96 | No | ns | 0,6174 |
| Prefrontal cortex vs. Brainstem | 16 | 17,84 | -1,838 | 4,622 | 9 | 9 | 0,5625 | 96 | No | ns | > 0,9999 |
| Prefrontal cortex vs. Striatum | 16 | 8,122 | 7,88 | 4,622 | 9 | 9 | 2,411 | 96 | No | ns | 0,8616 |
| Prefrontal cortex vs. Claustrum | 16 | 3,533 | 12,47 | 4,622 | 9 | 9 | 3,815 | 96 | No | ns | 0,2436 |
| Olfactory regions vs. Hippocampus | 2,74 | 0,05969 | 2,68 | 4,622 | 9 | 9 | 0,8201 | 96 | No | ns | > 0,9999 |
| Olfactory regions vs. Thalamus | 2,74 | 33,08 | -30,34 | 4,622 | 9 | 9 | 9,283 | 96 | Yes | **** | < 0,0001 |
| Olfactory regions vs. Amygdala | 2,74 | 2,236 | 0,5046 | 4,622 | 9 | 9 | 0,1544 | 96 | No | ns | > 0,9999 |
| Olfactory regions vs. Hypothalamus | 2,74 | 6,252 | -3,512 | 4,622 | 9 | 9 | 1,075 | 96 | No | ns | 0,9998 |
| Olfactory regions vs. Brainstem | 2,74 | 17,84 | -15,1 | 4,622 | 9 | 9 | 4,62 | 96 | No | ns | 0,0629 |
| Olfactory regions vs. Striatum | 2,74 | 8,122 | -5,382 | 4,622 | 9 | 9 | 1,647 | 96 | No | ns | 0,9904 |
| Olfactory regions vs. Claustrum | 2,74 | 3,533 | -0,793 | 4,622 | 9 | 9 | 0,2426 | 96 | No | ns | > 0,9999 |
| Hippocampus vs. Thalamus | 0,05969 | 33,08 | -33,02 | 4,622 | 9 | 9 | 10,1 | 96 | Yes | **** | < 0,0001 |
| Hippocampus vs. Amygdala | 0,05969 | 2,236 | -2,176 | 4,622 | 9 | 9 | 0,6658 | 96 | No | ns | > 0,9999 |
| Hippocampus vs. Hypothalamus | 0,05969 | 6,252 | -6,192 | 4,622 | 9 | 9 | 1,895 | 96 | No | ns | 0,9715 |
| Hippocampus vs. Brainstem | 0,05969 | 17,84 | -17,78 | 4,622 | 9 | 9 | 5,44 | 96 | Yes | * | 0,011 |
| Hippocampus vs. Striatum | 0,05969 | 8,122 | -8,062 | 4,622 | 9 | 9 | 2,467 | 96 | No | ns | 0,8428 |
| Hippocampus vs. Claustrum | 0,05969 | 3,533 | -3,473 | 4,622 | 9 | 9 | 1,063 | 96 | No | ns | 0,9998 |
| Thalamus vs. Amygdala | 33,08 | 2,236 | 30,84 | 4,622 | 9 | 9 | 9,438 | 96 | Yes | **** | < 0,0001 |
| Thalamus vs. Hypothalamus | 33,08 | 6,252 | 26,83 | 4,622 | 9 | 9 | 8,209 | 96 | Yes | **** | < 0,0001 |
| Thalamus vs. Brainstem | 33,08 | 17,84 | 15,24 | 4,622 | 9 | 9 | 4,663 | 96 | No | ns | 0,0579 |
| Thalamus vs. Striatum | 33,08 | 8,122 | 24,96 | 4,622 | 9 | 9 | 7,636 | 96 | Yes | **** | < 0,0001 |
| Thalamus vs. Claustrum | 33,08 | 3,533 | 29,55 | 4,622 | 9 | 9 | 9,041 | 96 | Yes | **** | < 0,0001 |
| Amygdala vs. Hypothalamus | 2,236 | 6,252 | -4,016 | 4,622 | 9 | 9 | 1,229 | 96 | No | ns | 0,9993 |
| Amygdala vs. Brainstem | 2,236 | 17,84 | -15,6 | 4,622 | 9 | 9 | 4,775 | 96 | Yes | * | 0,0465 |
| Amygdala vs. Striatum | 2,236 | 8,122 | -5,886 | 4,622 | 9 | 9 | 1,801 | 96 | No | ns | 0,9805 |
| Amygdala vs. Claustrum | 2,236 | 3,533 | -1,298 | 4,622 | 9 | 9 | 0,397 | 96 | No | ns | > 0,9999 |
| Hypothalamus vs. Brainstem | 6,252 | 17,84 | -11,59 | 4,622 | 9 | 9 | 3,546 | 96 | No | ns | 0,3484 |
| Hypothalamus vs. Striatum | 6,252 | 8,122 | -1,87 | 4,622 | 9 | 9 | 0,5722 | 96 | No | ns | > 0,9999 |
| Hypothalamus vs. Claustrum | 6,252 | 3,533 | 2,719 | 4,622 | 9 | 9 | 0,8319 | 96 | No | ns | > 0,9999 |
| Brainstem vs. Striatum | 17,84 | 8,122 | 9,718 | 4,622 | 9 | 9 | 2,974 | 96 | No | ns | 0,6222 |
| Brainstem vs. Claustrum | 17,84 | 3,533 | 14,31 | 4,622 | 9 | 9 | 4,378 | 96 | No | ns | 0,0986 |
| Striatum vs. Claustrum | 8,122 | 3,533 | 4,589 | 4,622 | 9 | 9 | 1,404 | 96 | No | ns | 0,9975 |

**Table Analyzed** | **AI output**

**ANOVA summary**

| | |
|---|---|
| F | 2,451 |
| P value | 0,0134 |
| P value summary | * |
| Are differences among means statistically significant? (P < ( | Yes |
| R square | 0,31 |

**Tukey's multiple comparisons test**

| | Mean 1 | Mean 2 | Mean Diff, | SE of diff, | n1 | n2 | q | DF | Significant | Summary | Adjusted P |
|---|---|---|---|---|---|---|---|---|---|---|---|
| Sensory Cortex vs. Motor cortex | 10,21 | 2,834 | 7,377 | 4,392 | 6 | 6 | 2,376 | 60 | No | ns | 0,8701 |
| Sensory Cortex vs. Association cortex | 10,21 | 9,014 | 1,197 | 4,392 | 6 | 6 | 0,3853 | 60 | No | ns | > 0,9999 |
| Sensory Cortex vs. Prefrontal cortex | 10,21 | 13,51 | -3,296 | 4,392 | 6 | 6 | 1,061 | 60 | No | ns | 0,9998 |
| Sensory Cortex vs. Olfactory regions | 10,21 | 3,959 | 6,252 | 4,392 | 6 | 6 | 2,013 | 60 | No | ns | 0,9541 |
| Sensory Cortex vs. Hippocampus | 10,21 | 0,03486 | 10,18 | 4,392 | 6 | 6 | 3,277 | 60 | No | ns | 0,4764 |
| Sensory Cortex vs. Thalamus | 10,21 | 14,85 | -4,641 | 4,392 | 6 | 6 | 1,495 | 60 | No | ns | 0,9954 |
| Sensory Cortex vs. Amygdala | 10,21 | 12,68 | -2,468 | 4,392 | 6 | 6 | 0,7949 | 60 | No | ns | > 0,9999 |
| Sensory Cortex vs. Hypothalamus | 10,21 | 5,217 | 4,994 | 4,392 | 6 | 6 | 1,608 | 60 | No | ns | 0,9915 |
| Sensory Cortex vs. Brainstem | 10,21 | 5,431 | 4,78 | 4,392 | 6 | 6 | 1,539 | 60 | No | ns | 0,9941 |
| Sensory Cortex vs. Striatum | 10,21 | 13,94 | -3,728 | 4,392 | 6 | 6 | 1,201 | 60 | No | ns | 0,9994 |
| Sensory Cortex vs. Claustrum | 10,21 | 8,322 | 1,889 | 4,392 | 6 | 6 | 0,6082 | 60 | No | ns | > 0,9999 |
| Motor cortex vs. Association cortex | 2,834 | 9,014 | -6,181 | 4,392 | 6 | 6 | 1,99 | 60 | No | ns | 0,9575 |
| Motor cortex vs. Prefrontal cortex | 2,834 | 13,51 | -10,67 | 4,392 | 6 | 6 | 3,437 | 60 | No | ns | 0,4027 |
| Motor cortex vs. Olfactory regions | 2,834 | 3,959 | -1,126 | 4,392 | 6 | 6 | 0,3625 | 60 | No | ns | > 0,9999 |
| Motor cortex vs. Hippocampus | 2,834 | 0,03486 | 2,799 | 4,392 | 6 | 6 | 0,9013 | 60 | No | ns | > 0,9999 |
| Motor cortex vs. Thalamus | 2,834 | 14,85 | -12,02 | 4,392 | 6 | 6 | 3,87 | 60 | No | ns | 0,2343 |
| Motor cortex vs. Amygdala | 2,834 | 12,68 | -9,846 | 4,392 | 6 | 6 | 3,171 | 60 | No | ns | 0,5273 |
| Motor cortex vs. Hypothalamus | 2,834 | 5,217 | -2,383 | 4,392 | 6 | 6 | 0,7674 | 60 | No | ns | > 0,9999 |
| Motor cortex vs. Brainstem | 2,834 | 5,431 | -2,597 | 4,392 | 6 | 6 | 0,8364 | 60 | No | ns | > 0,9999 |
| Motor cortex vs. Striatum | 2,834 | 13,94 | -11,11 | 4,392 | 6 | 6 | 3,576 | 60 | No | ns | 0,3429 |
| Motor cortex vs. Claustrum | 2,834 | 8,322 | -5,489 | 4,392 | 6 | 6 | 1,768 | 60 | No | ns | 0,9822 |
| Association cortex vs. Prefrontal cortex | 9,014 | 13,51 | -4,493 | 4,392 | 6 | 6 | 1,447 | 60 | No | ns | 0,9965 |
| Association cortex vs. Olfactory regions | 9,014 | 3,959 | 5,055 | 4,392 | 6 | 6 | 1,628 | 60 | No | ns | 0,9907 |
| Association cortex vs. Hippocampus | 9,014 | 0,03486 | 8,98 | 4,392 | 6 | 6 | 2,892 | 60 | No | ns | 0,662 |
| Association cortex vs. Thalamus | 9,014 | 14,85 | -5,838 | 4,392 | 6 | 6 | 1,88 | 60 | No | ns | 0,9717 |
| Association cortex vs. Amygdala | 9,014 | 12,68 | -3,665 | 4,392 | 6 | 6 | 1,18 | 60 | No | ns | 0,9994 |
| Association cortex vs. Hypothalamus | 9,014 | 5,217 | 3,798 | 4,392 | 6 | 6 | 1,223 | 60 | No | ns | 0,9992 |
| Association cortex vs. Brainstem | 9,014 | 5,431 | 3,584 | 4,392 | 6 | 6 | 1,154 | 60 | No | ns | 0,9996 |
| Association cortex vs. Striatum | 9,014 | 13,94 | -4,925 | 4,392 | 6 | 6 | 1,586 | 60 | No | ns | 0,9925 |
| Association cortex vs. Claustrum | 9,014 | 8,322 | 0,6921 | 4,392 | 6 | 6 | 0,2229 | 60 | No | ns | > 0,9999 |
| Prefrontal cortex vs. Olfactory regions | 13,51 | 3,959 | 9,548 | 4,392 | 6 | 6 | 3,075 | 60 | No | ns | 0,5739 |
| Prefrontal cortex vs. Hippocampus | 13,51 | 0,03486 | 13,47 | 4,392 | 6 | 6 | 4,338 | 60 | No | ns | 0,1148 |
| Prefrontal cortex vs. Thalamus | 13,51 | 14,85 | -1,345 | 4,392 | 6 | 6 | 0,4331 | 60 | No | ns | > 0,9999 |
| Prefrontal cortex vs. Amygdala | 13,51 | 12,68 | 0,8276 | 4,392 | 6 | 6 | 0,2665 | 60 | No | ns | > 0,9999 |
| Prefrontal cortex vs. Hypothalamus | 13,51 | 5,217 | 8,29 | 4,392 | 6 | 6 | 2,67 | 60 | No | ns | 0,7617 |
| Prefrontal cortex vs. Brainstem | 13,51 | 5,431 | 8,076 | 4,392 | 6 | 6 | 2,601 | 60 | No | ns | 0,79 |
| Prefrontal cortex vs. Striatum | 13,51 | 13,94 | -0,4321 | 4,392 | 6 | 6 | 0,1392 | 60 | No | ns | > 0,9999 |
| Prefrontal cortex vs. Claustrum | 13,51 | 8,322 | 5,185 | 4,392 | 6 | 6 | 1,67 | 60 | No | ns | 0,9886 |
| Olfactory regions vs. Hippocampus | 3,959 | 0,03486 | 3,924 | 4,392 | 6 | 6 | 1,264 | 60 | No | ns | 0,999 |
| Olfactory regions vs. Thalamus | 3,959 | 14,85 | -10,89 | 4,392 | 6 | 6 | 3,508 | 60 | No | ns | 0,3718 |
| Olfactory regions vs. Amygdala | 3,959 | 12,68 | -8,72 | 4,392 | 6 | 6 | 2,808 | 60 | No | ns | 0,7008 |
| Olfactory regions vs. Hypothalamus | 3,959 | 5,217 | -1,257 | 4,392 | 6 | 6 | 0,4049 | 60 | No | ns | > 0,9999 |
| Olfactory regions vs. Brainstem | 3,959 | 5,431 | -1,471 | 4,392 | 6 | 6 | 0,4739 | 60 | No | ns | > 0,9999 |
| Olfactory regions vs. Striatum | 3,959 | 13,94 | -9,98 | 4,392 | 6 | 6 | 3,214 | 60 | No | ns | 0,5065 |
| Olfactory regions vs. Claustrum | 3,959 | 8,322 | -4,363 | 4,392 | 6 | 6 | 1,405 | 60 | No | ns | 0,9973 |
| Hippocampus vs. Thalamus | 0,03486 | 14,85 | -14,82 | 4,392 | 6 | 6 | 4,772 | 60 | No | ns | 0,0536 |
| Hippocampus vs. Amygdala | 0,03486 | 12,68 | -12,64 | 4,392 | 6 | 6 | 4,072 | 60 | No | ns | 0,175 |
| Hippocampus vs. Hypothalamus | 0,03486 | 5,217 | -5,182 | 4,392 | 6 | 6 | 1,669 | 60 | No | ns | 0,9886 |
| Hippocampus vs. Brainstem | 0,03486 | 5,431 | -5,396 | 4,392 | 6 | 6 | 1,738 | 60 | No | ns | 0,9844 |
| Hippocampus vs. Striatum | 0,03486 | 13,94 | -13,9 | 4,392 | 6 | 6 | 4,478 | 60 | No | ns | 0,0907 |
| Hippocampus vs. Claustrum | 0,03486 | 8,322 | -8,287 | 4,392 | 6 | 6 | 2,669 | 60 | No | ns | 0,7621 |
| Thalamus vs. Amygdala | 14,85 | 12,68 | 2,173 | 4,392 | 6 | 6 | 0,6997 | 60 | No | ns | > 0,9999 |
| Thalamus vs. Hypothalamus | 14,85 | 5,217 | 9,635 | 4,392 | 6 | 6 | 3,103 | 60 | No | ns | 0,5602 |
| Thalamus vs. Brainstem | 14,85 | 5,431 | 9,421 | 4,392 | 6 | 6 | 3,034 | 60 | No | ns | 0,5937 |
| Thalamus vs. Striatum | 14,85 | 13,94 | 0,9129 | 4,392 | 6 | 6 | 0,294 | 60 | No | ns | > 0,9999 |
| Thalamus vs. Claustrum | 14,85 | 8,322 | 6,53 | 4,392 | 6 | 6 | 2,103 | 60 | No | ns | 0,9385 |
| Amygdala vs. Hypothalamus | 12,68 | 5,217 | 7,463 | 4,392 | 6 | 6 | 2,403 | 60 | No | ns | 0,8614 |
| Amygdala vs. Brainstem | 12,68 | 5,431 | 7,248 | 4,392 | 6 | 6 | 2,334 | 60 | No | ns | 0,8826 |
| Amygdala vs. Striatum | 12,68 | 13,94 | -1,26 | 4,392 | 6 | 6 | 0,4057 | 60 | No | ns | > 0,9999 |
| Amygdala vs. Claustrum | 12,68 | 8,322 | 4,357 | 4,392 | 6 | 6 | 1,403 | 60 | No | ns | 0,9973 |
| Hypothalamus vs. Brainstem | 5,217 | 5,431 | -0,2142 | 4,392 | 6 | 6 | 0,06897 | 60 | No | ns | > 0,9999 |
| Hypothalamus vs. Striatum | 5,217 | 13,94 | -8,722 | 4,392 | 6 | 6 | 2,809 | 60 | No | ns | 0,7005 |
| Hypothalamus vs. Claustrum | 5,217 | 8,322 | -3,106 | 4,392 | 6 | 6 | 1 | 60 | No | ns | 0,9999 |
| Brainstem vs. Striatum | 5,431 | 13,94 | -8,508 | 4,392 | 6 | 6 | 2,74 | 60 | No | ns | 0,7315 |
| Brainstem vs. Claustrum | 5,431 | 8,322 | -2,891 | 4,392 | 6 | 6 | 0,9311 | 60 | No | ns | > 0,9999 |
| Striatum vs. Claustrum | 13,94 | 8,322 | 5,617 | 4,392 | 6 | 6 | 1,809 | 60 | No | ns | 0,9788 |

**Table Analyzed** | **DLO output**

**ANOVA summary**

| | |
|---|---|
| F | 11,56 |
| P value | < 0,0001 |
| P value summary | **** |
| Are differences among means statistically significant? (P < ( | Yes |
| R square | 0,9138 |

**Tukey's multiple comparisons test**

| | Mean 1 | Mean 2 | Mean Diff, | SE of diff, | n1 | n2 | q | DF | Significant | Summary | Adjusted P |
|---|---|---|---|---|---|---|---|---|---|---|---|
| Sensory Cortex vs. Motor cortex | 7,966 | 7,727 | 0,2391 | 3,609 | 2 | 2 | 0,09369 | 12 | No | ns | > 0,9999 |
| Sensory Cortex vs. Association cortex | 7,966 | 3,2 | 4,766 | 3,609 | 2 | 2 | 1,868 | 12 | No | ns | 0,9598 |
| Sensory Cortex vs. Prefrontal cortex | 7,966 | 25,7 | -17,73 | 3,609 | 2 | 2 | 6,948 | 12 | Yes | * | 0,0113 |
| Sensory Cortex vs. Olfactory regions | 7,966 | 1,058 | 6,908 | 3,609 | 2 | 2 | 2,707 | 12 | No | ns | 0,7341 |
| Sensory Cortex vs. Hippocampus | 7,966 | 0,005417 | 7,961 | 3,609 | 2 | 2 | 3,119 | 12 | No | ns | 0,5728 |
| Sensory Cortex vs. Thalamus | 7,966 | 24,54 | -16,58 | 3,609 | 2 | 2 | 6,496 | 12 | Yes | * | 0,0187 |
| Sensory Cortex vs. Amygdala | 7,966 | 2,722 | 5,244 | 3,609 | 2 | 2 | 2,055 | 12 | No | ns | 0,9286 |
| Sensory Cortex vs. Hypothalamus | 7,966 | 2,39 | 5,576 | 3,609 | 2 | 2 | 2,185 | 12 | No | ns | 0,8999 |
| Sensory Cortex vs. Brainstem | 7,966 | 12,84 | -4,87 | 3,609 | 2 | 2 | 1,908 | 12 | No | ns | 0,954 |
| Sensory Cortex vs. Striatum | 7,966 | 8,778 | -0,8124 | 3,609 | 2 | 2 | 0,3183 | 12 | No | ns | > 0,9999 |
| Sensory Cortex vs. Claustrum | 7,966 | 3,075 | 4,891 | 3,609 | 2 | 2 | 1,916 | 12 | No | ns | 0,9528 |
| Motor cortex vs. Association cortex | 7,727 | 3,2 | 4,527 | 3,609 | 2 | 2 | 1,774 | 12 | No | ns | 0,9712 |
| Motor cortex vs. Prefrontal cortex | 7,727 | 25,7 | -17,97 | 3,609 | 2 | 2 | 7,042 | 12 | Yes | * | 0,0102 |
| Motor cortex vs. Olfactory regions | 7,727 | 1,058 | 6,669 | 3,609 | 2 | 2 | 2,613 | 12 | No | ns | 0,7686 |
| Motor cortex vs. Hippocampus | 7,727 | 0,005417 | 7,722 | 3,609 | 2 | 2 | 3,026 | 12 | No | ns | 0,6098 |
| Motor cortex vs. Thalamus | 7,727 | 24,54 | -16,82 | 3,609 | 2 | 2 | 6,589 | 12 | Yes | * | 0,0168 |
| Motor cortex vs. Amygdala | 7,727 | 2,722 | 5,005 | 3,609 | 2 | 2 | 1,961 | 12 | No | ns | 0,9457 |
| Motor cortex vs. Hypothalamus | 7,727 | 2,39 | 5,337 | 3,609 | 2 | 2 | 2,091 | 12 | No | ns | 0,9212 |
| Motor cortex vs. Brainstem | 7,727 | 12,84 | -5,109 | 3,609 | 2 | 2 | 2,002 | 12 | No | ns | 0,9386 |
| Motor cortex vs. Striatum | 7,727 | 8,778 | -1,051 | 3,609 | 2 | 2 | 0,412 | 12 | No | ns | > 0,9999 |
| Motor cortex vs. Claustrum | 7,727 | 3,075 | 4,652 | 3,609 | 2 | 2 | 1,823 | 12 | No | ns | 0,9656 |
| Association cortex vs. Prefrontal cortex | 3,2 | 25,7 | -22,5 | 3,609 | 2 | 2 | 8,816 | 12 | Yes | ** | 0,0016 |
| Association cortex vs. Olfactory regions | 3,2 | 1,058 | 2,142 | 3,609 | 2 | 2 | 0,8393 | 12 | No | ns | > 0,9999 |
| Association cortex vs. Hippocampus | 3,2 | 0,005417 | 3,194 | 3,609 | 2 | 2 | 1,252 | 12 | No | ns | 0,9979 |
| Association cortex vs. Thalamus | 3,2 | 24,54 | -21,34 | 3,609 | 2 | 2 | 8,363 | 12 | Yes | ** | 0,0025 |
| Association cortex vs. Amygdala | 3,2 | 2,722 | 0,4777 | 3,609 | 2 | 2 | 0,1872 | 12 | No | ns | > 0,9999 |
| Association cortex vs. Hypothalamus | 3,2 | 2,39 | 0,8096 | 3,609 | 2 | 2 | 0,3172 | 12 | No | ns | > 0,9999 |
| Association cortex vs. Brainstem | 3,2 | 12,84 | -9,636 | 3,609 | 2 | 2 | 3,776 | 12 | No | ns | 0,3389 |
| Association cortex vs. Striatum | 3,2 | 8,778 | -5,579 | 3,609 | 2 | 2 | 2,186 | 12 | No | ns | 0,8997 |
| Association cortex vs. Claustrum | 3,2 | 3,075 | 0,1245 | 3,609 | 2 | 2 | 0,04878 | 12 | No | ns | > 0,9999 |
| Prefrontal cortex vs. Olfactory regions | 25,7 | 1,058 | 24,64 | 3,609 | 2 | 2 | 9,655 | 12 | Yes | *** | 0,0007 |
| Prefrontal cortex vs. Hippocampus | 25,7 | 0,005417 | 25,69 | 3,609 | 2 | 2 | 10,07 | 12 | Yes | *** | 0,0005 |
| Prefrontal cortex vs. Thalamus | 25,7 | 24,54 | 1,155 | 3,609 | 2 | 2 | 0,4526 | 12 | No | ns | > 0,9999 |
| Prefrontal cortex vs. Amygdala | 25,7 | 2,722 | 22,98 | 3,609 | 2 | 2 | 9,003 | 12 | Yes | ** | 0,0013 |
| Prefrontal cortex vs. Hypothalamus | 25,7 | 2,39 | 23,31 | 3,609 | 2 | 2 | 9,133 | 12 | Yes | ** | 0,0011 |
| Prefrontal cortex vs. Brainstem | 25,7 | 12,84 | 12,86 | 3,609 | 2 | 2 | 5,04 | 12 | No | ns | 0,0943 |
| Prefrontal cortex vs. Striatum | 25,7 | 8,778 | 16,92 | 3,609 | 2 | 2 | 6,63 | 12 | Yes | * | 0,0161 |
| Prefrontal cortex vs. Claustrum | 25,7 | 3,075 | 22,62 | 3,609 | 2 | 2 | 8,865 | 12 | Yes | ** | 0,0015 |
| Olfactory regions vs. Hippocampus | 1,058 | 0,005417 | 1,052 | 3,609 | 2 | 2 | 0,4124 | 12 | No | ns | > 0,9999 |
| Olfactory regions vs. Thalamus | 1,058 | 24,54 | -23,49 | 3,609 | 2 | 2 | 9,203 | 12 | Yes | ** | 0,0011 |
| Olfactory regions vs. Amygdala | 1,058 | 2,722 | -1,664 | 3,609 | 2 | 2 | 0,6521 | 12 | No | ns | > 0,9999 |
| Olfactory regions vs. Hypothalamus | 1,058 | 2,39 | -1,332 | 3,609 | 2 | 2 | 0,5221 | 12 | No | ns | > 0,9999 |
| Olfactory regions vs. Brainstem | 1,058 | 12,84 | -11,78 | 3,609 | 2 | 2 | 4,615 | 12 | No | ns | 0,1486 |
| Olfactory regions vs. Striatum | 1,058 | 8,778 | -7,721 | 3,609 | 2 | 2 | 3,025 | 12 | No | ns | 0,61 |
| Olfactory regions vs. Claustrum | 1,058 | 3,075 | -2,017 | 3,609 | 2 | 2 | 0,7905 | 12 | No | ns | > 0,9999 |
| Hippocampus vs. Thalamus | 0,005417 | 24,54 | -24,54 | 3,609 | 2 | 2 | 9,615 | 12 | Yes | *** | 0,0007 |
| Hippocampus vs. Amygdala | 0,005417 | 2,722 | -2,717 | 3,609 | 2 | 2 | 1,065 | 12 | No | ns | 0,9995 |
| Hippocampus vs. Hypothalamus | 0,005417 | 2,39 | -2,385 | 3,609 | 2 | 2 | 0,9345 | 12 | No | ns | 0,9998 |
| Hippocampus vs. Brainstem | 0,005417 | 12,84 | -12,83 | 3,609 | 2 | 2 | 5,028 | 12 | No | ns | 0,0956 |
| Hippocampus vs. Striatum | 0,005417 | 8,778 | -8,773 | 3,609 | 2 | 2 | 3,438 | 12 | No | ns | 0,4518 |
| Hippocampus vs. Claustrum | 0,005417 | 3,075 | -3,07 | 3,609 | 2 | 2 | 1,203 | 12 | No | ns | 0,9985 |
| Thalamus vs. Amygdala | 24,54 | 2,722 | 21,82 | 3,609 | 2 | 2 | 8,551 | 12 | Yes | ** | 0,002 |
| Thalamus vs. Hypothalamus | 24,54 | 2,39 | 22,15 | 3,609 | 2 | 2 | 8,681 | 12 | Yes | ** | 0,0018 |
| Thalamus vs. Brainstem | 24,54 | 12,84 | 11,71 | 3,609 | 2 | 2 | 4,587 | 12 | No | ns | 0,1529 |
| Thalamus vs. Striatum | 24,54 | 8,778 | 15,76 | 3,609 | 2 | 2 | 6,177 | 12 | Yes | * | 0,0267 |
| Thalamus vs. Claustrum | 24,54 | 3,075 | 21,47 | 3,609 | 2 | 2 | 8,412 | 12 | Yes | ** | 0,0024 |
| Amygdala vs. Hypothalamus | 2,722 | 2,39 | 0,3318 | 3,609 | 2 | 2 | 0,13 | 12 | No | ns | > 0,9999 |
| Amygdala vs. Brainstem | 2,722 | 12,84 | -10,11 | 3,609 | 2 | 2 | 3,963 | 12 | No | ns | 0,2853 |
| Amygdala vs. Striatum | 2,722 | 8,778 | -6,056 | 3,609 | 2 | 2 | 2,373 | 12 | No | ns | 0,8486 |
| Amygdala vs. Claustrum | 2,722 | 3,075 | -0,3533 | 3,609 | 2 | 2 | 0,1384 | 12 | No | ns | > 0,9999 |
| Hypothalamus vs. Brainstem | 2,39 | 12,84 | -10,45 | 3,609 | 2 | 2 | 4,093 | 12 | No | ns | 0,252 |
| Hypothalamus vs. Striatum | 2,39 | 8,778 | -6,388 | 3,609 | 2 | 2 | 2,503 | 12 | No | ns | 0,807 |
| Hypothalamus vs. Claustrum | 2,39 | 3,075 | -0,6851 | 3,609 | 2 | 2 | 0,2684 | 12 | No | ns | > 0,9999 |
| Brainstem vs. Striatum | 12,84 | 8,778 | 4,057 | 3,609 | 2 | 2 | 1,59 | 12 | No | ns | 0,9865 |
| Brainstem vs. Claustrum | 12,84 | 3,075 | 9,761 | 3,609 | 2 | 2 | 3,825 | 12 | No | ns | 0,3243 |
| Striatum vs. Claustrum | 8,778 | 3,075 | 5,703 | 3,609 | 2 | 2 | 2,235 | 12 | No | ns | 0,8874 |
