## Supplementary 2 for "Anatomical input-output streams within mouse orbitofrontal cortex subdivisions"

**Table Analyzed** **visual input**

**ANOVA summary**

| | |
|---|---|
| F | 2,957 |
| P value | 0,0749 |
| P value summary | ns |
| Are differences among means : | No |
| R square | 0,5419 |

**Tukey's multiple comparisons test**

| | Mean 1 | Mean 2 | Mean Diff, | SE of diff, | n1 | n2 | q | DF | Significant | Summary | Adjusted P Value |
|---|---|---|---|---|---|---|---|---|---|---|---|
| VO vs. LO | 0,9016 | 0,0698 | 0,8318 | 0,3954 | 4 | 3 | 2,975 | 10 | No | ns | 0,2898 |
| VO vs. MO | 0,9016 | 1,071 | -0,1692 | 0,4484 | 4 | 2 | 0,5337 | 10 | No | ns | 0,995 |
| VO vs. AI | 0,9016 | 0,009445 | 0,8921 | 0,3954 | 4 | 3 | 3,19 | 10 | No | ns | 0,2356 |
| VO vs. DLO | 0,9016 | 0 | 0,9016 | 0,3954 | 4 | 3 | 3,224 | 10 | No | ns | 0,2279 |
| LO vs. MO | 0,0698 | 1,071 | -1,001 | 0,4726 | 3 | 2 | 2,995 | 10 | No | ns | 0,2843 |
| LO vs. AI | 0,0698 | 0,009445 | 0,06036 | 0,4227 | 3 | 3 | 0,2019 | 10 | No | ns | 0,9999 |
| LO vs. DLO | 0,0698 | 0 | 0,0698 | 0,4227 | 3 | 3 | 0,2335 | 10 | No | ns | 0,9998 |
| MO vs. AI | 1,071 | 0,009445 | 1,061 | 0,4726 | 2 | 3 | 3,176 | 10 | No | ns | 0,239 |
| MO vs. DLO | 1,071 | 0 | 1,071 | 0,4726 | 2 | 3 | 3,204 | 10 | No | ns | 0,2325 |
| AI vs. DLO | 0,009445 | 0 | 0,009445 | 0,4227 | 3 | 3 | 0,0316 | 10 | No | ns | > 0,9999 |

**Table Analyzed** **somatosensory input**

**ANOVA summary**

| | |
|---|---|
| F | 1,206 |
| P value | 0,3671 |
| P value summary | ns |
| Are differences among means : | No |
| R square | 0,3254 |

**Tukey's multiple comparisons test**

| | Mean 1 | Mean 2 | Mean Diff, | SE of diff, | n1 | n2 | q | DF | Significant | Summary | Adjusted P Value |
|---|---|---|---|---|---|---|---|---|---|---|---|
| VO vs. LO | 1,027 | 0,9381 | 0,08849 | 0,4787 | 4 | 3 | 0,2614 | 10 | No | ns | 0,9997 |
| VO vs. MO | 1,027 | 0,03076 | 0,9959 | 0,5428 | 4 | 2 | 2,595 | 10 | No | ns | 0,4068 |
| VO vs. AI | 1,027 | 0,3877 | 0,639 | 0,4787 | 4 | 3 | 1,888 | 10 | No | ns | 0,6779 |
| VO vs. DLO | 1,027 | 0,8974 | 0,1293 | 0,4787 | 4 | 3 | 0,3819 | 10 | No | ns | 0,9986 |
| LO vs. MO | 0,9381 | 0,03076 | 0,9074 | 0,5722 | 3 | 2 | 2,243 | 10 | No | ns | 0,5366 |
| LO vs. AI | 0,9381 | 0,3877 | 0,5505 | 0,5118 | 3 | 3 | 1,521 | 10 | No | ns | 0,8148 |
| LO vs. DLO | 0,9381 | 0,8974 | 0,04079 | 0,5118 | 3 | 3 | 0,1127 | 10 | No | ns | > 0,9999 |
| MO vs. AI | 0,03076 | 0,3877 | -0,3569 | 0,5722 | 2 | 3 | 0,8821 | 10 | No | ns | 0,9678 |
| MO vs. DLO | 0,03076 | 0,8974 | -0,8666 | 0,5722 | 2 | 3 | 2,142 | 10 | No | ns | 0,5764 |
| AI vs. DLO | 0,3877 | 0,8974 | -0,5097 | 0,5118 | 3 | 3 | 1,409 | 10 | No | ns | 0,8514 |

**Table Analyzed** **auditory input**

**ANOVA summary**

| | |
|---|---|
| F | 3,943 |
| P value | 0,0357 |
| P value summary | * |
| Are differences among means : | Yes |
| R square | 0,612 |

**Tukey's multiple comparisons test**

| | Mean 1 | Mean 2 | Mean Diff, | SE of diff, | n1 | n2 | q | DF | Significant | Summary | Adjusted P Value |
|---|---|---|---|---|---|---|---|---|---|---|---|
| VO vs. LO | 1,62 | 0,8136 | 0,806 | 0,6671 | 4 | 3 | 1,709 | 10 | No | ns | 0,7474 |
| VO vs. MO | 1,62 | 1,219 | 0,4005 | 0,7564 | 4 | 2 | 0,7489 | 10 | No | ns | 0,9821 |
| VO vs. AI | 1,62 | 2,797 | -1,177 | 0,6671 | 4 | 3 | 2,496 | 10 | No | ns | 0,4413 |
| VO vs. DLO | 1,62 | 0,1165 | 1,503 | 0,6671 | 4 | 3 | 3,186 | 10 | No | ns | 0,2365 |
| LO vs. MO | 0,8136 | 1,219 | -0,4054 | 0,7973 | 3 | 2 | 0,7191 | 10 | No | ns | 0,9846 |
| LO vs. AI | 0,8136 | 2,797 | -1,983 | 0,7131 | 3 | 3 | 3,933 | 10 | No | ns | 0,1095 |
| LO vs. DLO | 0,8136 | 0,1165 | 0,697 | 0,7131 | 3 | 3 | 1,382 | 10 | No | ns | 0,8594 |
| MO vs. AI | 1,219 | 2,797 | -1,578 | 0,7973 | 2 | 3 | 2,799 | 10 | No | ns | 0,3405 |
| MO vs. DLO | 1,219 | 0,1165 | 1,102 | 0,7973 | 2 | 3 | 1,955 | 10 | No | ns | 0,6509 |
| AI vs. DLO | 2,797 | 0,1165 | 2,68 | 0,7131 | 3 | 3 | 5,316 | 10 | Yes | * | 0,0242 |

**Table Analyzed** | **olfactory input**

**ANOVA summary**

| | |
|---|---|
| F | 10,38 |
| P value | 0,0014 |
| P value summary | ** |
| Are differences among means : | Yes |
| R square | 0,8059 |

**Tukey's multiple comparisons test**

| | Mean 1 | Mean 2 | Mean Diff, | SE of diff, | n1 | n2 | q | DF | Significant | Summary | Adjusted P Value |
|---|---|---|---|---|---|---|---|---|---|---|---|
| VO vs. LO | 0,8653 | 5,325 | -4,459 | 2,689 | 4 | 3 | 2,345 | 10 | No | ns | 0,4972 |
| VO vs. MO | 0,8653 | 2,938 | -2,072 | 3,049 | 4 | 2 | 0,9612 | 10 | No | ns | 0,9565 |
| VO vs. AI | 0,8653 | 17,17 | -16,3 | 2,689 | 4 | 3 | 8,573 | 10 | Yes | *** | 0,0009 |
| VO vs. DLO | 0,8653 | 9,053 | -8,188 | 2,689 | 4 | 3 | 4,306 | 10 | No | ns | 0,0732 |
| LO vs. MO | 5,325 | 2,938 | 2,387 | 3,214 | 3 | 2 | 1,05 | 10 | No | ns | 0,9412 |
| LO vs. AI | 5,325 | 17,17 | -11,84 | 2,875 | 3 | 3 | 5,825 | 10 | Yes | * | 0,0139 |
| LO vs. DLO | 5,325 | 9,053 | -3,728 | 2,875 | 3 | 3 | 1,834 | 10 | No | ns | 0,699 |
| MO vs. AI | 2,938 | 17,17 | -14,23 | 3,214 | 2 | 3 | 6,261 | 10 | Yes | ** | 0,0088 |
| MO vs. DLO | 2,938 | 9,053 | -6,115 | 3,214 | 2 | 3 | 2,691 | 10 | No | ns | 0,3747 |
| AI vs. DLO | 17,17 | 9,053 | 8,113 | 2,875 | 3 | 3 | 3,991 | 10 | No | ns | 0,1029 |

**Table Analyzed** | **gustatory input**

**ANOVA summary**

| | |
|---|---|
| F | 7,223 |
| P value | 0,0053 |
| P value summary | ** |
| Are differences among means : | Yes |
| R square | 0,7429 |

**Tukey's multiple comparisons test**

| | Mean 1 | Mean 2 | Mean Diff, | SE of diff, | n1 | n2 | q | DF | Significant | Summary | Adjusted P Value |
|---|---|---|---|---|---|---|---|---|---|---|---|
| VO vs. LO | 0,1899 | 1,313 | -1,124 | 1,895 | 4 | 3 | 0,8386 | 10 | No | ns | 0,9731 |
| VO vs. MO | 0,1899 | 0,004059 | 0,1859 | 2,148 | 4 | 2 | 0,1223 | 10 | No | ns | > 0,9999 |
| VO vs. AI | 0,1899 | 9,127 | -8,937 | 1,895 | 4 | 3 | 6,671 | 10 | Yes | ** | 0,0057 |
| VO vs. DLO | 0,1899 | 0,8076 | -0,6177 | 1,895 | 4 | 3 | 0,4611 | 10 | No | ns | 0,9971 |
| LO vs. MO | 1,313 | 0,004059 | 1,309 | 2,265 | 3 | 2 | 0,8177 | 10 | No | ns | 0,9754 |
| LO vs. AI | 1,313 | 9,127 | -7,814 | 2,026 | 3 | 3 | 5,456 | 10 | Yes | * | 0,0208 |
| LO vs. DLO | 1,313 | 0,8076 | 0,5059 | 2,026 | 3 | 3 | 0,3532 | 10 | No | ns | 0,999 |
| MO vs. AI | 0,004059 | 9,127 | -9,123 | 2,265 | 2 | 3 | 5,697 | 10 | Yes | * | 0,016 |
| MO vs. DLO | 0,004059 | 0,8076 | -0,8036 | 2,265 | 2 | 3 | 0,5018 | 10 | No | ns | 0,996 |
| AI vs. DLO | 9,127 | 0,8076 | 8,32 | 2,026 | 3 | 3 | 5,809 | 10 | Yes | * | 0,0142 |

**Table Analyzed** **visual output**

**ANOVA summary**

| | |
|---|---|
| F | 6,539 |
| P value | 0,0017 |
| P value summary | ** |
| Are differences among means s | Yes |
| R square | 0,5792 |

**Tukey's multiple comparisons test**

| | Mean 1 | Mean 2 | Mean Diff, | SE of diff, | n1 | n2 | q | DF | Significant | Summary | Adjusted P Value |
|---|---|---|---|---|---|---|---|---|---|---|---|
| VO vs. LO | 20,07 | 1,852 | 18,22 | 4,055 | 4 | 9 | 6,356 | 19 | Yes | ** | 0,002 |
| VO vs. MO | 20,07 | 7,015 | 13,06 | 5,154 | 4 | 3 | 3,584 | 19 | No | ns | 0,1244 |
| VO vs. AI | 20,07 | 0,2824 | 19,79 | 4,356 | 4 | 6 | 6,426 | 19 | Yes | ** | 0,0018 |
| VO vs. DLO | 20,07 | 0,1498 | 19,93 | 5,844 | 4 | 2 | 4,822 | 19 | Yes | * | 0,0217 |
| LO vs. MO | 1,852 | 7,015 | -5,163 | 4,498 | 9 | 3 | 1,623 | 19 | No | ns | 0,7796 |
| LO vs. AI | 1,852 | 0,2824 | 1,569 | 3,556 | 9 | 6 | 0,624 | 19 | No | ns | 0,9915 |
| LO vs. DLO | 1,852 | 0,1498 | 1,702 | 5,275 | 9 | 2 | 0,4563 | 19 | No | ns | 0,9974 |
| MO vs. AI | 7,015 | 0,2824 | 6,733 | 4,771 | 3 | 6 | 1,996 | 19 | No | ns | 0,6284 |
| MO vs. DLO | 7,015 | 0,1498 | 6,865 | 6,16 | 3 | 2 | 1,576 | 19 | No | ns | 0,797 |
| AI vs. DLO | 0,2824 | 0,1498 | 0,1326 | 5,51 | 6 | 2 | 0,03404 | 19 | No | ns | > 0,9999 |

**Table Analyzed** **somatosensory output**

**ANOVA summary**

| | |
|---|---|
| F | 1,304 |
| P value | 0,304 |
| P value summary | ns |
| Are differences among means s | No |
| R square | 0,2154 |

**Tukey's multiple comparisons test**

| | Mean 1 | Mean 2 | Mean Diff, | SE of diff, | n1 | n2 | q | DF | Significant | Summary | Adjusted P Value |
|---|---|---|---|---|---|---|---|---|---|---|---|
| VO vs. LO | 2,109 | 1,134 | 0,9747 | 2,013 | 4 | 9 | 0,6848 | 19 | No | ns | 0,9879 |
| VO vs. MO | 2,109 | 0,5843 | 1,524 | 2,558 | 4 | 3 | 0,8426 | 19 | No | ns | 0,974 |
| VO vs. AI | 2,109 | 2,883 | -0,7748 | 2,162 | 4 | 6 | 0,5067 | 19 | No | ns | 0,9962 |
| VO vs. DLO | 2,109 | 6,548 | -4,439 | 2,901 | 4 | 2 | 2,164 | 19 | No | ns | 0,5566 |
| LO vs. MO | 1,134 | 0,5843 | 0,5497 | 2,233 | 9 | 3 | 0,3481 | 19 | No | ns | 0,9991 |
| LO vs. AI | 1,134 | 2,883 | -1,749 | 1,765 | 9 | 6 | 1,401 | 19 | No | ns | 0,8561 |
| LO vs. DLO | 1,134 | 6,548 | -5,414 | 2,619 | 9 | 2 | 2,924 | 19 | No | ns | 0,2739 |
| MO vs. AI | 0,5843 | 2,883 | -2,299 | 2,369 | 3 | 6 | 1,373 | 19 | No | ns | 0,8649 |
| MO vs. DLO | 0,5843 | 6,548 | -5,964 | 3,058 | 3 | 2 | 2,758 | 19 | No | ns | 0,3264 |
| AI vs. DLO | 2,883 | 6,548 | -3,664 | 2,735 | 6 | 2 | 1,895 | 19 | No | ns | 0,671 |

**Table Analyzed** **auditory output**

**ANOVA summary**

| | |
|---|---|
| F | 9,744 |
| P value | 0,0002 |
| P value summary | *** |
| Are differences among means s | Yes |
| R square | 0,6723 |

**Tukey's multiple comparisons test**

| | Mean 1 | Mean 2 | Mean Diff, | SE of diff, | n1 | n2 | q | DF | Significant | Summary | Adjusted P Value |
|---|---|---|---|---|---|---|---|---|---|---|---|
| VO vs. LO | 5,155 | 0,4928 | 4,662 | 0,7606 | 4 | 9 | 8,668 | 19 | Yes | **** | < 0,0001 |
| VO vs. MO | 5,155 | 1,75 | 3,405 | 0,9667 | 4 | 3 | 4,981 | 19 | Yes | * | 0,0171 |
| VO vs. AI | 5,155 | 1,411 | 3,743 | 0,817 | 4 | 6 | 6,48 | 19 | Yes | ** | 0,0017 |
| VO vs. DLO | 5,155 | 0,9667 | 4,188 | 1,096 | 4 | 2 | 5,403 | 19 | Yes | ** | 0,009 |
| LO vs. MO | 0,4928 | 1,75 | -1,257 | 0,8438 | 9 | 3 | 2,106 | 19 | No | ns | 0,5811 |
| LO vs. AI | 0,4928 | 1,411 | -0,9184 | 0,6671 | 9 | 6 | 1,947 | 19 | No | ns | 0,649 |
| LO vs. DLO | 0,4928 | 0,9667 | -0,4739 | 0,9894 | 9 | 2 | 0,6774 | 19 | No | ns | 0,9884 |
| MO vs. AI | 1,75 | 1,411 | 0,3384 | 0,895 | 3 | 6 | 0,5347 | 19 | No | ns | 0,9953 |
| MO vs. DLO | 1,75 | 0,9667 | 0,7829 | 1,155 | 3 | 2 | 0,9582 | 19 | No | ns | 0,9589 |
| AI vs. DLO | 1,411 | 0,9667 | 0,4445 | 1,033 | 6 | 2 | 0,6083 | 19 | No | ns | 0,9923 |

**Table Analyzed** **olfactory output**

**ANOVA summary**

| | |
|---|---|
| F | 0,8898 |
| P value | 0,489 |
| P value summary | ns |
| Are differences among means s | No |
| R square | 0,1578 |

**Tukey's multiple comparisons test**

| | Mean 1 | Mean 2 | Mean Diff, | SE of diff, | n1 | n2 | q | DF | Significant? | Summary | Adjusted P Value |
|---|---|---|---|---|---|---|---|---|---|---|---|
| VO vs. LO | 1,621 | 2,92 | -1,299 | 1,601 | 4 | 9 | 1,147 | 19 | No | ns | 0,924 |
| VO vs. MO | 1,621 | 3,148 | -1,527 | 2,035 | 4 | 3 | 1,061 | 19 | No | ns | 0,9416 |
| VO vs. AI | 1,621 | 4,342 | -2,721 | 1,72 | 4 | 6 | 2,237 | 19 | No | ns | 0,5256 |
| VO vs. DLO | 1,621 | 1,154 | 0,4671 | 2,307 | 4 | 2 | 0,2863 | 19 | No | ns | 0,9996 |
| LO vs. MO | 2,92 | 3,148 | -0,228 | 1,776 | 9 | 3 | 0,1816 | 19 | No | ns | > 0,9999 |
| LO vs. AI | 2,92 | 4,342 | -1,422 | 1,404 | 9 | 6 | 1,432 | 19 | No | ns | 0,8463 |
| LO vs. DLO | 2,92 | 1,154 | 1,766 | 2,083 | 9 | 2 | 1,199 | 19 | No | ns | 0,9121 |
| MO vs. AI | 3,148 | 4,342 | -1,194 | 1,884 | 3 | 6 | 0,8965 | 19 | No | ns | 0,9676 |
| MO vs. DLO | 3,148 | 1,154 | 1,994 | 2,432 | 3 | 2 | 1,159 | 19 | No | ns | 0,9213 |
| AI vs. DLO | 4,342 | 1,154 | 3,188 | 2,175 | 6 | 2 | 2,072 | 19 | No | ns | 0,5956 |

**Table Analyzed** **gustatory output**

**ANOVA summary**

| | |
|---|---|
| F | 6,707 |
| P value | 0,0015 |
| P value summary | ** |
| Are differences among means s | Yes |
| R square | 0,5854 |

**Tukey's multiple comparisons test**

| | Mean 1 | Mean 2 | Mean Diff, | SE of diff, | n1 | n2 | q | DF | Significant? | Summary | Adjusted P Value |
|---|---|---|---|---|---|---|---|---|---|---|---|
| VO vs. LO | 0,1316 | 0,2724 | -0,1409 | 1,44 | 4 | 9 | 0,1384 | 19 | No | ns | > 0,9999 |
| VO vs. MO | 0,1316 | 0,2415 | -0,1099 | 1,83 | 4 | 3 | 0,08492 | 19 | No | ns | > 0,9999 |
| VO vs. AI | 0,1316 | 6,092 | -5,96 | 1,547 | 4 | 6 | 5,449 | 19 | Yes | ** | 0,0084 |
| VO vs. DLO | 0,1316 | 0,3266 | -0,1951 | 2,075 | 4 | 2 | 0,1329 | 19 | No | ns | > 0,9999 |
| LO vs. MO | 0,2724 | 0,2415 | 0,03098 | 1,598 | 9 | 3 | 0,02742 | 19 | No | ns | > 0,9999 |
| LO vs. AI | 0,2724 | 6,092 | -5,819 | 1,263 | 9 | 6 | 6,516 | 19 | Yes | ** | 0,0016 |
| LO vs. DLO | 0,2724 | 0,3266 | -0,0542 | 1,873 | 9 | 2 | 0,04091 | 19 | No | ns | > 0,9999 |
| MO vs. AI | 0,2415 | 6,092 | -5,85 | 1,694 | 3 | 6 | 4,883 | 19 | Yes | * | 0,0198 |
| MO vs. DLO | 0,2415 | 0,3266 | -0,08517 | 2,188 | 3 | 2 | 0,05506 | 19 | No | ns | > 0,9999 |
| AI vs. DLO | 6,092 | 0,3266 | 5,765 | 1,957 | 6 | 2 | 4,167 | 19 | No | ns | 0,0565 |
