## Supplementary 3 for "Anatomical input-output streams within mouse orbitofrontal cortex subdivisions"

| Table Analyzed | MO |
|---|---|
| Column B | VSP |
| vs. | vs, |
| Column A | DSP |
| **Paired t test** | |
| P value | 0,0226 |
| P value summary | * |
| Significantly different? (P < 0.05) | Yes |
| One- or two-tailed P value? | Two-tailed |
| t, df | t=6,546 df=2 |
| Number of pairs | 3 |

| Table Analyzed | VO |
|---|---|
| Column B | VSP |
| vs. | vs, |
| Column A | DSP |
| **Paired t test** | |
| P value | 0,012 |
| P value summary | * |
| Significantly different? (P < 0.05) | Yes |
| One- or two-tailed P value? | Two-tailed |
| t, df | t=5,473 df=3 |
| Number of pairs | 4 |

| Table Analyzed | LO |
|---|---|
| Column B | VSP |
| vs. | vs, |
| Column A | DSP |
| **Paired t test** | |
| P value | 0,0013 |
| P value summary | ** |
| Significantly different? (P < 0.05) | Yes |
| One- or two-tailed P value? | Two-tailed |
| t, df | t=4,821 df=8 |
| Number of pairs | 9 |

| Table Analyzed | AI |
|---|---|
| Column B | VSP |
| vs. | vs, |
| Column A | DSP |
| **Paired t test** | |
| P value | 0,4645 |
| P value summary | ns |
| Significantly different? (P < 0.05) | No |
| One- or two-tailed P value? | Two-tailed |
| t, df | t=0,7915 df=5 |
| Number of pairs | 6 |

| Table Analyzed | DLO |
|---|---|
| Column B | VSP |
| vs. | vs, |
| Column A | DSP |
| **Paired t test** | |
| P value | < 0,0001 |
| P value summary | **** |
| Significantly different? (P < 0.05) | Yes |
| One- or two-tailed P value? | Two-tailed |
| t, df | t=28,42 df=3 |
| Number of pairs | 4 |

**Table Analyzed** **DA**

**ANOVA summary**

| | |
|---|---|
| F | 4,823 |
| P value | 0,0171 |
| P value summary | * |
| Are differences among means statistically signi[illegible] | Yes |
| R square | 0,6369 |

**Tukey's multiple comparisons test**

| | Mean 1 | Mean 2 | Mean Diff, | SE of diff, | n1 | n2 | q | DF | Significant? | Summary | Adjusted P |
|---|---|---|---|---|---|---|---|---|---|---|---|
| VO vs. LO | 3,998 | 2,083 | 1,914 | 1,088 | 4 | 3 | 2,489 | 11 | No | ns | 0,4402 |
| VO vs. MO | 3,998 | 0,5334 | 3,464 | 1,088 | 4 | 3 | 4,505 | 11 | No | ns | 0,0541 |
| VO vs. AI | 3,998 | 0,1665 | 3,831 | 1,088 | 4 | 3 | 4,982 | 11 | Yes | * | 0,0312 |
| VO vs. DLO | 3,998 | 0,2009 | 3,797 | 1,088 | 4 | 3 | 4,937 | 11 | Yes | * | 0,0328 |
| LO vs. MO | 2,083 | 0,5334 | 1,55 | 1,163 | 3 | 3 | 1,885 | 11 | No | ns | 0,6781 |
| LO vs. AI | 2,083 | 0,1665 | 1,917 | 1,163 | 3 | 3 | 2,332 | 11 | No | ns | 0,4995 |
| LO vs. DLO | 2,083 | 0,2009 | 1,883 | 1,163 | 3 | 3 | 2,29 | 11 | No | ns | 0,5158 |
| MO vs. AI | 0,5334 | 0,1665 | 0,3669 | 1,163 | 3 | 3 | 0,4463 | 11 | No | ns | 0,9975 |
| MO vs. DLO | 0,5334 | 0,2009 | 0,3325 | 1,163 | 3 | 3 | 0,4045 | 11 | No | ns | 0,9983 |
| AI vs. DLO | 0,1665 | 0,2009 | -0,0344 | 1,163 | 3 | 3 | 0,04184 | 11 | No | ns | > 0,9999 |

**Table Analyzed** **Serotonin**

**ANOVA summary**

| | |
|---|---|
| F | 5,369 |
| P value | 0,0143 |
| P value summary | * |
| Are differences among means statistically signi[illegible] | Yes |
| R square | 0,6823 |

**Tukey's multiple comparisons test**

| | Mean 1 | Mean 2 | Mean Diff, | SE of diff, | n1 | n2 | q | DF | Significant? | Summary | Adjusted P |
|---|---|---|---|---|---|---|---|---|---|---|---|
| VO vs. LO | 1,418 | 1,604 | -0,1862 | 1,906 | 4 | 3 | 0,1382 | 10 | No | ns | > 0,9999 |
| VO vs. MO | 1,418 | 0,6493 | 0,7684 | 2,161 | 4 | 2 | 0,5028 | 10 | No | ns | 0,996 |
| VO vs. AI | 1,418 | 8,594 | -7,176 | 1,906 | 4 | 3 | 5,324 | 10 | Yes | * | 0,024 |
| VO vs. DLO | 1,418 | 0,7858 | 0,6319 | 1,906 | 4 | 3 | 0,4688 | 10 | No | ns | 0,9969 |
| LO vs. MO | 1,604 | 0,6493 | 0,9547 | 2,278 | 3 | 2 | 0,5926 | 10 | No | ns | 0,9925 |
| LO vs. AI | 1,604 | 8,594 | -6,99 | 2,038 | 3 | 3 | 4,851 | 10 | Yes | * | 0,0403 |
| LO vs. DLO | 1,604 | 0,7858 | 0,8182 | 2,038 | 3 | 3 | 0,5678 | 10 | No | ns | 0,9936 |
| MO vs. AI | 0,6493 | 8,594 | -7,944 | 2,278 | 2 | 3 | 4,931 | 10 | Yes | * | 0,0369 |
| MO vs. DLO | 0,6493 | 0,7858 | -0,1365 | 2,278 | 2 | 3 | 0,08475 | 10 | No | ns | > 0,9999 |
| AI vs. DLO | 8,594 | 0,7858 | 7,808 | 2,038 | 3 | 3 | 5,419 | 10 | Yes | * | 0,0216 |
