## Supplementary 4 for "Anatomical input-output streams within mouse orbitofrontal cortex subdivisions"

| Table Analyzed | sensory cortex inputs |
|---|---|

**ANOVA summary**

| | |
|---|---|
| F | 6,981 |
| P value | 0,006 |
| P value summary | ** |
| Are differences among means statis | Yes |
| R square | 0,7363 |

**Tukey's multiple comparisons test**

| | Mean 1 | Mean 2 | Mean Diff, | SE of diff, | n1 | n2 | q | DF | Significant | Summary | Adjusted P Value |
|---|---|---|---|---|---|---|---|---|---|---|---|
| VO vs. LO | 3,503 | 4,271 | -0,7677 | 2,442 | 4 | 3 | 0,4447 | 10 | No | ns | 0,9975 |
| VO vs. MO | 3,503 | 2,949 | 0,5547 | 2,768 | 4 | 2 | 0,2834 | 10 | No | ns | 0,9996 |
| VO vs. AI | 3,503 | 14,72 | -11,22 | 2,442 | 4 | 3 | 6,499 | 10 | Yes | ** | 0,0068 |
| VO vs. DLO | 3,503 | 4,761 | -1,257 | 2,442 | 4 | 3 | 0,7282 | 10 | No | ns | 0,9839 |
| LO vs. MO | 4,271 | 2,949 | 1,322 | 2,918 | 3 | 2 | 0,6409 | 10 | No | ns | 0,9899 |
| LO vs. AI | 4,271 | 14,72 | -10,45 | 2,61 | 3 | 3 | 5,664 | 10 | Yes | * | 0,0166 |
| LO vs. DLO | 4,271 | 4,761 | -0,4895 | 2,61 | 3 | 3 | 0,2652 | 10 | No | ns | 0,9997 |
| MO vs. AI | 2,949 | 14,72 | -11,78 | 2,918 | 2 | 3 | 5,707 | 10 | Yes | * | 0,0158 |
| MO vs. DLO | 2,949 | 4,761 | -1,812 | 2,918 | 2 | 3 | 0,8781 | 10 | No | ns | 0,9683 |
| AI vs. DLO | 14,72 | 4,761 | 9,964 | 2,61 | 3 | 3 | 5,398 | 10 | Yes | * | 0,0221 |

| Table Analyzed | Motor cortex inputs |
|---|---|

**ANOVA summary**

| | |
|---|---|
| F | 1,522 |
| P value | 0,2684 |
| P value summary | ns |
| Are differences among means statis | No |
| R square | 0,3784 |

**Tukey's multiple comparisons test**

| | Mean 1 | Mean 2 | Mean Diff, | SE of diff, | n1 | n2 | q | DF | Significant | Summary | Adjusted P Value |
|---|---|---|---|---|---|---|---|---|---|---|---|
| VO vs. LO | 2,718 | 7,314 | -4,596 | 3,584 | 4 | 3 | 1,813 | 10 | No | ns | 0,7072 |
| VO vs. MO | 2,718 | 0,5037 | 2,214 | 4,064 | 4 | 2 | 0,7705 | 10 | No | ns | 0,9802 |
| VO vs. AI | 2,718 | 1,22 | 1,498 | 3,584 | 4 | 3 | 0,5911 | 10 | No | ns | 0,9926 |
| VO vs. DLO | 2,718 | 7,96 | -5,242 | 3,584 | 4 | 3 | 2,068 | 10 | No | ns | 0,6057 |
| LO vs. MO | 7,314 | 0,5037 | 6,81 | 4,284 | 3 | 2 | 2,248 | 10 | No | ns | 0,5346 |
| LO vs. AI | 7,314 | 1,22 | 6,094 | 3,832 | 3 | 3 | 2,249 | 10 | No | ns | 0,5342 |
| LO vs. DLO | 7,314 | 7,96 | -0,6466 | 3,832 | 3 | 3 | 0,2386 | 10 | No | ns | 0,9998 |
| MO vs. AI | 0,5037 | 1,22 | -0,7161 | 4,284 | 2 | 3 | 0,2364 | 10 | No | ns | 0,9998 |
| MO vs. DLO | 0,5037 | 7,96 | -7,457 | 4,284 | 2 | 3 | 2,461 | 10 | No | ns | 0,454 |
| AI vs. DLO | 1,22 | 7,96 | -6,74 | 3,832 | 3 | 3 | 2,488 | 10 | No | ns | 0,4444 |

| Table Analyzed | Prefrontal cortex inputs |
|---|---|

**ANOVA summary**

| | |
|---|---|
| F | 6,894 |
| P value | 0,0062 |
| P value summary | ** |
| Are differences among means statis | Yes |
| R square | 0,7339 |

**Tukey's multiple comparisons test**

| | Mean 1 | Mean 2 | Mean Diff, | SE of diff, | n1 | n2 | q | DF | Significant | Summary | Adjusted P Value |
|---|---|---|---|---|---|---|---|---|---|---|---|
| VO vs. LO | 9,472 | 6,124 | 3,348 | 2,622 | 4 | 3 | 1,806 | 10 | No | ns | 0,7102 |
| VO vs. MO | 9,472 | 20,29 | -10,82 | 2,973 | 4 | 2 | 5,145 | 10 | Yes | * | 0,0292 |
| VO vs. AI | 9,472 | 5,658 | 3,814 | 2,622 | 4 | 3 | 2,057 | 10 | No | ns | 0,6102 |
| VO vs. DLO | 9,472 | 7,09 | 2,382 | 2,622 | 4 | 3 | 1,285 | 10 | No | ns | 0,8873 |
| LO vs. MO | 6,124 | 20,29 | -14,16 | 3,134 | 3 | 2 | 6,392 | 10 | Yes | ** | 0,0076 |
| LO vs. AI | 6,124 | 5,658 | 0,4667 | 2,803 | 3 | 3 | 0,2355 | 10 | No | ns | 0,9998 |
| LO vs. DLO | 6,124 | 7,09 | -0,9654 | 2,803 | 3 | 3 | 0,487 | 10 | No | ns | 0,9965 |
| MO vs. AI | 20,29 | 5,658 | 14,63 | 3,134 | 2 | 3 | 6,602 | 10 | Yes | ** | 0,0061 |
| MO vs. DLO | 20,29 | 7,09 | 13,2 | 3,134 | 2 | 3 | 5,956 | 10 | Yes | * | 0,0121 |
| AI vs. DLO | 5,658 | 7,09 | -1,432 | 2,803 | 3 | 3 | 0,7225 | 10 | No | ns | 0,9843 |

| Table Analyzed | Association cortex inputs |
|---|---|

**ANOVA summary**

| | |
|---|---|
| F | 2,543 |
| P value | 0,1054 |
| P value summary | ns |
| Are differences among means statis | No |
| R square | 0,5043 |

**Tukey's multiple comparisons test**

| | Mean 1 | Mean 2 | Mean Diff, | SE of diff, | n1 | n2 | q | DF | Significant? | Summary | Adjusted P Value |
|---|---|---|---|---|---|---|---|---|---|---|---|
| VO vs. LO | 7,089 | 7,668 | -0,5791 | 2,439 | 4 | 3 | 0,3357 | 10 | No | ns | 0,9992 |
| VO vs. MO | 7,089 | 13,71 | -6,616 | 2,766 | 4 | 2 | 3,383 | 10 | No | ns | 0,1945 |
| VO vs. AI | 7,089 | 12,88 | -5,788 | 2,439 | 4 | 3 | 3,356 | 10 | No | ns | 0,1999 |
| VO vs. DLO | 7,089 | 8,911 | -1,823 | 2,439 | 4 | 3 | 1,057 | 10 | No | ns | 0,94 |
| LO vs. MO | 7,668 | 13,71 | -6,037 | 2,916 | 3 | 2 | 2,928 | 10 | No | ns | 0,3026 |
| LO vs. AI | 7,668 | 12,88 | -5,209 | 2,608 | 3 | 3 | 2,825 | 10 | No | ns | 0,3327 |
| LO vs. DLO | 7,668 | 8,911 | -1,244 | 2,608 | 3 | 3 | 0,6745 | 10 | No | ns | 0,9878 |
| MO vs. AI | 13,71 | 12,88 | 0,8283 | 2,916 | 2 | 3 | 0,4018 | 10 | No | ns | 0,9983 |
| MO vs. DLO | 13,71 | 8,911 | 4,794 | 2,916 | 2 | 3 | 2,325 | 10 | No | ns | 0,5049 |
| AI vs. DLO | 12,88 | 8,911 | 3,965 | 2,608 | 3 | 3 | 2,15 | 10 | No | ns | 0,573 |

**Table Analyzed** **Olfactory inputs**

**ANOVA summary**

| | |
|---|---|
| F | 7,108 |
| P value | 0,0056 |
| P value summary | ** |
| Are differences among means statis | Yes |
| R square | 0,7398 |

**Tukey's multiple comparisons test**

| | Mean 1 | Mean 2 | Mean Diff, | SE of diff, | n1 | n2 | q | DF | Significant? | Summary | Adjusted P Value |
|---|---|---|---|---|---|---|---|---|---|---|---|
| VO vs. LO | 0,34 | 3,755 | -3,415 | 2,374 | 4 | 3 | 2,035 | 10 | No | ns | 0,6191 |
| VO vs. MO | 0,34 | 1,543 | -1,203 | 2,692 | 4 | 2 | 0,6323 | 10 | No | ns | 0,9904 |
| VO vs. AI | 0,34 | 12,24 | -11,9 | 2,374 | 4 | 3 | 7,088 | 10 | Yes | ** | 0,0037 |
| VO vs. DLO | 0,34 | 6,058 | -5,718 | 2,374 | 4 | 3 | 3,407 | 10 | No | ns | 0,1899 |
| LO vs. MO | 3,755 | 1,543 | 2,212 | 2,837 | 3 | 2 | 1,103 | 10 | No | ns | 0,9308 |
| LO vs. AI | 3,755 | 12,24 | -8,482 | 2,538 | 3 | 3 | 4,727 | 10 | Yes | * | 0,0462 |
| LO vs. DLO | 3,755 | 6,058 | -2,303 | 2,538 | 3 | 3 | 1,283 | 10 | No | ns | 0,8877 |
| MO vs. AI | 1,543 | 12,24 | -10,69 | 2,837 | 2 | 3 | 5,331 | 10 | Yes | * | 0,0238 |
| MO vs. DLO | 1,543 | 6,058 | -4,515 | 2,837 | 2 | 3 | 2,25 | 10 | No | ns | 0,5337 |
| AI vs. DLO | 12,24 | 6,058 | 6,179 | 2,538 | 3 | 3 | 3,444 | 10 | No | ns | 0,1828 |

**Table Analyzed** **Claustrum inputs**

**ANOVA summary**

| | |
|---|---|
| F | 0,624 |
| P value | 0,656 |
| P value summary | ns |
| Are differences among means statis | No |
| R square | 0,1997 |

**Tukey's multiple comparisons test**

| | Mean 1 | Mean 2 | Mean Diff, | SE of diff, | n1 | n2 | q | DF | Significant? | Summary | Adjusted P Value |
|---|---|---|---|---|---|---|---|---|---|---|---|
| VO vs. LO | 6,018 | 8,697 | -2,68 | 4,507 | 4 | 3 | 0,8409 | 10 | No | ns | 0,9728 |
| VO vs. MO | 6,018 | 7,645 | -1,627 | 5,11 | 4 | 2 | 0,4504 | 10 | No | ns | 0,9974 |
| VO vs. AI | 6,018 | 10,61 | -4,589 | 4,507 | 4 | 3 | 1,44 | 10 | No | ns | 0,8416 |
| VO vs. DLO | 6,018 | 12,64 | -6,617 | 4,507 | 4 | 3 | 2,077 | 10 | No | ns | 0,6024 |
| LO vs. MO | 8,697 | 7,645 | 1,052 | 5,386 | 3 | 2 | 0,2763 | 10 | No | ns | 0,9996 |
| LO vs. AI | 8,697 | 10,61 | -1,909 | 4,818 | 3 | 3 | 0,5603 | 10 | No | ns | 0,9939 |
| LO vs. DLO | 8,697 | 12,64 | -3,938 | 4,818 | 3 | 3 | 1,156 | 10 | No | ns | 0,9193 |
| MO vs. AI | 7,645 | 10,61 | -2,961 | 5,386 | 2 | 3 | 0,7774 | 10 | No | ns | 0,9795 |
| MO vs. DLO | 7,645 | 12,64 | -4,99 | 5,386 | 2 | 3 | 1,31 | 10 | No | ns | 0,8804 |
| AI vs. DLO | 10,61 | 12,64 | -2,029 | 4,818 | 3 | 3 | 0,5955 | 10 | No | ns | 0,9924 |

**Table Analyzed** **Striatum inputs**

**ANOVA summary**

| | |
|---|---|
| F | 2,498 |
| P value | 0,1096 |
| P value summary | ns |

| | |
|---|---|
| Are differences among means statis | No |
| R square | 0,4998 |

**Tukey's multiple comparisons test**

| | Mean 1 | Mean 2 | Mean Diff, | SE of diff, | n1 | n2 | q | DF | Significant? | Summary | Adjusted P Value |
|---|---|---|---|---|---|---|---|---|---|---|---|
| VO vs. LO | 4,569 | 5,664 | -1,095 | 1,339 | 4 | 3 | 1,157 | 10 | No | ns | 0,9191 |
| VO vs. MO | 4,569 | 2,873 | 1,696 | 1,518 | 4 | 2 | 1,58 | 10 | No | ns | 0,7946 |
| VO vs. AI | 4,569 | 6,63 | -2,061 | 1,339 | 4 | 3 | 2,176 | 10 | No | ns | 0,5628 |
| VO vs. DLO | 4,569 | 7,283 | -2,714 | 1,339 | 4 | 3 | 2,867 | 10 | No | ns | 0,3203 |
| LO vs. MO | 5,664 | 2,873 | 2,791 | 1,6 | 3 | 2 | 2,466 | 10 | No | ns | 0,4521 |
| LO vs. AI | 5,664 | 6,63 | -0,9651 | 1,432 | 3 | 3 | 0,9535 | 10 | No | ns | 0,9577 |
| LO vs. DLO | 5,664 | 7,283 | -1,619 | 1,432 | 3 | 3 | 1,599 | 10 | No | ns | 0,7876 |
| MO vs. AI | 2,873 | 6,63 | -3,756 | 1,6 | 2 | 3 | 3,319 | 10 | No | ns | 0,2074 |
| MO vs. DLO | 2,873 | 7,283 | -4,41 | 1,6 | 2 | 3 | 3,897 | 10 | No | ns | 0,1139 |
| AI vs. DLO | 6,63 | 7,283 | -0,6537 | 1,432 | 3 | 3 | 0,6458 | 10 | No | ns | 0,9897 |

**Table Analyzed** **Hippocampus inputs**

**ANOVA summary**

| | |
|---|---|
| F | 17,83 |
| P value | 0,0002 |
| P value summary | *** |
| Are differences among means statis | Yes |
| R square | 0,877 |

**Tukey's multiple comparisons test**

| | Mean 1 | Mean 2 | Mean Diff, | SE of diff, | n1 | n2 | q | DF | Significant? | Summary | Adjusted P Value |
|---|---|---|---|---|---|---|---|---|---|---|---|
| VO vs. LO | 0,5784 | 0,07276 | 0,5056 | 0,1013 | 4 | 3 | 7,057 | 10 | Yes | ** | 0,0038 |
| VO vs. MO | 0,5784 | 0,7232 | -0,1448 | 0,1149 | 4 | 2 | 1,783 | 10 | No | ns | 0,719 |
| VO vs. AI | 0,5784 | 0,06216 | 0,5162 | 0,1013 | 4 | 3 | 7,205 | 10 | Yes | ** | 0,0033 |
| VO vs. DLO | 0,5784 | 0 | 0,5784 | 0,1013 | 4 | 3 | 8,073 | 10 | Yes | ** | 0,0014 |
| LO vs. MO | 0,07276 | 0,7232 | -0,6504 | 0,1211 | 3 | 2 | 7,596 | 10 | Yes | ** | 0,0023 |
| LO vs. AI | 0,07276 | 0,06216 | 0,0106 | 0,1083 | 3 | 3 | 0,1384 | 10 | No | ns | > 0,9999 |
| LO vs. DLO | 0,07276 | 0 | 0,07276 | 0,1083 | 3 | 3 | 0,95 | 10 | No | ns | 0,9582 |
| MO vs. AI | 0,7232 | 0,06216 | 0,661 | 0,1211 | 2 | 3 | 7,72 | 10 | Yes | ** | 0,002 |
| MO vs. DLO | 0,7232 | 0 | 0,7232 | 0,1211 | 2 | 3 | 8,446 | 10 | Yes | ** | 0,001 |
| AI vs. DLO | 0,06216 | 0 | 0,06216 | 0,1083 | 3 | 3 | 0,8116 | 10 | No | ns | 0,976 |

**Table Analyzed** **Amygdala inputs**

**ANOVA summary**

| | |
|---|---|
| F | 2,435 |
| P value | 0,1157 |
| P value summary | ns |
| Are differences among means statis | No |
| R square | 0,4934 |

**Tukey's multiple comparisons test**

| | Mean 1 | Mean 2 | Mean Diff, | SE of diff, | n1 | n2 | q | DF | Significant? | Summary | Adjusted P Value |
|---|---|---|---|---|---|---|---|---|---|---|---|
| VO vs. LO | 6,647 | 4,454 | 2,193 | 1,904 | 4 | 3 | 1,629 | 10 | No | ns | 0,7768 |
| VO vs. MO | 6,647 | 9,629 | -2,981 | 2,159 | 4 | 2 | 1,953 | 10 | No | ns | 0,6519 |
| VO vs. AI | 6,647 | 9,187 | -2,54 | 1,904 | 4 | 3 | 1,886 | 10 | No | ns | 0,6784 |
| VO vs. DLO | 6,647 | 9,563 | -2,915 | 1,904 | 4 | 3 | 2,165 | 10 | No | ns | 0,5671 |
| LO vs. MO | 4,454 | 9,629 | -5,175 | 2,276 | 3 | 2 | 3,216 | 10 | No | ns | 0,2298 |
| LO vs. AI | 4,454 | 9,187 | -4,733 | 2,036 | 3 | 3 | 3,288 | 10 | No | ns | 0,2139 |
| LO vs. DLO | 4,454 | 9,563 | -5,109 | 2,036 | 3 | 3 | 3,549 | 10 | No | ns | 0,1641 |
| MO vs. AI | 9,629 | 9,187 | 0,4416 | 2,276 | 2 | 3 | 0,2744 | 10 | No | ns | 0,9996 |
| MO vs. DLO | 9,629 | 9,563 | 0,06609 | 2,276 | 2 | 3 | 0,04106 | 10 | No | ns | > 0,9999 |
| AI vs. DLO | 9,187 | 9,563 | -0,3756 | 2,036 | 3 | 3 | 0,2609 | 10 | No | ns | 0,9997 |

**Table Analyzed** **Hypothalamus inputs**

**ANOVA summary**

| | |
|---|---|
| F | 2,473 |
| P value | 0,112 |
| P value summary | ns |
| Are differences among means statis | No |
| R square | 0,4973 |

**Tukey's multiple comparisons test**

| | Mean 1 | Mean 2 | Mean Diff, | SE of diff, | n1 | n2 | q | DF | Significant? | Summary | Adjusted P Value |
|---|---|---|---|---|---|---|---|---|---|---|---|
| VO vs. LO | 0,9411 | 0,4891 | 0,452 | 0,4702 | 4 | 3 | 1,359 | 10 | No | ns | 0,8662 |
| VO vs. MO | 0,9411 | 1,578 | -0,6371 | 0,5331 | 4 | 2 | 1,69 | 10 | No | ns | 0,7543 |
| VO vs. AI | 0,9411 | 0,2671 | 0,674 | 0,4702 | 4 | 3 | 2,027 | 10 | No | ns | 0,6222 |
| VO vs. DLO | 0,9411 | 0,0214 | 0,9197 | 0,4702 | 4 | 3 | 2,766 | 10 | No | ns | 0,3506 |
| LO vs. MO | 0,4891 | 1,578 | -1,089 | 0,562 | 3 | 2 | 2,741 | 10 | No | ns | 0,3587 |
| LO vs. AI | 0,4891 | 0,2671 | 0,222 | 0,5026 | 3 | 3 | 0,6246 | 10 | No | ns | 0,9909 |
| LO vs. DLO | 0,4891 | 0,0214 | 0,4677 | 0,5026 | 3 | 3 | 1,316 | 10 | No | ns | 0,8787 |
| MO vs. AI | 1,578 | 0,2671 | 1,311 | 0,562 | 2 | 3 | 3,299 | 10 | No | ns | 0,2115 |
| MO vs. DLO | 1,578 | 0,0214 | 1,557 | 0,562 | 2 | 3 | 3,918 | 10 | No | ns | 0,1113 |
| AI vs. DLO | 0,2671 | 0,0214 | 0,2457 | 0,5026 | 3 | 3 | 0,6914 | 10 | No | ns | 0,9867 |

**Table Analyzed** **Thalamus inputs**

**ANOVA summary**

| | |
|---|---|
| F | 4,174 |
| P value | 0,0304 |
| P value summary | * |
| Are differences among means statis | Yes |
| R square | 0,6254 |

**Tukey's multiple comparisons test**

| | Mean 1 | Mean 2 | Mean Diff, | SE of diff, | n1 | n2 | q | DF | Significant? | Summary | Adjusted P Value |
|---|---|---|---|---|---|---|---|---|---|---|---|
| VO vs. LO | 52,6 | 47,44 | 5,163 | 9,09 | 4 | 3 | 0,8033 | 10 | No | ns | 0,9769 |
| VO vs. MO | 52,6 | 37,38 | 15,22 | 10,31 | 4 | 2 | 2,088 | 10 | No | ns | 0,5977 |
| VO vs. AI | 52,6 | 17,77 | 34,83 | 9,09 | 4 | 3 | 5,419 | 10 | Yes | * | 0,0216 |
| VO vs. DLO | 52,6 | 34,73 | 17,87 | 9,09 | 4 | 3 | 2,78 | 10 | No | ns | 0,3464 |
| LO vs. MO | 47,44 | 37,38 | 10,06 | 10,86 | 3 | 2 | 1,309 | 10 | No | ns | 0,8806 |
| LO vs. AI | 47,44 | 17,77 | 29,66 | 9,718 | 3 | 3 | 4,317 | 10 | No | ns | 0,0723 |
| LO vs. DLO | 47,44 | 34,73 | 12,71 | 9,718 | 3 | 3 | 1,849 | 10 | No | ns | 0,6931 |
| MO vs. AI | 37,38 | 17,77 | 19,61 | 10,86 | 2 | 3 | 2,552 | 10 | No | ns | 0,4215 |
| MO vs. DLO | 37,38 | 34,73 | 2,649 | 10,86 | 2 | 3 | 0,3448 | 10 | No | ns | 0,9991 |
| AI vs. DLO | 17,77 | 34,73 | -16,96 | 9,718 | 3 | 3 | 2,468 | 10 | No | ns | 0,4515 |

**Table Analyzed** **Brainstem inputs**

**ANOVA summary**

| | |
|---|---|
| F | 3,194 |
| P value | 0,0621 |
| P value summary | ns |
| Are differences among means statis | No |
| R square | 0,5609 |

**Tukey's multiple comparisons test**

| | Mean 1 | Mean 2 | Mean Diff, | SE of diff, | n1 | n2 | q | DF | Significant? | Summary | Adjusted P Value |
|---|---|---|---|---|---|---|---|---|---|---|---|
| VO vs. LO | 5,525 | 4,054 | 1,471 | 2,322 | 4 | 3 | 0,8959 | 10 | No | ns | 0,966 |
| VO vs. MO | 5,525 | 1,183 | 4,342 | 2,633 | 4 | 2 | 2,332 | 10 | No | ns | 0,5022 |
| VO vs. AI | 5,525 | 8,76 | -3,235 | 2,322 | 4 | 3 | 1,97 | 10 | No | ns | 0,6451 |
| VO vs. DLO | 5,525 | 0,9867 | 4,538 | 2,322 | 4 | 3 | 2,764 | 10 | No | ns | 0,3515 |
| LO vs. MO | 4,054 | 1,183 | 2,871 | 2,776 | 3 | 2 | 1,463 | 10 | No | ns | 0,8342 |
| LO vs. AI | 4,054 | 8,76 | -4,706 | 2,483 | 3 | 3 | 2,681 | 10 | No | ns | 0,3779 |
| LO vs. DLO | 4,054 | 0,9867 | 3,067 | 2,483 | 3 | 3 | 1,747 | 10 | No | ns | 0,7328 |
| MO vs. AI | 1,183 | 8,76 | -7,577 | 2,776 | 2 | 3 | 3,861 | 10 | No | ns | 0,1183 |
| MO vs. DLO | 1,183 | 0,9867 | 0,196 | 2,776 | 2 | 3 | 0,09987 | 10 | No | ns | > 0,9999 |
| AI vs. DLO | 8,76 | 0,9867 | 7,773 | 2,483 | 3 | 3 | 4,428 | 10 | No | ns | 0,0641 |

**Table Analyzed** **Sensory cortex output**

**ANOVA summary**

| | |
|---|---|
| F | 4,008 |
| P value | 0,016 |
| P value summary | * |
| Are differences among means statist | Yes |
| R square | 0,4576 |

**Tukey's multiple comparisons test**

| | Mean 1 | Mean 2 | Mean Diff, | SE of diff, | n1 | n2 | q | DF | Significant | Summary | Adjusted P Value |
|---|---|---|---|---|---|---|---|---|---|---|---|
| VO vs. LO | 26,19 | 3,653 | 22,54 | 5,661 | 4 | 9 | 5,63 | 19 | Yes | ** | 0,0063 |
| VO vs. MO | 26,19 | 8,934 | 17,26 | 7,195 | 4 | 3 | 3,392 | 19 | No | ns | 0,1587 |
| VO vs. AI | 26,19 | 10,21 | 15,98 | 6,081 | 4 | 6 | 3,716 | 19 | No | ns | 0,1047 |
| VO vs. DLO | 26,19 | 7,966 | 18,22 | 8,158 | 4 | 2 | 3,159 | 19 | No | ns | 0,21 |
| LO vs. MO | 3,653 | 8,934 | -5,281 | 6,28 | 9 | 3 | 1,189 | 19 | No | ns | 0,9144 |
| LO vs. AI | 3,653 | 10,21 | -6,558 | 4,965 | 9 | 6 | 1,868 | 19 | No | ns | 0,6822 |
| LO vs. DLO | 3,653 | 7,966 | -4,313 | 7,364 | 9 | 2 | 0,8282 | 19 | No | ns | 0,9756 |
| MO vs. AI | 8,934 | 10,21 | -1,277 | 6,661 | 3 | 6 | 0,2711 | 19 | No | ns | 0,9997 |
| MO vs. DLO | 8,934 | 7,966 | 0,9681 | 8,6 | 3 | 2 | 0,1592 | 19 | No | ns | > 0,9999 |
| AI vs. DLO | 10,21 | 7,966 | 2,245 | 7,692 | 6 | 2 | 0,4127 | 19 | No | ns | 0,9983 |

**Table Analyzed** **Motor cortex output**

**ANOVA summary**

| | |
|---|---|
| F | 2,047 |
| P value | 0,1284 |
| P value summary | ns |
| Are differences among means statist | No |
| R square | 0,3011 |

**Tukey's multiple comparisons test**

| | Mean 1 | Mean 2 | Mean Diff, | SE of diff, | n1 | n2 | q | DF | Significant | Summary | Adjusted P Value |
|---|---|---|---|---|---|---|---|---|---|---|---|
| VO vs. LO | 1,423 | 4,2 | -2,777 | 1,86 | 4 | 9 | 2,112 | 19 | No | ns | 0,5787 |
| VO vs. MO | 1,423 | 0,9539 | 0,4692 | 2,364 | 4 | 3 | 0,2807 | 19 | No | ns | 0,9996 |
| VO vs. AI | 1,423 | 2,834 | -1,411 | 1,998 | 4 | 6 | 0,9986 | 19 | No | ns | 0,9526 |
| VO vs. DLO | 1,423 | 7,727 | -6,304 | 2,68 | 4 | 2 | 3,327 | 19 | No | ns | 0,1719 |
| LO vs. MO | 4,2 | 0,9539 | 3,246 | 2,063 | 9 | 3 | 2,225 | 19 | No | ns | 0,5307 |
| LO vs. AI | 4,2 | 2,834 | 1,367 | 1,631 | 9 | 6 | 1,185 | 19 | No | ns | 0,9154 |
| LO vs. DLO | 4,2 | 7,727 | -3,527 | 2,419 | 9 | 2 | 2,062 | 19 | No | ns | 0,6001 |
| MO vs. AI | 0,9539 | 2,834 | -1,88 | 2,188 | 3 | 6 | 1,215 | 19 | No | ns | 0,9082 |
| MO vs. DLO | 0,9539 | 7,727 | -6,773 | 2,825 | 3 | 2 | 3,391 | 19 | No | ns | 0,1588 |
| AI vs. DLO | 2,834 | 7,727 | -4,893 | 2,527 | 6 | 2 | 2,739 | 19 | No | ns | 0,3328 |

**Table Analyzed** **Prefrontal cortex output**

**ANOVA summary**

| | |
|---|---|
| F | 1,685 |
| P value | 0,195 |
| P value summary | ns |
| Are differences among means statist | No |
| R square | 0,2618 |

**Tukey's multiple comparisons test**

| | Mean 1 | Mean 2 | Mean Diff, | SE of diff, | n1 | n2 | q | DF | Significant | Summary | Adjusted P Value |
|---|---|---|---|---|---|---|---|---|---|---|---|
| VO vs. LO | 10,3 | 16 | -5,697 | 7,093 | 4 | 9 | 1,136 | 19 | No | ns | 0,9264 |
| VO vs. MO | 10,3 | 30,24 | -19,94 | 9,015 | 4 | 3 | 3,128 | 19 | No | ns | 0,2177 |
| VO vs. AI | 10,3 | 13,51 | -3,202 | 7,619 | 4 | 6 | 0,5944 | 19 | No | ns | 0,9929 |
| VO vs. DLO | 10,3 | 25,7 | -15,39 | 10,22 | 4 | 2 | 2,13 | 19 | No | ns | 0,5712 |
| LO vs. MO | 16 | 30,24 | -14,24 | 7,869 | 9 | 3 | 2,56 | 19 | No | ns | 0,3969 |
| LO vs. AI | 16 | 13,51 | 2,495 | 6,221 | 9 | 6 | 0,5671 | 19 | No | ns | 0,9941 |
| LO vs. DLO | 16 | 25,7 | -9,697 | 9,227 | 9 | 2 | 1,486 | 19 | No | ns | 0,8286 |
| MO vs. AI | 30,24 | 13,51 | 16,74 | 8,347 | 3 | 6 | 2,836 | 19 | No | ns | 0,301 |
| MO vs. DLO | 30,24 | 25,7 | 4,545 | 10,78 | 3 | 2 | 0,5965 | 19 | No | ns | 0,9928 |
| AI vs. DLO | 13,51 | 25,7 | -12,19 | 9,638 | 6 | 2 | 1,789 | 19 | No | ns | 0,7147 |

**Table Analyzed** **Association cortex output**

**ANOVA summary**

| | |
|---|---|
| F | 4,347 |

| | |
|---|---|
| P value | 0,0115 |
| P value summary | * |
| Are differences among means statist | Yes |
| R square | 0,4779 |

**Tukey's multiple comparisons test**

| | Mean 1 | Mean 2 | Mean Diff, | SE of diff, | n1 | n2 | q | DF | Significant | Summary | Adjusted P Value |
|---|---|---|---|---|---|---|---|---|---|---|---|
| VO vs. LO | 6,064 | 2,284 | 3,78 | 2,422 | 4 | 9 | 2,207 | 19 | No | ns | 0,5385 |
| VO vs. MO | 6,064 | 11,44 | -5,379 | 3,079 | 4 | 3 | 2,471 | 19 | No | ns | 0,4309 |
| VO vs. AI | 6,064 | 9,014 | -2,95 | 2,602 | 4 | 6 | 1,603 | 19 | No | ns | 0,787 |
| VO vs. DLO | 6,064 | 3,2 | 2,864 | 3,491 | 4 | 2 | 1,16 | 19 | No | ns | 0,9211 |
| LO vs. MO | 2,284 | 11,44 | -9,159 | 2,688 | 9 | 3 | 4,82 | 19 | Yes | * | 0,0218 |
| LO vs. AI | 2,284 | 9,014 | -6,731 | 2,125 | 9 | 6 | 4,48 | 19 | Yes | * | 0,036 |
| LO vs. DLO | 2,284 | 3,2 | -0,9161 | 3,151 | 9 | 2 | 0,4111 | 19 | No | ns | 0,9983 |
| MO vs. AI | 11,44 | 9,014 | 2,428 | 2,851 | 3 | 6 | 1,205 | 19 | No | ns | 0,9107 |
| MO vs. DLO | 11,44 | 3,2 | 8,243 | 3,68 | 3 | 2 | 3,168 | 19 | No | ns | 0,2079 |
| AI vs. DLO | 9,014 | 3,2 | 5,815 | 3,292 | 6 | 2 | 2,498 | 19 | No | ns | 0,4202 |

**Table Analyzed** **Olfactory areas output**

**ANOVA summary**

| | |
|---|---|
| F | 0,8257 |
| P value | 0,525 |
| P value summary | ns |
| Are differences among means statist | No |
| R square | 0,1481 |

**Tukey's multiple comparisons test**

| | Mean 1 | Mean 2 | Mean Diff, | SE of diff, | n1 | n2 | q | DF | Significant | Summary | Adjusted P Value |
|---|---|---|---|---|---|---|---|---|---|---|---|
| VO vs. LO | 1,397 | 2,74 | -1,343 | 1,544 | 4 | 9 | 1,23 | 19 | No | ns | 0,9043 |
| VO vs. MO | 1,397 | 2,868 | -1,471 | 1,962 | 4 | 3 | 1,06 | 19 | No | ns | 0,9417 |
| VO vs. AI | 1,397 | 3,959 | -2,563 | 1,658 | 4 | 6 | 2,185 | 19 | No | ns | 0,5476 |
| VO vs. DLO | 1,397 | 1,058 | 0,3389 | 2,225 | 4 | 2 | 0,2154 | 19 | No | ns | 0,9999 |
| LO vs. MO | 2,74 | 2,868 | -0,1278 | 1,713 | 9 | 3 | 0,1056 | 19 | No | ns | > 0,9999 |
| LO vs. AI | 2,74 | 3,959 | -1,219 | 1,354 | 9 | 6 | 1,273 | 19 | No | ns | 0,8932 |
| LO vs. DLO | 2,74 | 1,058 | 1,682 | 2,008 | 9 | 2 | 1,185 | 19 | No | ns | 0,9155 |
| MO vs. AI | 2,868 | 3,959 | -1,091 | 1,817 | 3 | 6 | 0,8496 | 19 | No | ns | 0,9733 |
| MO vs. DLO | 2,868 | 1,058 | 1,81 | 2,345 | 3 | 2 | 1,091 | 19 | No | ns | 0,9357 |
| AI vs. DLO | 3,959 | 1,058 | 2,901 | 2,098 | 6 | 2 | 1,956 | 19 | No | ns | 0,6451 |

**Table Analyzed** **Claustrum output**

**ANOVA summary**

| | |
|---|---|
| F | 2,777 |
| P value | 0,0568 |
| P value summary | ns |
| Are differences among means statist | No |
| R square | 0,3689 |

**Tukey's multiple comparisons test**

| | Mean 1 | Mean 2 | Mean Diff, | SE of diff, | n1 | n2 | q | DF | Significant | Summary | Adjusted P Value |
|---|---|---|---|---|---|---|---|---|---|---|---|
| VO vs. LO | 2,69 | 3,533 | -0,8433 | 1,996 | 4 | 9 | 0,5976 | 19 | No | ns | 0,9928 |
| VO vs. MO | 2,69 | 6,655 | -3,965 | 2,536 | 4 | 3 | 2,211 | 19 | No | ns | 0,5367 |
| VO vs. AI | 2,69 | 8,322 | -5,632 | 2,144 | 4 | 6 | 3,716 | 19 | No | ns | 0,1047 |
| VO vs. DLO | 2,69 | 3,075 | -0,3855 | 2,876 | 4 | 2 | 0,1896 | 19 | No | ns | > 0,9999 |
| LO vs. MO | 3,533 | 6,655 | -3,122 | 2,214 | 9 | 3 | 1,994 | 19 | No | ns | 0,6288 |
| LO vs. AI | 3,533 | 8,322 | -4,789 | 1,75 | 9 | 6 | 3,87 | 19 | No | ns | 0,0852 |
| LO vs. DLO | 3,533 | 3,075 | 0,4577 | 2,596 | 9 | 2 | 0,2494 | 19 | No | ns | 0,9998 |
| MO vs. AI | 6,655 | 8,322 | -1,667 | 2,348 | 3 | 6 | 1,004 | 19 | No | ns | 0,9517 |
| MO vs. DLO | 6,655 | 3,075 | 3,58 | 3,032 | 3 | 2 | 1,67 | 19 | No | ns | 0,7618 |
| AI vs. DLO | 8,322 | 3,075 | 5,247 | 2,711 | 6 | 2 | 2,737 | 19 | No | ns | 0,3336 |

**Table Analyzed** **Striatum output**

**ANOVA summary**

| | |
|---|---|
| F | 2,213 |
| P value | 0,1062 |
| P value summary | ns |
| Are differences among means statist | No |
| R square | 0,3178 |

**Tukey's multiple comparisons test**

| | Mean 1 | Mean 2 | Mean Diff, | SE of diff, | n1 | n2 | q | DF | Significant | Summary | Adjusted P Value |
|---|---|---|---|---|---|---|---|---|---|---|---|
| VO vs. LO | 5,311 | 8,122 | -2,81 | 2,906 | 4 | 9 | 1,368 | 19 | No | ns | 0,8664 |
| VO vs. MO | 5,311 | 8,653 | -3,342 | 3,693 | 4 | 3 | 1,28 | 19 | No | ns | 0,8915 |
| VO vs. AI | 5,311 | 13,94 | -8,628 | 3,121 | 4 | 6 | 3,909 | 19 | No | ns | 0,0808 |
| VO vs. DLO | 5,311 | 8,778 | -3,467 | 4,188 | 4 | 2 | 1,171 | 19 | No | ns | 0,9186 |
| LO vs. MO | 8,122 | 8,653 | -0,5313 | 3,224 | 9 | 3 | 0,2331 | 19 | No | ns | 0,9998 |
| LO vs. AI | 8,122 | 13,94 | -5,817 | 2,548 | 9 | 6 | 3,228 | 19 | No | ns | 0,1935 |
| LO vs. DLO | 8,122 | 8,778 | -0,6566 | 3,78 | 9 | 2 | 0,2457 | 19 | No | ns | 0,9998 |
| MO vs. AI | 8,653 | 13,94 | -5,286 | 3,419 | 3 | 6 | 2,186 | 19 | No | ns | 0,5471 |
| MO vs. DLO | 8,653 | 8,778 | -0,1253 | 4,414 | 3 | 2 | 0,04015 | 19 | No | ns | > 0,9999 |
| AI vs. DLO | 13,94 | 8,778 | 5,161 | 3,948 | 6 | 2 | 1,849 | 19 | No | ns | 0,6902 |

**Table Analyzed** **Hippocampus output**

**ANOVA summary**

| | |
|---|---|
| F | 0,3627 |
| P value | 0,832 |
| P value summary | ns |
| Are differences among means statist | No |
| R square | 0,07095 |

**Tukey's multiple comparisons test**

| | Mean 1 | Mean 2 | Mean Diff, | SE of diff, | n1 | n2 | q | DF | Significant | Summary | Adjusted P Value |
|---|---|---|---|---|---|---|---|---|---|---|---|
| VO vs. LO | 0,01064 | 0,05969 | -0,04905 | 0,04851 | 4 | 9 | 1,43 | 19 | No | ns | 0,8471 |
| VO vs. MO | 0,01064 | 0,03402 | -0,02338 | 0,06165 | 4 | 3 | 0,5363 | 19 | No | ns | 0,9952 |
| VO vs. AI | 0,01064 | 0,03486 | -0,02422 | 0,0521 | 4 | 6 | 0,6574 | 19 | No | ns | 0,9896 |
| VO vs. DLO | 0,01064 | 0,005417 | 0,005225 | 0,0699 | 4 | 2 | 0,1057 | 19 | No | ns | > 0,9999 |
| LO vs. MO | 0,05969 | 0,03402 | 0,02567 | 0,05381 | 9 | 3 | 0,6746 | 19 | No | ns | 0,9886 |
| LO vs. AI | 0,05969 | 0,03486 | 0,02483 | 0,04254 | 9 | 6 | 0,8254 | 19 | No | ns | 0,9759 |
| LO vs. DLO | 0,05969 | 0,005417 | 0,05427 | 0,0631 | 9 | 2 | 1,216 | 19 | No | ns | 0,9078 |
| MO vs. AI | 0,03402 | 0,03486 | -0,00084 | 0,05708 | 3 | 6 | 0,02083 | 19 | No | ns | > 0,9999 |
| MO vs. DLO | 0,03402 | 0,005417 | 0,0286 | 0,07368 | 3 | 2 | 0,549 | 19 | No | ns | 0,9948 |
| AI vs. DLO | 0,03486 | 0,005417 | 0,02944 | 0,06591 | 6 | 2 | 0,6318 | 19 | No | ns | 0,9911 |

**Table Analyzed** **Amygdala output**

**ANOVA summary**

| | |
|---|---|
| F | 15,51 |
| P value | < 0,0001 |
| P value summary | **** |
| Are differences among means statist | Yes |
| R square | 0,7656 |

**Tukey's multiple comparisons test**

| | Mean 1 | Mean 2 | Mean Diff, | SE of diff, | n1 | n2 | q | DF | Significant | Summary | Adjusted P Value |
|---|---|---|---|---|---|---|---|---|---|---|---|
| VO vs. LO | 1,126 | 2,236 | -1,11 | 1,705 | 4 | 9 | 0,9207 | 19 | No | ns | 0,9643 |
| VO vs. MO | 1,126 | 3,472 | -2,347 | 2,167 | 4 | 3 | 1,532 | 19 | No | ns | 0,8129 |
| VO vs. AI | 1,126 | 12,68 | -11,55 | 1,831 | 4 | 6 | 8,922 | 19 | Yes | **** | < 0,0001 |
| VO vs. DLO | 1,126 | 2,722 | -1,597 | 2,457 | 4 | 2 | 0,9189 | 19 | No | ns | 0,9646 |
| LO vs. MO | 2,236 | 3,472 | -1,237 | 1,891 | 9 | 3 | 0,9248 | 19 | No | ns | 0,9638 |
| LO vs. AI | 2,236 | 12,68 | -10,44 | 1,495 | 9 | 6 | 9,877 | 19 | Yes | **** | < 0,0001 |
| LO vs. DLO | 2,236 | 2,722 | -0,4866 | 2,218 | 9 | 2 | 0,3103 | 19 | No | ns | 0,9994 |
| MO vs. AI | 3,472 | 12,68 | -9,207 | 2,006 | 3 | 6 | 6,49 | 19 | Yes | ** | 0,0017 |
| MO vs. DLO | 3,472 | 2,722 | 0,7503 | 2,59 | 3 | 2 | 0,4097 | 19 | No | ns | 0,9983 |
| AI vs. DLO | 12,68 | 2,722 | 9,957 | 2,317 | 6 | 2 | 6,079 | 19 | Yes | ** | 0,0032 |

**Table Analyzed** **Hypothalamus output**

**ANOVA summary**

| | |
|---|---|
| F | 0,4584 |
| P value | 0,7652 |
| P value summary | ns |
| Are differences among means statist | No |
| R square | 0,08802 |

**Tukey's multiple comparisons test**

| | Mean 1 | Mean 2 | Mean Diff, | SE of diff, | n1 | n2 | q | DF | Significant | Summary | Adjusted P Value |
|---|---|---|---|---|---|---|---|---|---|---|---|

| | Mean 1 | Mean 2 | Mean Diff, | SE of diff, | n1 | n2 | q | DF | Significant | Summary | Adjusted P Value |
|---|---|---|---|---|---|---|---|---|---|---|---|
| VO vs. LO | 6,8 | 6,252 | 0,5481 | 3,11 | 4 | 9 | 0,2492 | 19 | No | ns | 0,9998 |
| VO vs. MO | 6,8 | 3,058 | 3,742 | 3,953 | 4 | 3 | 1,339 | 19 | No | ns | 0,875 |
| VO vs. AI | 6,8 | 5,217 | 1,583 | 3,341 | 4 | 6 | 0,6703 | 19 | No | ns | 0,9888 |
| VO vs. DLO | 6,8 | 2,39 | 4,41 | 4,482 | 4 | 2 | 1,391 | 19 | No | ns | 0,8592 |
| LO vs. MO | 6,252 | 3,058 | 3,194 | 3,45 | 9 | 3 | 1,309 | 19 | No | ns | 0,8834 |
| LO vs. AI | 6,252 | 5,217 | 1,035 | 2,728 | 9 | 6 | 0,5368 | 19 | No | ns | 0,9952 |
| LO vs. DLO | 6,252 | 2,39 | 3,862 | 4,046 | 9 | 2 | 1,35 | 19 | No | ns | 0,8717 |
| MO vs. AI | 3,058 | 5,217 | -2,159 | 3,66 | 3 | 6 | 0,8343 | 19 | No | ns | 0,975 |
| MO vs. DLO | 3,058 | 2,39 | 0,6675 | 4,724 | 3 | 2 | 0,1998 | 19 | No | ns | > 0,9999 |
| AI vs. DLO | 5,217 | 2,39 | 2,826 | 4,226 | 6 | 2 | 0,9459 | 19 | No | ns | 0,9608 |

**Table Analyzed** **Thalamus output**

**ANOVA summary**

| | |
|---|---|
| F | 0,7145 |
| P value | 0,5923 |
| P value summary | ns |
| Are differences among means statist | No |
| R square | 0,1307 |

**Tukey's multiple comparisons test**

| | Mean 1 | Mean 2 | Mean Diff, | SE of diff, | n1 | n2 | q | DF | Significant | Summary | Adjusted P Value |
|---|---|---|---|---|---|---|---|---|---|---|---|
| VO vs. LO | 26,42 | 33,08 | -6,662 | 12,68 | 4 | 9 | 0,7432 | 19 | No | ns | 0,9836 |
| VO vs. MO | 26,42 | 20,59 | 5,823 | 16,11 | 4 | 3 | 0,5112 | 19 | No | ns | 0,996 |
| VO vs. AI | 26,42 | 14,85 | 11,57 | 13,62 | 4 | 6 | 1,201 | 19 | No | ns | 0,9115 |
| VO vs. DLO | 26,42 | 24,54 | 1,875 | 18,27 | 4 | 2 | 0,1451 | 19 | No | ns | > 0,9999 |
| LO vs. MO | 33,08 | 20,59 | 12,49 | 14,06 | 9 | 3 | 1,256 | 19 | No | ns | 0,8979 |
| LO vs. AI | 33,08 | 14,85 | 18,23 | 11,12 | 9 | 6 | 2,319 | 19 | No | ns | 0,4919 |
| LO vs. DLO | 33,08 | 24,54 | 8,536 | 16,49 | 9 | 2 | 0,7321 | 19 | No | ns | 0,9845 |
| MO vs. AI | 20,59 | 14,85 | 5,742 | 14,92 | 3 | 6 | 0,5445 | 19 | No | ns | 0,9949 |
| MO vs. DLO | 20,59 | 24,54 | -3,949 | 19,26 | 3 | 2 | 0,29 | 19 | No | ns | 0,9996 |
| AI vs. DLO | 14,85 | 24,54 | -9,691 | 17,22 | 6 | 2 | 0,7958 | 19 | No | ns | 0,9789 |

**Table Analyzed** **Brainstem output**

**ANOVA summary**

| | |
|---|---|
| F | 2,299 |
| P value | 0,0964 |
| P value summary | ns |
| Are differences among means statist | No |
| R square | 0,3261 |

**Tukey's multiple comparisons test**

| | Mean 1 | Mean 2 | Mean Diff, | SE of diff, | n1 | n2 | q | DF | Significant | Summary | Adjusted P Value |
|---|---|---|---|---|---|---|---|---|---|---|---|
| VO vs. LO | 12,27 | 17,84 | -5,573 | 5,608 | 4 | 9 | 1,405 | 19 | No | ns | 0,8549 |
| VO vs. MO | 12,27 | 3,091 | 9,176 | 7,127 | 4 | 3 | 1,821 | 19 | No | ns | 0,7017 |
| VO vs. AI | 12,27 | 5,431 | 6,836 | 6,024 | 4 | 6 | 1,605 | 19 | No | ns | 0,7864 |
| VO vs. DLO | 12,27 | 12,84 | -0,569 | 8,082 | 4 | 2 | 0,09957 | 19 | No | ns | > 0,9999 |
| LO vs. MO | 17,84 | 3,091 | 14,75 | 6,221 | 9 | 3 | 3,353 | 19 | No | ns | 0,1664 |
| LO vs. AI | 17,84 | 5,431 | 12,41 | 4,918 | 9 | 6 | 3,568 | 19 | No | ns | 0,127 |
| LO vs. DLO | 17,84 | 12,84 | 5,004 | 7,295 | 9 | 2 | 0,97 | 19 | No | ns | 0,9571 |
| MO vs. AI | 3,091 | 5,431 | -2,34 | 6,599 | 3 | 6 | 0,5016 | 19 | No | ns | 0,9963 |
| MO vs. DLO | 3,091 | 12,84 | -9,745 | 8,519 | 3 | 2 | 1,618 | 19 | No | ns | 0,7816 |
| AI vs. DLO | 5,431 | 12,84 | -7,405 | 7,619 | 6 | 2 | 1,374 | 19 | No | ns | 0,8644 |
